# Inference of cell state transitions and cell fate plasticity from single-cell with MARGARET

**DOI:** 10.1101/2021.10.22.465455

**Authors:** Kushagra Pandey, Hamim Zafar

## Abstract

Despite recent advances in inferring cellular dynamics using single-cell RNA-seq data, existing trajectory inference (TI) methods face difficulty in accurately reconstructing cell-state manifold and inferring trajectory and cell fate plasticity for complex topologies. We present MARGARET, a novel TI method that utilizes a deep unsupervised metric learning-based approach for inferring the cellular embeddings and employs a novel measure of connectivity between cell clusters and a graph-partitioning approach to reconstruct complex trajectory topologies. MARGARET utilizes the inferred trajectory for determining terminal states and inferring cell-fate plasticity using a scalable absorbing Markov Chain model. On a diverse simulated benchmark, MARGARET out-performed state-of-the-art methods in recovering global topology and cell pseudotime ordering. When applied to experimental datasets from hematopoiesis, embryogenesis, and colon differentiation, MARGARET reconstructed major lineages and associated gene expression trends, better characterized key branching events and transitional cell types, and identified novel cell types, and branching events that were previously uncharacterized.

## 1 Introduction

Dynamic cellular processes such as differentiation involve cell-state transitions that are characterized by cascades of epigenetic and transcriptional changes (1; 2). High-throughput single-cell RNA sequencing (scRNA-seq) datasets allow us to identify cellular identities at a single-cell resolution (3; 4) and thus can be utilized for elucidating the cellular heterogeneity of a dynamic cellular process and tracking cell fate decisions in normal as well as pathological development (5). Despite recent advances (6; 7) in inferring cellular dynamics from the underlying developmental process, existing computational trajectory inference (TI) methods (8; 9; 7; 10; 11) face several critical challenges. First, most TI methods have largely overlooked the importance of dimensionality reduction by focusing more on trajectory modelling and relying on generalized dimension reduction techniques such as UMAP (12), local linear embedding (11), or diffusion maps (9) which may obscure the identification of some intermediate cell states. Second, most TI methods impose strong assumptions on the topology of the trajectory and cannot generalize to disconnected or hybrid topologies without imposing further restrictions (13). Third, while Palantir introduced a Markov chain model to quantify the plasticity of cell fates along a trajectory, it predominantly assumes a connected trajectory and cannot generalize to disconnected trajectories. Lastly, accurate detection of terminal cell states remain difficult as only a few methods (e.g., Slingshot (10), Palantir (9), Monocle3 (7)) can automatically identify cell fates (Supplementary Table 1).

To overcome these challenges, we developed MARGARET (<u>M</u>etric le<u>AR</u>ned <u>G</u>raph p<u>AR</u>tition<u>E</u>d <u>T</u>rajectory), a graph-based TI method that employs an unsupervised metric learning-based approach for inferring the cell-state manifold where the distinct cell states are represented by compact cell clusters. To capture complex trajectory topologies, MARGARET employs the inferred cellular embeddings and the cell clusters to construct a cluster connectivity graph by using a novel measure of connectivity between cell clusters. The cluster connectivity graph is used in conjunction with the cell-nearest-neighbor graph to compute a pseudotime ordering of cells. To identify terminal states in the trajectory, MARGARET introduces a shortest-path betweenness-based measure. Using a novel algorithm that computes the probability for a cell to differentiate into each terminal state, MARGARET also quantifies the cell fate plasticity on the inferred trajectory by adopting and refining the absorbing Markov chain model of Palantir.

We validate the performance of MARGARET in trajectory inference and cell-fate prediction across a variety of synthetic and experimental scRNA-seq datasets. For simulated datasets (13) consisting of diverse topologies, MARGARET outperformed state-of-the-art TI methods both in terms of capturing the global topology and recovering the underlying pseudotime ordering. When applied to real biological datasets (9; 14; 15) representing human hematopoiesis, embryogenesis and colon differentiation, MARGARET accurately identified all major lineages along a pseudotemporal order that epitomized the expression trends of canonical cell-type markers in these processes. In hematopoiesis, MARGARET helped identify transitional progenitors associated with key branching events, which were also characterized by a drastic shift in MARGARET inferred cell-fate plasticity. For embryoid body differentiation, MARGARET helped identify and functionally characterize novel precursor populations. For colon differentiation, MARGARET characterized the lineage for BEST4/OTOP2 cells and the heterogeneity in goblet cell lineage in normal colon and inflamed ulcerative colitis.

## Results

### Overview of MARGARET

The overall framework of MARGARET consists of learning a lower-dimensional cell-state manifold from preprocessed scRNA-seq data (*N* cells) and inferring the underlying developmental trajectory by learning a cluster connectivity graph over the reduced-space cell-state embedding.

MARGARET uses an unsupervised metric learning-based approach (Fig. 1a, Methods) for learning a cell-state manifold where compact cell clusters represent the distinct cell states. Since the cluster assignment for the cells are unknown apriori, we compute an initial clustering of the data using an initial embedding (e.g., PCA). The initial embedding is episodically refined by a non-linear mapping network and minimizing the Triplet-Margin loss (Methods) for all the cells. The episodic training algorithm ultimately infers the lower-dimensional cell-state embedding 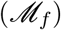 where cells are also partitioned into clusters. Given a refined partition of cells *y*\*, MARGARET employs a novel measure of connectivity (Methods) between cell clusters to construct an undirected graph 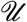 over *y*\* (Fig. 1b) to model the connected and disconnected regions in the manifold. Using 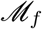, we first compute the k-nearest-neighbor (kNN) graph 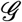, which is then used to estimate the connectivity between clusters given the partition *y*\*. The spurious edges in 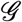 resulting from dropouts and artifacts in scRNA-seq data are pruned by using 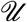 as a reference model (Fig. 1c, Methods). Given user-defined start cell(s), the pruned nearest-neighbor graph 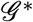 is then used to compute a robust estimate of pseudotime for each cell by computing the shortest path of each cell from the start cell(s). The final trajectory 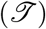 is then obtained by orienting the edges from the clusters with smaller average pseudotime to the ones with higher average pseudotime.

**Figure 1:**
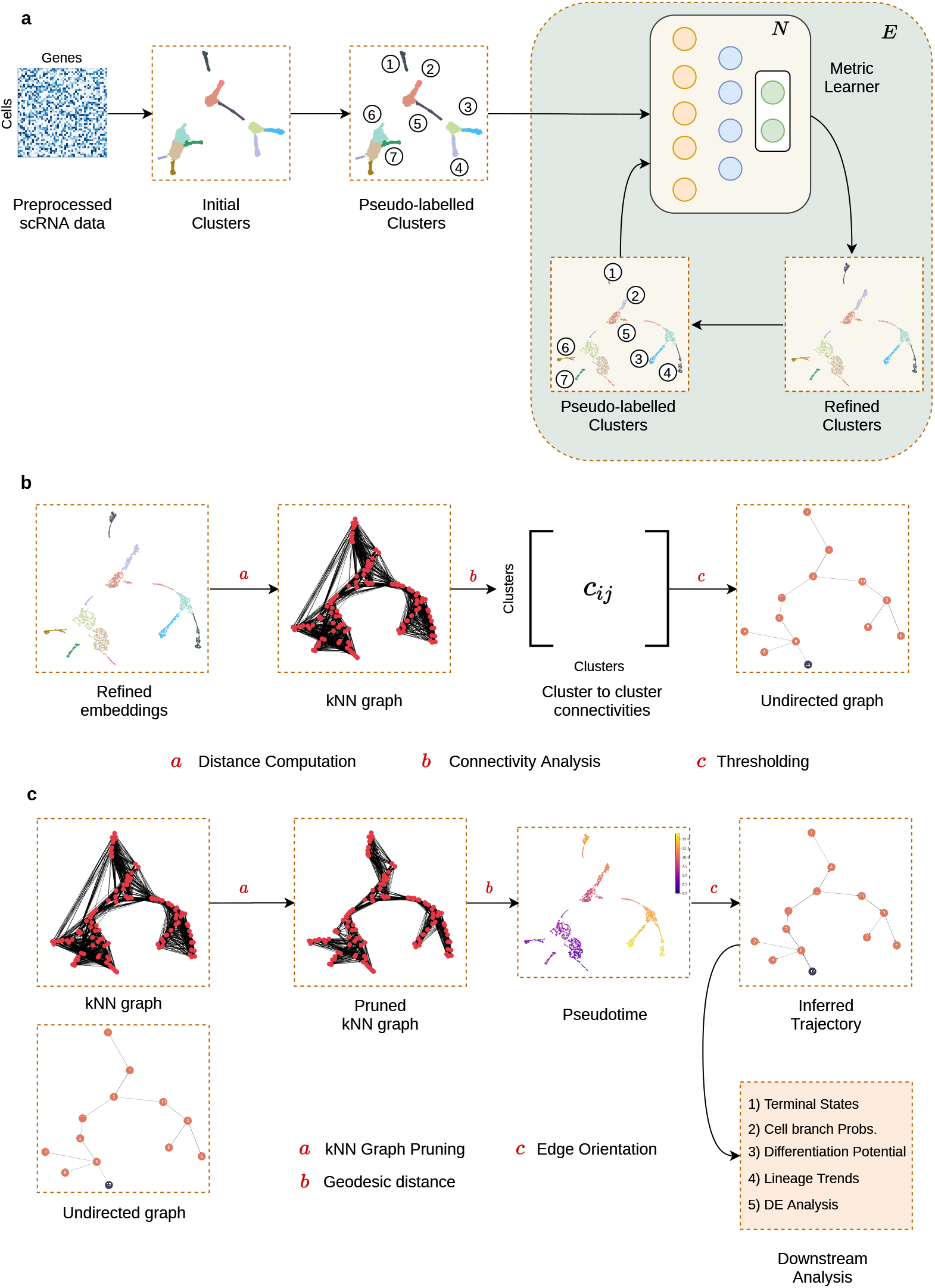
Overview of MARGARET. **(a)** Given a preprocessed scRNA-seq dataset, MARGARET uses an unsupervised metric-learning-based approach to learn compact cell-state representations from pseudolabels generated from an initial cell embedding. The cell cluster assignments and the MARGARET embedding are refined through an episodic training which results in the final refined embeddings and refined cell type clusters. **(b)** MARGARET infers the connectivity graph between the refined cell partitions by computing connectivities between clusters using a cell-level k-nearest-neighbor (kNN) graph. **(c)** MARGARET prunes the kNN graph by removing *short-circuit* edges and infers cell pseudotime from the pruned kNN graph by computing geodesic distances from the start cell(s). The pseudotime values are then utilized to infer a directed trajectory. MARGARET utilizes the inferred cell-state embeddings and the trajectory for several downstream tasks.

MARGARET then uses the refined cell embeddings, refined cell clusters and the cluster connectivity graph for downstream analysis including the prediction of terminal states, computation of cell branch probabilities, and differentiation potential (Supplementary Fig. 1, Methods). MARGARET employs a shortest-path betweenness-based measure (Methods) for identifying nodes in 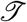 as terminal states (Supplementary Fig. 1a, Methods). Finally, MARGARET also encompasses a novel algorithm for calculating the cellular differentiation potential (DP) that adopts the absorbing Markov chain model of cellular differentiation from Palantir (9) and generalizes it for disconnected trajectories.

### MARGARET outperforms other TI methods on a diverse simulated benchmark

We benchmarked MARGARET’s performance on a variety of synthetic datasets (Methods) consisting of multifurcating and disconnected trajectories (with multifurcating components)(Supplementary Table 3) against state-of-the-art TI methods Partition-Based Graph Abstraction (PAGA) (16), Palantir (9), and Monocle3 (7).

A qualitative comparison (Fig. 2a-b) of the trajectories inferred by the algorithms showed that for the disconnected dataset, Palantir failed to capture the disconnected components (Fig. 2b), PAGA and Monocle3 captured the disconnected topology but underestimated the number of clusters in each disconnected component. For the multifurcating dataset, PAGA and Palantir failed to capture the correct branching topology, whereas Monocle3 failed to recover the global topology as it inferred the overall trajectory as two separate components (Fig. 2b). In contrast, MARGARET outperformed other methods on these datasets by accurately recovering the underlying global topology without over/under clustering in the embedding space.

**Figure 2:**
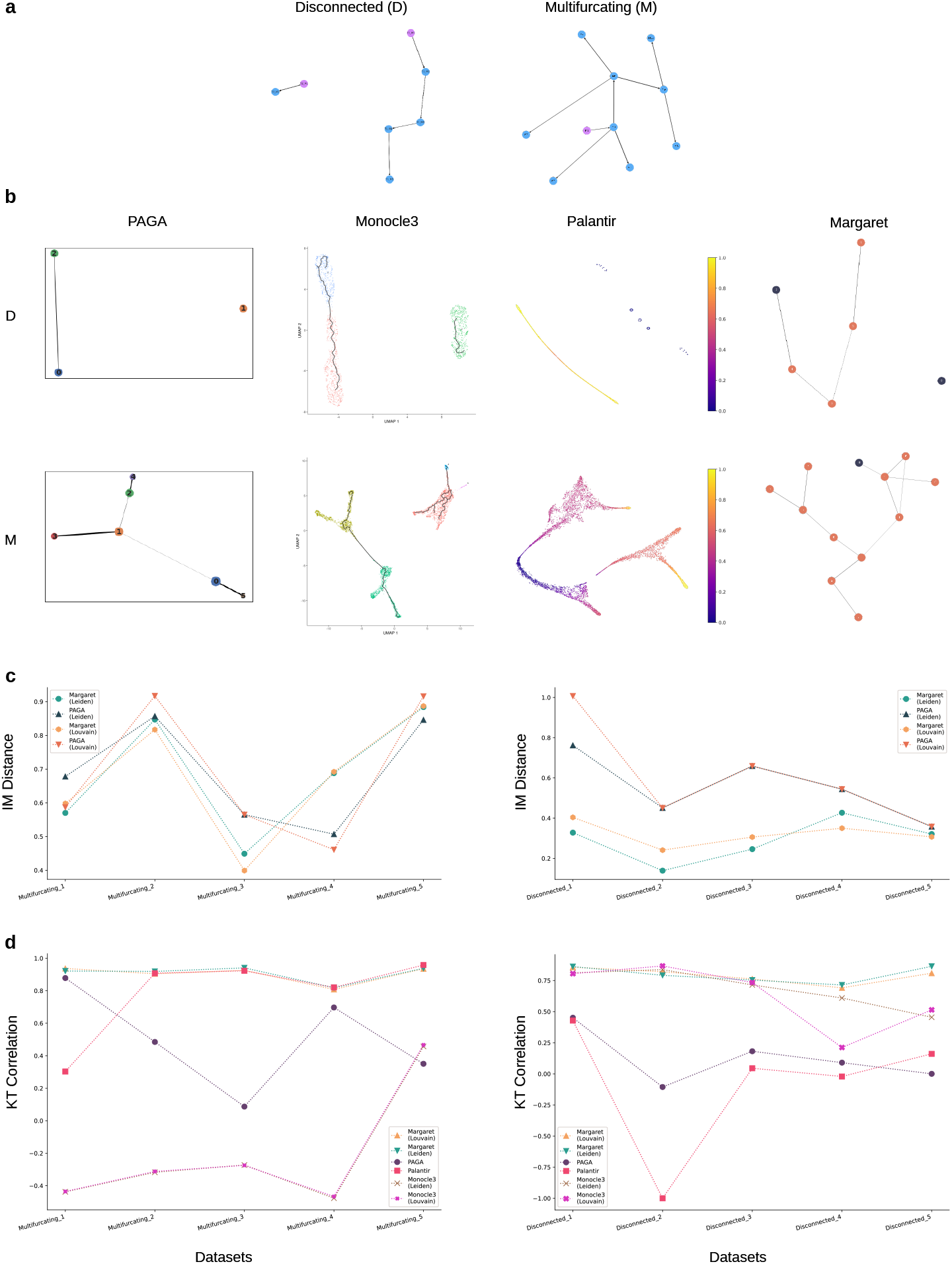
MARGARET outperforms state-of-the-art TI methods. MARGARET out-performed state-of-the-art TI methods on qualitative and quantitative metrics when applied to a simulated benchmark consisting of ten datasets with diverse trajectory types. **(a)** (Left) Sample disconnected dataset (1966 cells) (Right) Sample multifurcating dataset (5000 cells) **(b)** Visualization of the cell embedding landscape inferred by different TI methods on the disconnected (Top) and multifurcating (Bottom) datasets. Palantir embedding landscapes shown with projected pseudotime values. Monocle3 inferred trajectory graph projected on the embedding with colors denoting cluster information. PAGA and MARGARET outputs shown as connectivity graphs. PAGA, Monocle3 and MARGARET inferred graphs were computed using Louvain clustering with 0.4 resolution. **(c)** IM distance comparison between MARGARET and PAGA on the multifurcating (Left) and the disconnected (Right) benchmarks (lower score is better).**(d)** Kendall’s Tau (KT) correlation comparison between MARGARET and other TI methods on the multifurcating (Left) and the disconnected (Right) benchmarks (higher score is better). Refer to Supplementary Table 2 for exact KT scores and SR scores.

For quantitative benchmarking, first we assessed a TI method’s ability in recovering the underlying global topology of cells by evaluating the Ipsen-Mikhailov (IM) distance (Methods) between the ground truth and the inferred trajectory graph. Two trajectories (represented as graphs) were compared using the ‘network of milestones’ representation framework (13). Since among the state-of-the-art methods, only PAGA-inferred trajectories can be represented using this framework, we compared the accuracy of global topology inference only against PAGA. On this task, MARGARET outperformed PAGA on all the disconnected-trajectory datasets and most of the multifurcating-trajectory benchmarks (Fig. 2c). Overall, MARGARET exhibited superior performance on the global topology prediction task compared to PAGA across a range of clustering methods and resolutions (Supplementary Fig. 2a-b).

Next, we evaluated the TI methods’ accuracy in recovering the pseudotime ordering of cells by computing the Kendall’s Tau (KT) and the Spearman Rank (SR) correlation (Methods) between the ground truth and the inferred ordering. On the multifurcating benchmark, PAGA and Monocle3 failed to capture the underlying ordering of cells, with Monocle3 consistently showing negative KT correlation across most of the datasets (Fig. 2d). Palantir performed better than PAGA and Monocle3 on the multifurcating benchmark. However, Palantir and PAGA performed much worse on the disconnected-trajectory benchmark, with Monocle3 performing comparably better. Across both the benchmarks, MARGARET outperformed other methods on most of the datasets across both Louvain and Leiden clustering (Fig. 2d; Supplementary Fig. 3; Supplementary Table 2). Even across different clustering resolutions, MARGARET consistently exhibited high KT and SR scores on the simulated benchmark (Supplementary Fig. 2c-d).

MARGARET was also able to capture the global cyclic topology for a synthetic cyclic trajectory (Supplementary Note 1, Supplementary Fig. 4). Finally, the DP inference by MARGARET was superior to that of Palantir for disconnected trajectories indicating the applicability of MARGARET’s DP inference to a wider variety of trajectories (Supplementary Note 1, Supplementary Fig. 5).

### MARGARET learns modular clusters and refines other single-cell embeddings

For two scRNA-seq human hematopoiesis datasets (9), MARGARET consistently learned cluster representations whose modularity score improved in each episode across Louvain and Leiden clustering backends (Supplementary Note 2, Supplementary Fig. 6). By applying MARGARET on four biological datasets with available ground-truth clustering annotations (Methods), we found that MARGARET was able to improve the clustering quality (as measured by adjusted rand index (ARI) and normalized mutual information (NMI) metrics) of the single-cell embeddings learned by PCA and scVI (17) (Supplementary Note 2, Supplementary Fig. 7).

### MARGARET correctly predicts human hematopoietic differentiation trajectory and associated gene expression changes

Due to the availability of established lineage-specific markers, hematopoiesis (18) has been used as a model biological system by several trajectory inference methods (19; 20; 9). Using MARGARET, we first explored early human hematopoiesis where hematopoietic stem cells (HSCs), through a hierarchy of progenitors and bifurcation events, give rise to different mature cell types (18). We applied MARGARET to two human bone marrow scRNA-seq datasets (10X Chromium) (9) (replicates 1 and 2 consisting of 5780 and 6501 cells respectively). For both datasets, MARGARET correctly identified all major hematopoietic cell types, including hematopoietic stem cells (HSCs), common lymphoid and myeloid progenitors (CLPs and CMPs respectively), as well as cells committed towards erythroid (erythrocytes and megakaryocytes), monocytic and dendritic cell (classical and plasmacytoid dendritic cells (cDCs and pDCs)) lineages (Fig. 3a, Supplementary Fig. 8a). The cell type clusters inferred by MARGARET were characterized by the expression of key marker genes, obtained by manually curating a set of marker genes for major hematopoietic cell types through prior literature review (9; 21; 22; 23) (Fig. 3d, Supplementary Fig. 8f, 9). The expression of the marker genes corresponding to major hematopoietic cell types correlated well with the topology of the MARGARET inferred trajectory (Supplementary Fig. 10). We utilized the starting cell information provided by (9) for pseudotime inference (Fig. 3b, Supplementary Fig. 8c). For both replicates, MARGARET inferred pseudotime follows expected progression, where the pseudotime increases as cells progress towards more specialized cell types from *CD34* enriched stem cells. Moreover, the probability of cells branching to different lineages diminishes as cells commit towards specific lineages (Supplementary Figs. 11-13). Consequently, as expected, MARGARET inferred DP (Fig. 3c, Supplementary Fig. 8b) decreases as we move towards terminal states in the trajectory since commitment towards a specific lineage is accompanied by a gradual reduction in cell plasticity. Fig. 3e and Supplementary Fig. 8d represent the annotated hematopoietic trajectory inferred by MARGARET for the two replicates, where the arrows represent transition between cell types.

**Figure 3:**
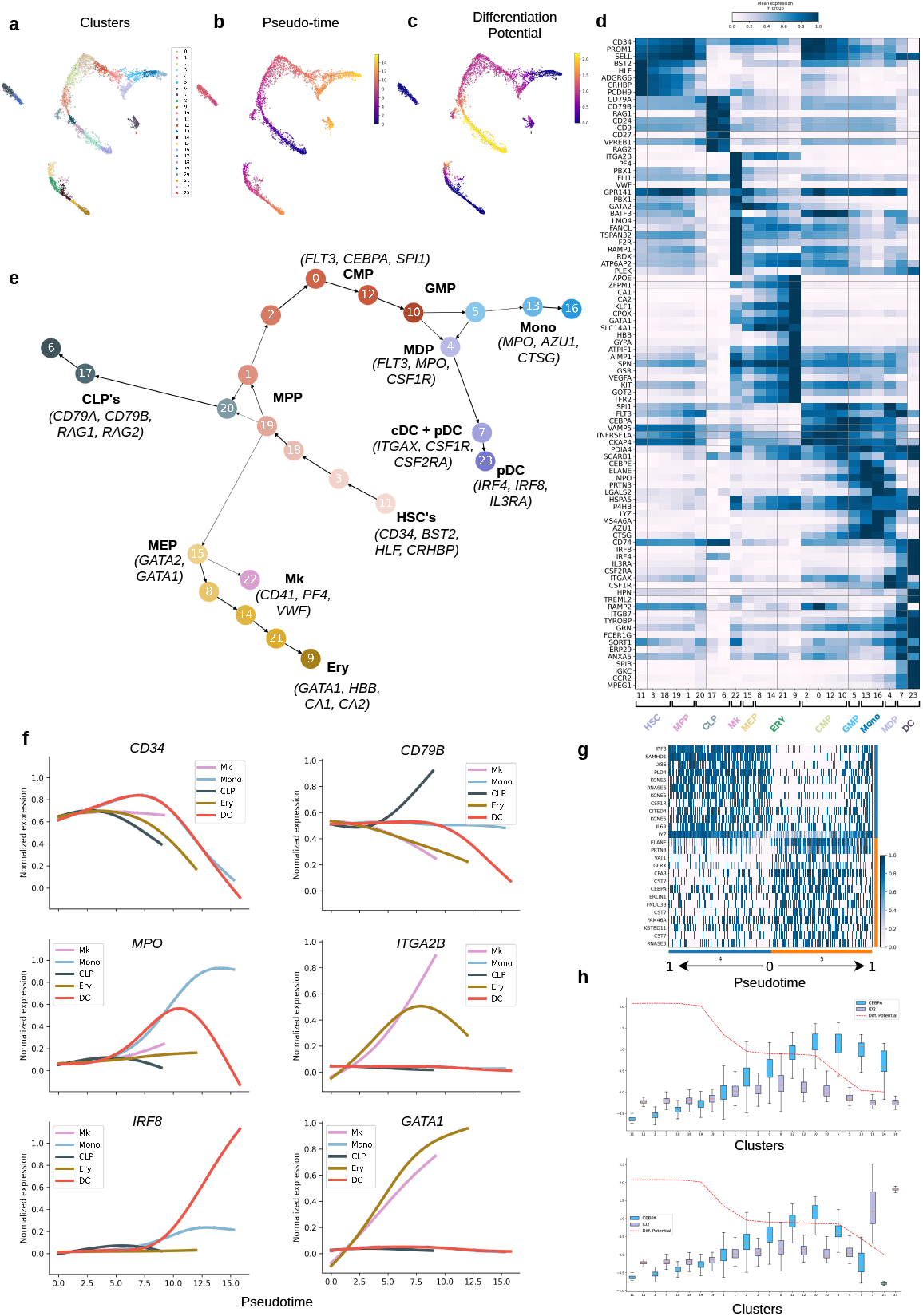
MARGARET delineates major lineages and identifies important transcriptional switches in early human hematopoiesis. Analysis of scRNA-seq data for human hematopoiesis replicate 1 by MARGARET. **(a)** tSNE plot of cell-state embedding inferred by MARGARET for the human hematopoiesis dataset, cells are colored by MARGARET inferred clusters. **(b)** MARGARET pseudotime and **(c)** differentiation potential calculated using one HSC as a start cell. **(d)** Heat map for marker genes for all MARGARET inferred clusters. **(e)** MARGARET inferred trajectory annotated with cell-type and lineage information (important marker genes are mentioned within parantheses with the cell type annotation). Ery: Erythrocyte; Mk: Megakary-ocyte; MEP: Megakaryocyte-Erythroid Progenitors; HSC: Hematopoeitic Stem Cells; CLP: Common Lymphoid Progenitors; CMP: Common Myeloid Progenitors; GMP: Granulocyte-Monocyte Progenitors; MDP: Monocyte-Dendritic Cell Progenitors; cDC: Classical Dendritic Cells; pDC: Plasmacytoid Dendritic Cells; Mono: Monocytes. **(f)** Gene expression trends for essential genes for major inferred lineages. **(g)** Differential expression (DE) analysis between cluster 4 (MDP) and cluster 5 (GMP) **(h)** Variation of *CEBPA* and *ID2* gene expressions in the monocyte (Mono) lineage (top) and the dendritic cell (DC) lineage (bottom) for replicate 1. The boxplots summarize the expression of the gene in each cluster in the lineage, where the box depicts the interquartile range (IQR, the range between the 25th and 75th percentile) with the median value, whiskers indicate the maximum and minimum value within 1.5 times the IQR. The red dotted line represents the mean differentiation potential for each cluster in the lineage.

To validate MARGARET inferred trajectories, we computed expression trends for essential marker genes for all major hematopoietic lineages (Fig. 3f). As expected, expression of *CD34* decreases with increasing pseudotime as cells commit to particular lineages (18). In contrast, *CD79B* is selectively upregulated in the lymphoid lineage (24) while *MPO* and *IRF8* are upregulated in the monocyte (25) and dendritic cell (DC) (26) lineages, respectively. *ITGA2B* and *GATA1* are selectively upregulated in the megakaryocytic (27) and erythroid (28) lineages, respectively. Similar expression trends were observed for replicate 2 demonstrating MARGARET’s robustness (Supplementary Fig. 8e).

### MARGARET characterized progenitor populations for monocytic and dendritic cell lineages

Interestingly, we observed an initial upregulation in *MPO* expression in both the monocyte and DC lineages (Fig. 3f). However, with pseudotime progression, *MPO* expression is upregulated in the monocyte lineage but gets downregulated in the DC lineage. To explore the branching of monocyte and DC lineages from CMPs in replicate 1, we investigated the transitions from cluster 10 to cluster 4 and cluster 10 to cluster 5 that are also associated with substantial changes in DP indicating that these transitions accompany important molecular events corresponding to lineage commitment (9). At the transition from CMPs to monocytes, we observed elevated expressions of monocyte markers ((21)) including *CEBPA*, and *CEBPE* (Fig. 3d,h (top), Fig. 4a), which correlated with a decrease in DP. The transcription factor (TF) *CEBPA* plays a crucial role in cell fate decisions in granulocyte-monocyte progenitors (GMPs) (29; 30) to differentiate into granulocyte and monocyte (21). Apart from elevated expression of *CEBPE* and monocyte markers *MPO*, *LYZ*, and *MS4A6A* (22), the monocyte clusters also strongly expressed granule genes such as *CTSG*, *PRTN3* and *ELANE* (Fig. 3d), each of which were also highly correlated (> 0.92) with the MARGARET inferred monocytic branch probabilities (Fig. 4c). Monocytes derived from granulocyte-monocyte progenitors (GMPs) are known to express these granule proteases (31). Therefore, expression of *CEBPA*, *CEBPE*, and granule proteases and reduced *FLT3* expression in cluster 5 (Fig. 4a (left)) indicates the presence of GMPs in cluster 5 (31). We also observed similar trends for replicate 2 (Supplementary Fig. 14).

**Figure 4:**
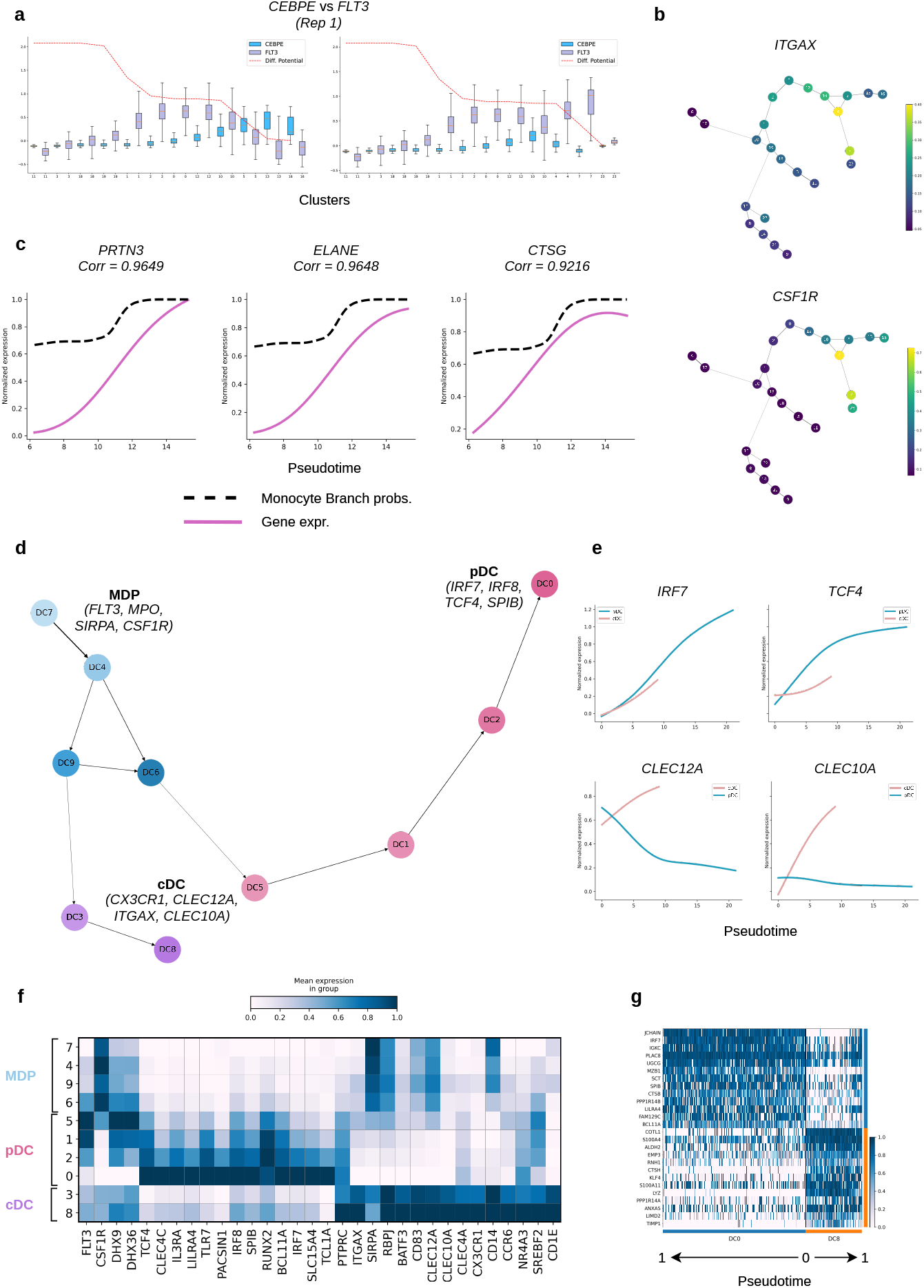
MARGARET characterizes cellular heterogeneity in the monocytic and dendritic cell lineages. **(a)** Variation of the expression of *CEBPE* and *FLT3* across the monocyte (Mono) lineage (left) and the DC lineage (right) for replicate 1. The boxplots summarize the expression of the gene in each cluster in the lineage, where the box depicts the interquartile range (IQR, the range between the 25th and 75th percentile) with the median value, whiskers indicate the maximum and minimum value within 1.5 times the IQR. The red dotted line represents the mean differentiation potential for each cluster in the lineage. **(b)** Mean expression of *ITGAX* (top) and *CSF1R* (bottom), projected on the MARGARET inferred connectivity graph. **(c)** Gene expression trends for neutrophil-like monocyte marker genes: *PRTN3*, *ELANE*, *CTSG* are highly correlated with monocyte branch probability (dotted black line). **(d)** MARGARET inferred trajectory for the DC sub-lineage obtained from the combined analysis of replicates 1 and 2. MARGARET accurately recovers the cDC and pDC lineages. **(e)** Gene expression trends for pDC markers (top): *IRF7*, *TCF4* and cDC markers (bottom): *CLEC12A*, *CLEC10A*. **(f)** Heat map of DC-lineage specific marker genes for MARGARET inferred DC sub-lineage clusters. **(g)** Comparison of differentially Expressed genes between cDCs (cluster DC8) and pDCs (cluster DC0).

In contrast, we observed elevated expressions of *FLT3* in cluster 4 (Fig. 4a (right)) which also correlated with a decrease in DP on the transition from cluster 10 (*FLT*3^+^ CMP) to cluster 4. Moreover, this cluster showed elevated expression of *CSF1R* (*CD115*), dendritic cell marker *ITGAX* (*CD11c*) (Fig. 4b), and *MPO* (Fig. 3d). Altogether, the *FLT* 3^+^*CD*115^*hi*^ signature of this cluster suggests the presence of monocyte-dendritic dell progenitors (MDPs) in cluster 4 which gives rise to both monocytes and dendritic cells (31). Therefore, MARGARET inferred branching structure in the myeloid lineage characterized the differentiation of monocytes and dendritic cells from GMPs and MDPs respectively and these important events were also marked by a decrease in MARGARET inferred DP. These progenitor populations and their lineage branchings were not characterized in the original study (9) that used Palantir.

We further characterized the heterogeneity in the DC lineage using MARGARET where it identified the cDCs and pDCs as the terminal states (Fig. 4d-g, Supplementary Note 3). Furthermore, the cDC population was found to consist of mostly type 2 conventional dendritic cells (cDC2) as marked by elevated expression of TFs *NR4A3, SREBF2* and marker genes *CLEC10A, CD1E, CLEC12A, CX3CR1* (32). The DP change also characterized the branching point in the erythroid lineage where we identified the presence of megakaryocyte-erythroid progenitors (MEPs) in cluster 15 (Supplementary Note 3, Supplementary Figs. 15-16).

### MARGARET accurately uncovers the differentiation landscape in embryoid bodies

To investigate MARGARET’s ability in extracting novel insights from a complex biological system, we applied MARGARET to an scRNA-seq dataset generated from embryoid bodies (EB) (14), which recapitulates the differentiation process in early embryogenesis where pluripotent embryonic stem cells (ESCs) give rise to early lineage precursors. The dataset consisted of 16821 cells, sampled at 3-d intervals over a 27-d differentiation time course (14). Even though the sampling time information was not utilized to learn the cell-state manifold, MARGARET inferred embedding consisting of 26 clusters (Fig. 5a) retained the time trend accurately (Fig. 5b), thus preserving the global topology of the data. Based on expression of marker genes (14) (Fig. 5d, Supplementary Fig. 17a) and gene ontology (GO) analysis (Methods, Supplementary Table 4), we identified endoderm (EN), mesoderm (ME), neural crest (NC), neuroectoderm (NE), and neuronal subtypes (NS) (including neural progenitors (NP)) lineages along with ESCs in the data (Supplementary Fig. 18). Moreover, MARGARET inferred pseudotime (Fig. 5c) and DP followed the progression of cell types where decrease in DP was concordant with the major lineage commitments in EB differentiation (Supplementary Note 4, Supplementary Figs. 19-21). The lineage specification map of embryoid bodies inferred by MARGARET (Fig. 5e) was characterized by the expression trends of key marker genes for the terminal cell types (Fig. 5f, Supplementary Note 4).

**Figure 5:**
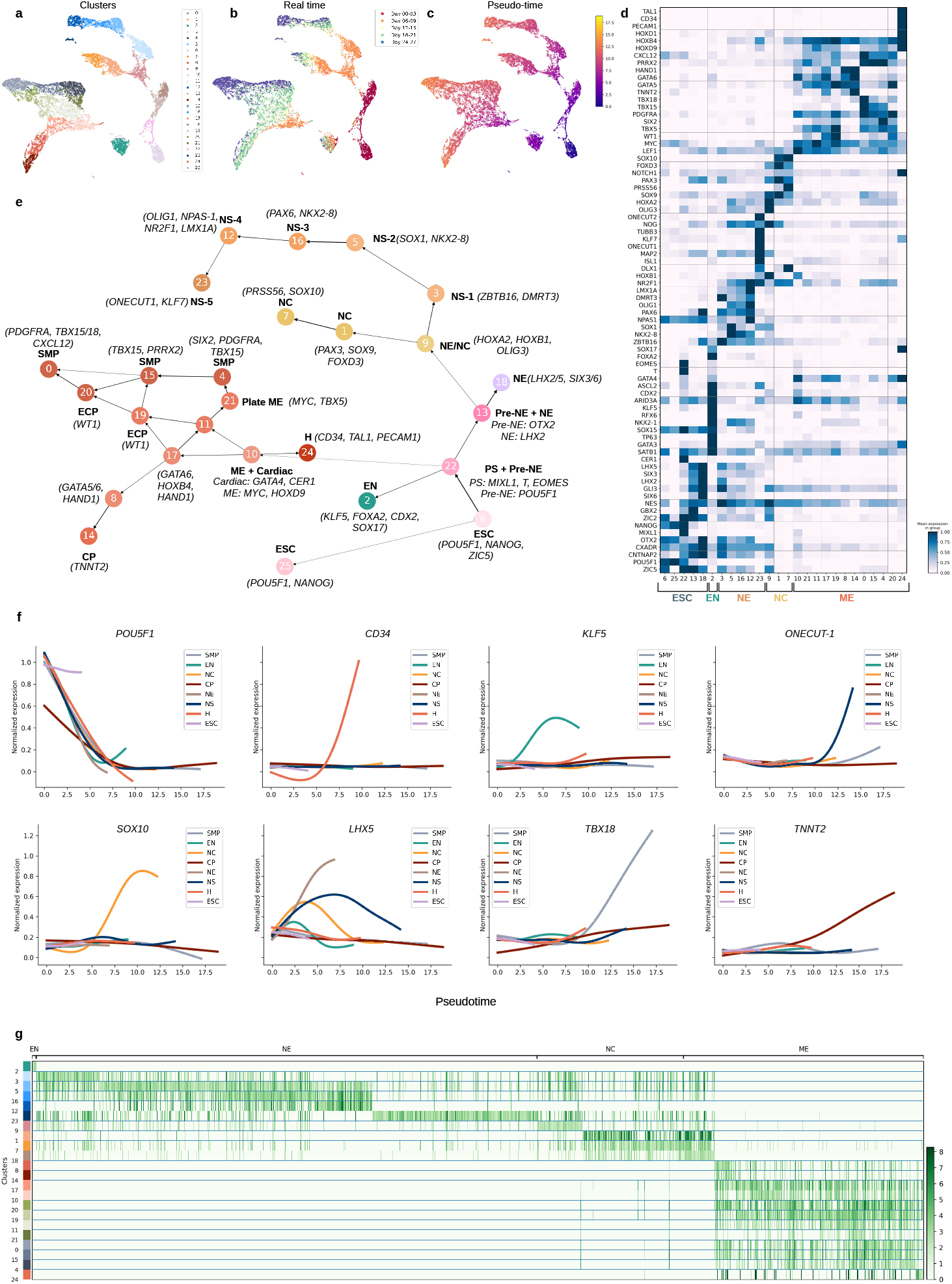
MARGARET characterizes the differentiation trajectory in human embryoid bodies. MARGARET reconstructs a detailed lineage map and identifies novel cell types and differentiation intermediates in the embryoid body (EB) dataset (*n* =). **(a)** UMAP plot of cell-state embedding inferred by MARGARET for EB dataset. **(b)** MARGARET inferred cell embedding preserves real time information. **(c)** MARGARET pseudotime projected on the cell embeddings. **(d)** Heat map of marker genes for all MARGARET inferred clusters. **(e)** MARGARET inferred trajectory annotated with cell-type and lineage information (characteristic marker genes are mentioned within parantheses with the cell-type annotation). ESC: Embryoid stem cells; PS: Primitive streak; NE: Neuroectoderm; NC: Neural crest; NS: Neuronal subtypes; EN: Endoderm; H: Hemangioblasts; ME: Mesoderm; SMP: Smooth muscle precursors; ECP: Epicardial precursors; CP: Cardiac precursors. **(f)** Gene expression trends for the inferred lineages. **(g)** Gene ontology (GO) analysis of MARGARET inferred clusters (grouped by major lineages). The heatmap value for a GO term was set to 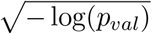, where *p*_*val*_ is the p-value for the corresponding GO term.

In the ectodermal lineage, similar to (14), MARGARET was able to identify the bipotent precursors (cluster 9 expressing *HOXA2, HOXB1* and *OLIG3* (Fig. 5d)) that originated from the neuroectoderm cells expressing *GBX2* and bifurcated from cluster 9 into the neural crest and neuronal sub-lineages (Fig. 5e). Further characterization of this branching revealed correlation between a decrease in MARGARET inferred DP and the up/down regulation of important TFs in the neural crest and neuronal lineages (Supplementary Fig. 22). The DP drop in the neural crest lineage was concordant with the upregulation of canonical TFs *SOX9/10* (33; 34) (Fig. 5d,f, Supplementary Fig. 22a,b (right)) while neuronal-subtype cluster 3 exhibited upregulation of TFs *SOX1* and *LHX2* (Fig. 5d, Supplementary Fig. 22a,b (left)), which have been shown to be important for subtype specification in certain types of neurons (35; 36). GO analysis at finer resolution (Fig. 5g, Supplementary Table 5) also revealed the enrichment of both neural crest and neuronal differentiation-related functions in cluster 9 further validating its bi-potency. We further validated the neural crest sub-branch detected by MARGARET using the bulk RNA-seq data provided by (14) for FACS purified *CD*49*d*^+^*CD*63^−^ cells (Supplementary Fig. 23a, Supplementary Note 4).

In the mesoderm lineage, MARGARET trajectory (Fig. 5e) identified a series of intermediate precursors expressing (*CER1, GATA1*), (*GATA6, HOXB4*) and (*GATA5/6, HAND1*), respectively which finally gave rise to cardiac precursors (expressing *TNNT2*). Bulk-RNA analysis for FACS purified *CD*82^+^*CD*142^+^ cells revealed the highest correlation of the single-cell expression profiles in the mesoderm sub-lineage harboring cardiac precursors (cells with high *CD82* and *CD142* expressions) (Supplementary Fig. 23b), suggesting that MARGARET accurately detects this sub-lineage. Moreover, we found three types of smooth muscle precursors (SMPs) which expressed different markers (*SIX2*, *PRRX2*, and *TBX15/18*) suggesting that SMP differentiation in the mesoderm lineage proceeds via one or more differentiation intermediates from plate ME cells. Thus, MARGARET accurately resolved the underlying trajectory structure in the EB dataset by inferring branchings and intermediate cell populations essential for cell lineage commitment in early embryogenesis.

### MARGARET elucidates colon differentiation

To investigate colonic epithelial differentiation, we applied MARGARET to a scRNA-seq dataset comprising three conditions - healthy (4249 cells), clinically inflamed ulcerative colitis (UC) (2848 cells), and non-inflamed UC (4078 cells) across three replicates (15). For the healthy colon (Fig. 6), MARGARET accurately localized the absorptive and secretory cell lineages from the data (Fig. 6a,d,f) while also identifying absorptive progenitors, colonocytes, crypt-top (CT) colonocytes, and BEST4/OTOP2 cells in the absorptive lineage; and secretory progenitors, goblet cells, and enteroendocrine cells (EECs) in the secretory lineage (Fig. 6e, more details in Supplementary Note 5). MARGARET inferred branch probabilities (Supplementary Fig. 24) and gene expression trends (Fig. 6h) further validated the cell populations.

**Figure 6:**
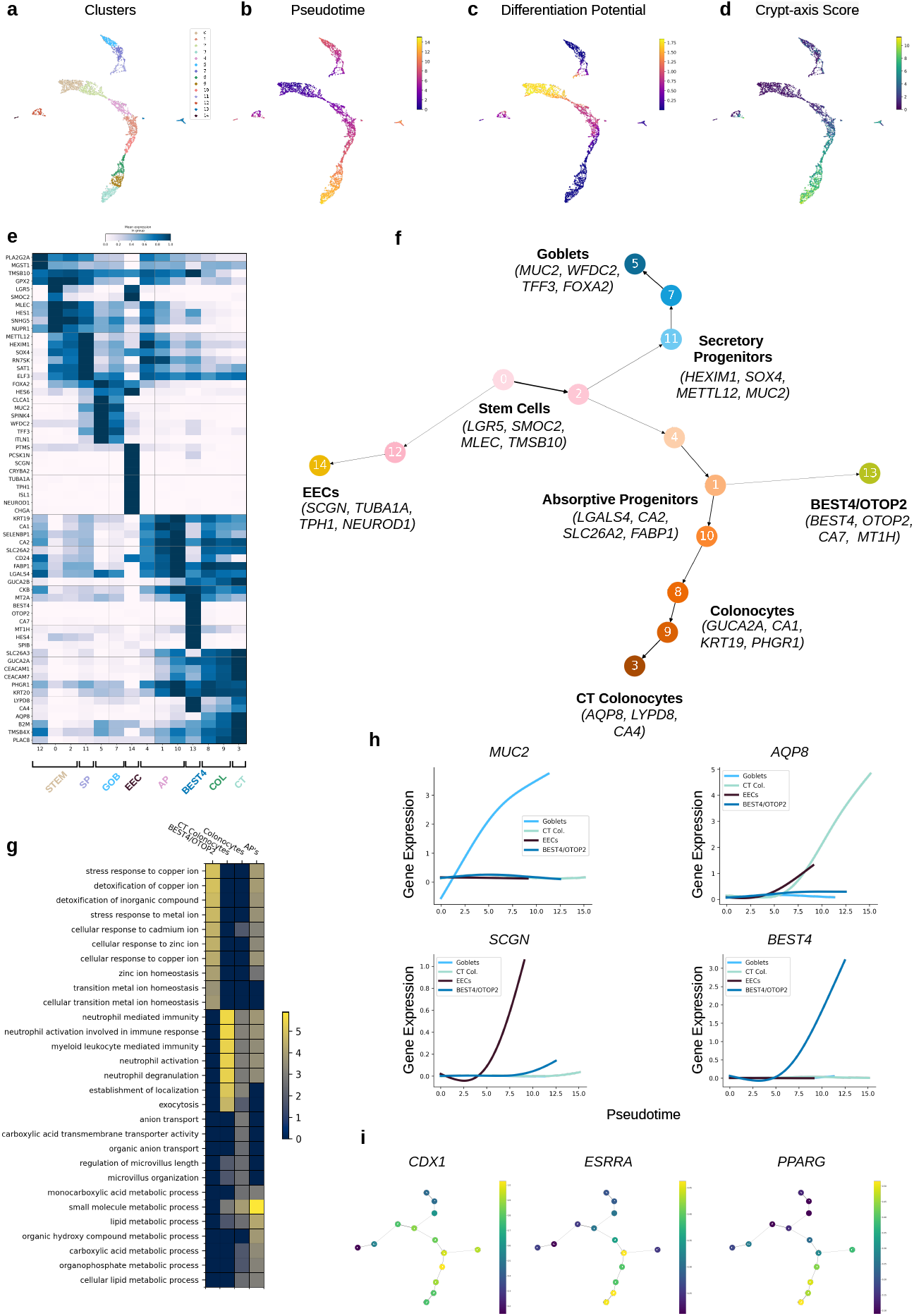
MARGARET delineates the differentiation trajectory for colonic epithelial cells for healthy humans. **(a)** UMAP plot of cell-state embedding inferred by MARGARET for healthy human colonic epithelium (*n* = 3), cells are colored by clusters inferred by MARGARET. **(b)** MARGARET pseudotime and **(c)** differentiation potential for each cell projected on the cell embeddings. **(d)** Crypt-axis score (See Methods) projected on MARGARET cell embeddings. **(e)** Heat map showing marker genes for each MARGARET inferred cluster. Marker genes were curated from (15). **(f)** MARGARET inferred trajectory annotated with cell type and lineage information (important marker genes for each cell type are mentioned within parantheses with the cell type information). CT: Crypt-top; EEC: enteroendocrine cells **(g)** Gene Ontology (GO) analysis for major inferred cell types in the absorptive cell lineage. Top GO:BP terms were included for each cell type. The value for a GO term in the heat map was set to 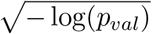, where *p*_*val*_ is the *p*-value for the corresponding GO term. **(h)** Gene expression trends for essential genes for major inferred lineages. **(i)** Mean expression of TFs *CDX1*, *ESRRA* and *PPARG*, projected on the MARGARET inferred connectivity graph.

In the absorptive cell lineage, BEST4/OTOP2 cells were detected as a terminal state that branched from the absorptive progenitors (APs) before they gave rise to colonocytes. GO analysis of absorptive lineage clusters revealed the role of BEST4/OTOP2 cells in maintaining metalion transport and homeostasis in the colonic epithelium (Fig. 6g). Notably, GO terms for the BEST4/OTOP2 cell cluster did not include any overlapping terms with CT colonocytes and colonocytes, suggesting the functional variability between these cell types. However, BEST4/OTOP2 cells exhibited high CA scores (Fig. 6d) and expressed colonocyte-specific markers *GUCA2A*, *CEACAM1*, and *CEACAM7*, which suggests their similarity to mature colonocytes and localization towards the crypt top. Moreover, APs showed enrichment of GO terms corresponding to both BEST4 cells and colonocytes (Fig. 6g). The branching of BEST4/OTOP2 cells was also marked by the downregulation of the TF *ESRRA* and the high expression of TFs *CDX1* and *PPARG* (Fig. 6i). We observed a similar branching in the absorptive lineage under non-inflamed UC (Supplementary Fig. 25) with similar GO enrichment for major absorptive cell types. Therefore, our findings suggest that BEST4/OTOP2 cells are mature colonocytes with a different functional profile as compared to CT colonocytes and originate from APs as a different sub-lineage within the absorptive lineage.

Given the crucial role of goblet cells in colonic barrier maintenance (37), we further characterized the transcriptional landscape of the goblet cell lineages under healthy and inflamed UC conditions using MARGARET (Supplementary Fig. 26). Under both conditions, MARGARET reconstructed a linear trajectory in the goblet cell lineage from immature goblet cells to mature goblet cells (Supplementary Fig. 26g,h) (maturity of cells indicated by pseudotime order (Supplementary Fig. 26c,f)). We observed relatively higher expression levels of *WFDC2* in immature goblet cells than in mature goblet cells under both healthy and inflamed conditions. In contrast, *MUC2* was more expressed in mature goblet cells (Supplementary Fig. 26i). Moreover, mature goblet cells had higher CA scores than immature goblet cells, suggesting that these cells reside towards crypt top (Supplementary Fig 26b,e). The functional characterization (Supplementary Fig. 26j) of the goblet cell clusters revealed the enrichment of GO terms related to wound healing, immune, and stress response in the mature goblet cells under inflamed UC condition indicating their potential role in inflammatory responses to inflammatory bowel disease (IBD). The goblet cells in inflamed UC further showed spatial and crypt-wide transcriptional heterogeneity. The mature goblets in inflamed UC showed elevated expressions of *SPINK1* and *SPINK4* (Supplementary Fig. 27a), genes that are normally expressed by immature goblets in healthy colon. Moreover, we also observed transcriptional dysregulation of interferon-regulated cytokines including *CD164*, *CD55*, and *IRF7* (38) throughout the goblet cell lineage in inflamed UC (Supplementary Fig. 27b).

## Discussion

As single-cell datasets grow in size and complexity, MARGARET addresses the need for the accurate detection of cell-state lineages, prediction of cell fates, and the inference of cell plasticity in complex topologies underlying dynamic cellular processes. The end-to-end computational framework of MARGARET alleviates the challenges faced by existing TI methods: inability of the classical dimensionality reduction methods to accurately recover the underlying topology, insufficient generalizability to connected and disconnected graph trajectories beyond tree-structured topologies, accurate detection of less-sampled cell fates, and inference of cell fate plasticity for complex trajectory types. Our analysis of a diverse simulated benchmark consisting of challenging topologies showed that MARGARET generalizes to complex trajectories and can accurately infer the underlying topology and pseudotime order of cells in the trajectory while outperforming state-of-the-art methods on the same. Moreover, using synthetic disconnected trajectories, we showed that MARGARET can accurately infer DP for each component in the disconnected trajectories whereas Palantir’s DP inference is mostly incorrect for such datasets. (Supplementary Fig. 5).

Using multiple biological datasets, we also showed that MARGARET’s metric learning-based approach can *refine* the cellular latent space inferred by other dimensionality reduction methods. On a variety of important biological systems, we showed that MARGARET identified all the major lineages and the established cell fates and recovered the marker gene expression trends. In early human hematopoiesis, MARGARET identified progenitor populations associated with the branching points (MDP and GMP in the myeloid and MEP in the erythroid lineages respectively) which were not characterized in the original study that used Palantir. MARGARET also identified cDC2 as a terminal state in the dendritic cell lineage along with pDCs. For embryoid body differentiation, MARGARET reconstructed a detailed lineage map of all major and sub-lineages and further validated the presence of novel differentiation intermediates in the neuroectoderm and mesoderm lineages and identified novel smooth muscle precursor populations. For colonic epithelial differentiation, MARGARET helped identify a branch point for BEST4/OTOP2 cells in the absorptive lineage, while also uncovering the dysregulation of essential marker genes and cytokines in goblet cells under the inflamed UC condition.

Similar to other pseudotime methods, MARGARET assumes unidirectional cell differentiation where immature stem cells differentiate into more mature cell types. This assumption is violated for trans-differentiation and de-differentiation, events that can lead to ancestral cell-states and scRNA-seq data alone might be insufficient in characterizing such events (9). Recently, (39; 40) utilized naturally occurring somatic mutations for lineage tracing. It would be an interesting direction to extend MARGARET by incorporating auxiliary signals like lineage information and real-time information for elucidating reprogramming. Lastly, while we focus on scRNA-seq datasets in this study, it is worth noting that our framework can easily be extended for other single-cell omics as well as multi-omics datasets. The modular structure of our method also allows for effortless integration with other dimension reduction and omics-integration methods. Given the explosion of single-cell datasets fuelled by collaborative efforts such as Human Cell Atlas (HCA) project (41) and Human Biomolecular Atlas Program (HubMAP) (42), we anticipate MARGARET to be a valuable tool for multifaceted exploration of dynamic cellular processes from varied biological systems.

## Methods

### Preprocessing scRNA-seq data

We downloaded the filtered, normalized and log-transformed count matrices for early Human Hematopoiesis datasets (replicates 1 and 2) provided by (9) (see Data Availability). Pre-processed replicate 1 consisted of 5780 cells and 14651 genes while replicate 2 consisted of 6501 cells and 14913 genes. For each replicate, we performed imputation using MAGIC (43), and then computed 300 dimensional PCA embeddings on the imputed data as suggested in (9). For the embryoid body (EB) dataset, we downloaded the filtered, normalized and square-root transformed count matrix as provided in (14) (see Data Availability). The EB dataset consisted of 16821 cells and 17845 genes. We then performed initial denoising on the dataset to extract 50 dimensional PCA embeddings as performed in (14). For studying the epithelial differentiation in early intestinal development, we downloaded the filtered, normalized and log-transformed count matrix provided by (44). Top 2000 highly variable genes were identified in this dataset and then PCA was performed to compute 300-dimensional embeddings, which were used for further analysis. All preprocessing was performed using Scanpy (8).

### MARGARET Overview

The overall framework of MARGARET consists of two main steps: inferring the lower-dimensional cell-state manifold using preprocessed data and trajectory modelling.

### Inference of Lower-dimensional Cell-state Manifold

For inferring the cell-state manifold, Given preprocessed scRNA-seq data, we propose an unsupervised metric-learning-based approach to learn a meaningful lower-dimensional representation of each cell in the scRNA-seq dataset (see Fig. 1a). The central idea is to learn a non-linear manifold over cell representations such that the cells which belong to the same cell type (cluster) are packed compactly while the cells belonging to different clusters are far apart in the manifold. The key steps of our proposed approach are as follows:

1. **Computing initial clusters:** Given the preprocessed scRNA-seq data with *N* cells and an initial embedding of dimension *D*, we perform initial clustering on the data. The cluster label assigned to each cell is then treated as a pseudo-label, which is used in subsequent steps. The pseudo-labelled dataset is denoted as 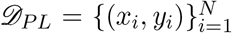, where the pair (*x*_*i*_, *y*_*i*_) represents the initial embedding and the pseudo-label of the *i*^*th*^ cell in the dataset respectively. It is worth noting that the choice of the clustering method in this step can be arbitrary. However, we advocate the use of community detection methods such as Louvain and Leiden clustering (45) in this work.
2. **Metric learning:** We then learn a low-dimensional representation of each pseudo-labelled cell using a non-linear mapping *f*_*θ*_ : ℝ^*D*^ → ℝ^*d*^ parameterized by *θ* where *d* ≤ *D*. In this work, we represent *f*_*θ*_ using a feed-forward deep neural network with parameters *θ*. Given this parameterization and a pseudo-labelled dataset 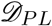, we learn a low-dimensional representation of each cell as follows: Given an initial representation (*x*_*i*_, *y*_*i*_) for the *i*^*th*^ cell, two additional cells (*x*_*j*_, *y*_*j*_) and (*x*_*k*_, *y*_*k*_) are sampled from 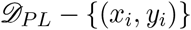 such that *y*_*i*_ = *y*_*j*_ and *y*_*i*_ ≠ *y*_*k*_. Let us assume that *f*_*a*_, *f*_*p*_ and *f*_*n*_ denote the low-dimensional embeddings of *x*_*i*_, *x*_*j*_ and *x*_*k*_ such that:

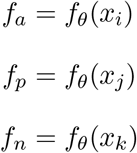

*θ* is updated such that the Triplet-Margin loss function (46) 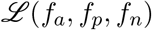 is minimized, where the loss 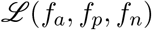 is given by:

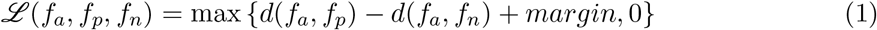

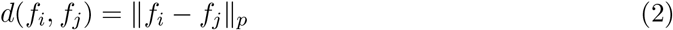 The value of *margin* and *p* are chosen as 1 and 2 respectively. Intuitively, for each cell *x*_*i*_ (denoting the anchor), we sample positive (*x*_*j*_) and negative samples (*x*_*k*_) such that the distance between the anchor and the positive sample is minimized while the distance between the anchor and the negative sample is maximized. This training step is repeated for *e* epochs, where *e* is a hyperparameter.
3. **Cluster refinement:** We generate the low-dimensional representations of all cells in the dataset using pre-trained *f*_*θ*_ from step 2. Using the updated low-dimensional embedding *f*_*θ*_(*x*_*i*_) for all *N* cells, cells are clustered again to generate refined cluster assignments, which act as new pseudo-labels for the cells 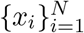.
4. **Episodic training:** The sequence of steps 2 and 3 form a single *episode*. We repeat steps 2 and 3 alternatively for a total of *E* episodes, where *E* is a hyperparameter. We can also alternate between the two steps until convergence which can be assessed by monitoring the quality of the clusters generated from step 3. For example, when using Leiden or Louvain clustering, convergence can be assessed by monitoring the *modularity* score of the refined clusters.
5. **Inferring final low-dimensional manifold:**: After training, the final low-dimensional representation, 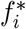 for cell *i* can be obtained by *f*_*θ\**_(*x*_*i*_) where *θ\** denotes the trained parameters of the deep neural network *f*. As a by-product of training, we also obtain the refined cluster assignments, 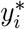 for each cell in the scRNA-seq dataset. The refined embeddings 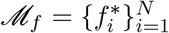 and cluster assignments 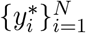 are used in subsequent steps of MARGARET.

### Network Architecture and Training Hyperparameters

We use a simple feed-forward deep neural network architecture in MARGARET consisting of 2-fully connected layers of sizes 128 and 64 respectively. The size of the input layer depends on the size of the embedding of the preprocessed scRNA-seq dataset. The number of neurons in the final output layer is the same as the size of the desired low-dimensional embedding which is a hyperparameter. In addition, each fully connected layer in MARGARET is followed by a Batch Normalization (BN) (47) layer followed by the ReLU activation. To regularize and enrich the intermediate representations, we use Dropout (48) after each Linear-BN-ReLU module. The dropout rate is set to 0.3 for all the experiments. Moreover, the layer sizes were also kept fixed for all the experiments.

During training, we used Stochastic Gradient Descent (SGD) with an initial learning rate of 0.01 to update the metric learner parameters. To adjust the learning rate during training, we used a poly-learning rate scheduler (49) with the following update schedule:

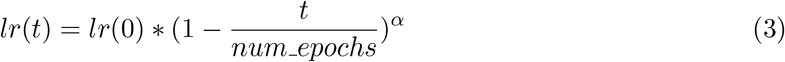

*α* = 0.9 was used during training.

### Trajectory Modelling

Given a low dimensional representation of cells, 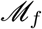 and refined cluster assignments 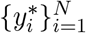, MARGARET learns a connectivity graph over the set of refined clusters similar to PAGA (16) to model the trajectory over the underlying dynamic process. The key steps involved in learning the trajectory are described below.

1. **Learning an undirected graph:** MARGARET first learns an undirected graph 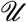 over the refined partition of cells 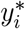. This learned graph 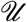 models the connectivity of the cell clusters and identifies the connected and disconnected neighborhoods in the cell-state manifold. To learn 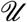, we first compute the k-nearest-neighbor (kNN) graph, 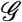 at the single-cell level using the learned low-dimensional manifold 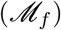. The resulting graph, 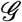 is represented as a *N* × *N* sparse adjacency matrix. We then assess the connectivity between two clusters *c*_*i*_ and *c*_*j*_ by introducing a novel measure of connectedness. Formally, we define the connectivity between two clusters *c*_*i*_ and *c*_*j*_ as:

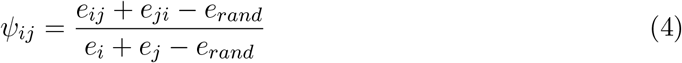

where *e*_*ij*_ denotes the number of kNN graph edges from cluster *c*_*i*_ to *c*_*j*_, *e*_*ji*_ denotes the number of kNN graph edges from cluster *c*_*j*_ to *c*_*i*_, *e*_*i*_ denotes the number of outgoing edges from cluster *c*_*i*_, *e*_*j*_ denotes the number of outgoing edges from cluster *c*_*j*_, and *e*_*rand*_ denotes the number of edges from cluster *c*_*i*_ to *c*_*j*_ and vice-versa under the random assignment of edges. Intuitively, when computing the connectivity between two clusters, we adjust the connectivity score to account for the random assignment of edges from cluster *c*_*i*_ to *c*_*j*_ to prevent spurious connections in 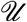. Following PAGA (16), we model the random assignment of edges between two clusters using a binomial distribution. In this scenario, *e*_*rand*_ is given by:

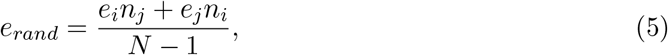

where *n*_*i*_ and *n*_*j*_ represent the number of cells (size) in the clusters *c*_*i*_ and *c*_*j*_ respectively and *N* represents the total number of nodes in the kNN graph (i.e. the number of cells). Given a threshold (lower-bound for *ψ*_*ij*_) *t*_*c*_, two clusters *c*_*i*_ and *c*_*j*_ are said to be connected when *ψ*_*ij*_ > *t*_*c*_. In this work we define a simple statistical test and compute the z-score between two clusters *c*_*i*_ and *c*_*j*_ given by:

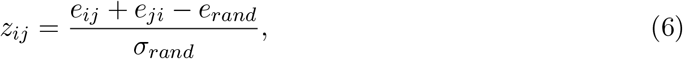

where *σ*_*rand*_ denotes the standard deviation of the binomial model and can be specified as:

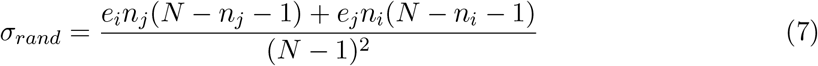 The statistical test proposed in Equation (6) is a direct consequence of the fact that under sufficiently large partitions, binomial random variables can be well approximated by a normal distribution. Therefore, the connectivity between two clusters *c*_*i*_ and *c*_*j*_ is given by *ψ*_*ij*_ when *z*_*ij*_ > *t*_*c*_ where *t*_*c*_ is a user-defined threshold.
2. **Pseudotime computation:** As in prior works (16; 9; 7; 50), MARGARET learns a temporal ordering over cells to uncover the dynamics of the underlying dynamic process. To determine the temporal order of the cells, for each cell, we infer *pseudotime*, which denotes the position of the cell in the underlying cell-state manifold representing the dynamic process. Given the kNN graph 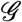 and a prior starting cell index *s*, one possible way to estimate the pseudotime can be to compute the shortest-path distance of each cell from the starting cell *s* because the shortest-path distances better approximate the geodesic distances in a non-linear manifold (51). However, the kNN graph 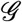 can be inherently noisy due to spurious connections between cells. Hence, directly computing the shortest-path distances using 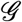 would give inaccurate estimates of the distance of each cell in the manifold from *s*. To mitigate this problem, we prune the kNN graph 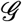 by using our undirected graph 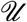 as a reference model. Formally, given an undirected graph 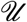 and the kNN graph 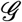, we prune an edge between two cells 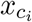 and 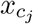 belonging to two clusters *c*_*i*_ and *c*_*j*_ respectively, iff 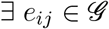 and *ψ*_*ij*_ = 0 where *e*_*ij*_ represents a (short-circuit) edge between cells 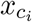 and 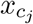 in the kNN graph 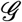. Given a pruned kNN graph 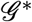, we compute the shortest-path distance of each cell from a user-defined starting cell *s* to infer the pseudotime for each cell in the scRNA-seq dataset.
3. **Orientation of Edges in Trajectory**: Given an undirected graph 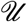 and the pseudotime 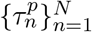, we compute the mean-pseudotime for each cluster *c*_*i*_ as:

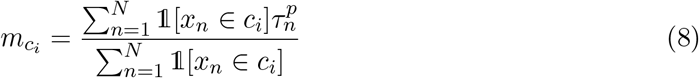 We then orient an edge from cluster *c*_*i*_ to *c*_*j*_ iff: 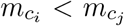 and *ψ*_*ij*_ ≠ 0 to obtain the final trajectory 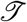 with node-set 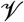 and directed edge-set 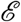.

### Prediction of Terminal States

Given a trajectory 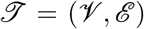, we compute the shortest-path betweenness (52; 53; 54) of every node in 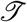. Formally, the shortest-path betweenness can be defined as:

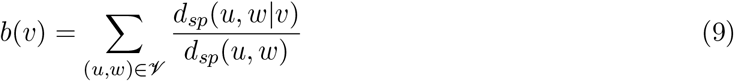

where *b*(*v*) is the betweenness for node *v*, *d*_*sp*_(*u, w*|*v*) represents the shortest-path distance between nodes *u* and *w* that passes through node *v* and *d*_*sp*_(*u, w*) represents the shortest-path distance between nodes *u* and *w*. Intuitively, the betweenness for any node 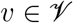 is the sum of the fraction of all-pairs shortest paths that pass through node *v* thus indicating its importance in the network. Given the shortest-path betweenness values 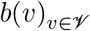, we compute the median (*b*_*med*_) and the median absolute-deviation (MAD) (*b*_*mad*_) of the betweenness values respectively. A node *v* is added to the set of terminal states if

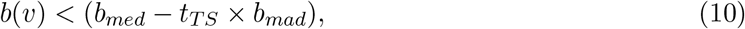

where *t*_*TS*_ is a user-defined scalar multiplier. Higher values of *t*_*TS*_ typically lead to nodes with no outgoing edges in 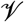 being selected as the terminal states. At the single-cell level, we select the cells having the maximum pseudotime value in each terminal state as the terminal cell for the underlying developmental process.

### Inferring Cell branch Probabilities and Differentiation Potential

Similar to (9), we model differentiation as a stochastic process on our learned cell-state manifold where the cells can follow the paths in the pruned kNN graph 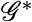 to reach any of the terminal states. Following (9), we model this stochastic process using an absorbing Markov Chain with the terminal cells acting as the absorbing states. Essentially, this formulation enables us to calculate the differentiation potential (DP) for each cell, a quantity that represents the potency of a cell to differentiate into specialized cell types. Given a set of terminal cells 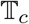, for each cell *i*, we compute the branch probabilities 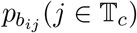, which represents the probability of cell *i* reaching a terminal cell *j*. Since DP of a cell 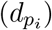 quantitatively characterizes the potency of a cell to maturate to different terminal states it can be obtained by computing the entropy of the branch probabilities of each cell as follows:

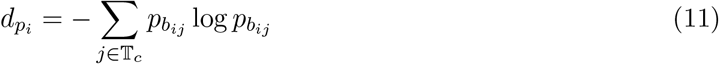

This is a reasonable way of modelling DP because cells with heterogeneous branch probabilities can be expected to have lesser potency of differentiating into diverse cell types. In the remainder of this section, we discuss the key steps involved in computing the branch probabilities for each cell *i*.

1. **Waypoint sampling for scalable modelling of DP**: Given the scRNA-seq dataset with *N* cells, fitting an absorbing Markov Chain can be computationally intractable for large *N*. To scale our absorbing Markov Chain model to large datasets, we sample a subset of cells *M*, from the scRNA-seq dataset such that *M << N*. We call these subset of cells as *landmarks* or *waypoints*. Specifically, given a set of *K* disjoint partitions of the embedding space *y*\*, we sample *k* waypoints per cluster by applying k-means++ (55) initialization for each cluster. We use the refined clusters obtained from the Metric-learning step for sampling the waypoints. Using such a scheme for waypoint sampling has two main advantages. Firstly, using a kmeans++ like scheme in each cluster ensures high intra-cluster waypoint coverage. Secondly, sampling waypoints from each cluster guarantees coverage of the entire embedding landscape. In contrast, (19; 56) use random sampling to compute waypoints which provides no coverage guarantees. Palantir (9) uses Max-min sampling (57) to compute waypoints which is more efficient than random sampling but might require a large number of waypoints to cover the embedding space.
2. **Computing Waypoint to Terminal Cell(s) probabilities**: Given a set of waypoints 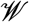 consisting of *M* waypoints, we compute a nearest-neighbor graph 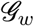 using the low-dimensional representations of 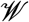. We then prune 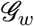 using the short-circuit edge pruning (See Pseudotime computation) with the undirected connectivity graph 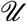 as a reference model. Furthermore, following (9), we also remove edges in 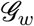 which violate the pseudotime ordering between waypoint cells. Formally, an edge 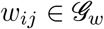 from waypoint *w*_*i*_ to *w*_*j*_ is pruned iff:

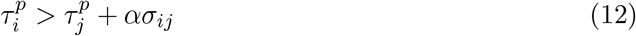

where 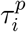 and 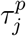 represent the pseudotime for waypoints *w*_*i*_ and *w*_*j*_ respectively and *σ*_*ij*_ represents the scaling factor for cell *w*_*i*_ given by the distance of *w*_*i*_ to its *l*^*th*^ neighbor in 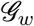. The statistical test in Equation 12 differs from the formulation in (9) in terms of the parameter *α*. We found that parameterizing Equation 12 with the user-defined parameter *α* provides an additional flexibility in controlling the number of edges that are pruned which is an important aspect of fitting an absorbing Markov Chain. Given a pruned waypoint nearest-neighbor graph 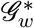 and a set of terminal cells 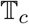, we row-normalize 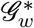 to obtain the transition matrix *T*. For two waypoints *w*_*i*_ and *w*_*j*_, the entry *t*_*ij*_ in *T* represents the probability of transitioning from waypoint *w*_*i*_ to *w*_*j*_. An absorbing Markov Chain is specified by a transition matrix of the form 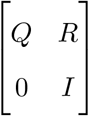 where *Q* represents the transition probabilities of moving between intermediate states and *R* represents the transition probabilities of moving from intermediate states to the terminal states. We represent our transition matrix *T* using this formulation and compute the waypoint to terminal cell branch probabilities as follows:

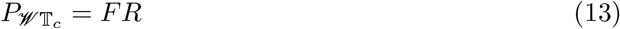

where *F* is the fundamental matrix given by *F* = (*I* − *Q*)^−1^. Since our Transition matrix *T* can be sparse, we recommend computing the fundamental matrix using the Moore-Penrose pseudoinverse to avoid numerical issues.
3. **Computing Cell to Waypoint Connectivity**: Given the pruned nearest neighbor graph 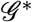, we compute cell to cell connectivity using a local random walk (LRW) (58) formulation. LRW is a quasi-local method to estimate connectivity between nodes in a graph based on limiting a random-walk to a fixed number of steps. Hence, the approach is computationally much more efficient than using a global random walk until convergence. Formally, the LRW connectivity *ζ*^(*i,j*)^ between two cells indexed by *i* and *j* is given by:

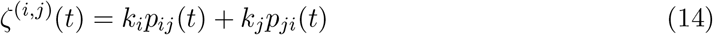

where *p*_*ij*_(*t*) and *p*_*ji*_(*t*) represent the probabilities obtained when moving from cell *i* to *j* and vice-versa at time *t* respectively. The constants *k*_*i*_ and *k*_*j*_ are set to 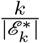 where *k* is the number of nearest neighbors and 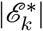 is the total number of edges in 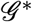. Given the similarity matrix 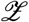 representing LRW-based connectivities between different cells we can index 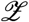 to compute cell to waypoint similarities, 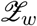.

Given the cell to waypoint similarities 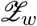, and waypoint to terminal state probabilities 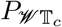, we compute the cell to terminal states branch probabilities 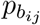 by a simple projection:

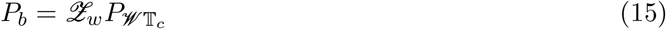

where *P*_*b*_ is a 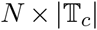 matrix representing branch probabilities for each cell. We then compute the DP for each cell using Equation 11.

### Inference of Gene Expression Trends Along Lineages

To visualize the variation in the expression of a gene *g* across different lineages with pseudotime, we fit generalized additive models (GAMs) (59) to the gene expressions and pseudotime values (*g*_*i*_, 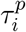) of cells along a particular lineage *j* weighted by their branch probabilities 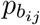 for that lineage (see Supplementary Note 2 in (9) for more details). We imputed the preprocessed expression value of gene *g* using MAGIC (43), when computing the lineage trends for the hematopoiesis and EB datasets. No imputation was performed for the colon IBD dataset. We used the LinearGAM implementation available in the *pyGAM* package (60) to fit the lineage trends with the regularization penalty set to 10 and the order of the splines set to 4 for all the experiments.

### Differential Expression and Gene Ontology analysis

We used the Wilcoxon-rank sum test available in *Scanpy* (8) to estimate differentially expressed (DE) genes for each cluster and the Benjamini-Hochberg correction for adjusting the p-values. All genes were ranked in the DE analysis, which is the default behavior in Scanpy 1.7.2. To assess the functional significance of MARGARET inferred clusters, we performed Gene Ontology (GO) analysis using the *gprofiler-official* (61) package. To determine the GO terms for a cluster, we selected the top DE genes with a log fold change value greater than 1.0 and the adjusted p-value less than 0.05. In case more than 500 genes were included, we selected the top 500 genes to obtain a list of GO terms associated with that cluster.

### Performance Metrics for Trajectory Inference

Here we describe the quantitative performance metrics used for evaluating the trajectory inferred by a TI method:

1. *Ipsen-Mikhailov (IM) Distance*: We used the IM distance (62) metric for global topology comparison between PAGA and MARGARET. Before computing the IM distance, the trajectories inferred from both the methods (including the ground-truth trajectory) were coerced to an undirected graph. Formally, the Laplacian spectra for a graph 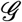 can be specified as a mixture of Lorentz distributions (Equation (16)) with the same half-width at half-maximum *γ* and centered at the frequencies *ω*_*k*_ given by 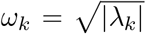, where *λ*_*k*_ is the *k*^*th*^ eigenvalue of the graph Laplacian of 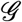. The constant *C* is a normalization constant for the resulting probability distribution.

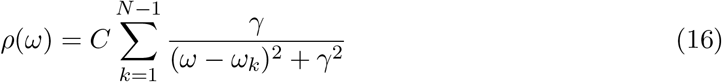 The IM distance measures the difference between the Laplacian spectra of two graphs as follows:

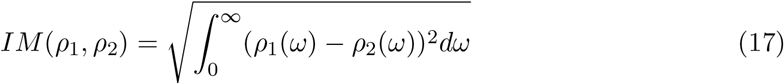 Since it is a measure of distance between the spectra of two graphs, a low IM distance corresponds to greater similarity in the global topology of the two graphs.
2. *Rank Correlation Metrics*: We used two rank correlation metrics to compare the ground truth ordering and pseudotime orderings inferred by the TI methods. More specifically, we used the *Kendall’s Tau (KT)* and the *Spearman’s Rank (SR)* correlation coefficients to estimate the similarity in rank orderings of the data. As suggested in (63), the KT correlation coefficient can be specified as follows:

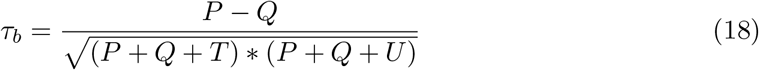

where *P* and *Q* represent the number of concordant and discordant pairs, respectively. *T* and *U* represent the number of ties in the two orderings, respectively. The SR correlation coefficient is simply defined as the Pearson’s correlation coefficient applied to the ranks of the variables measured in the two orderings. We used the *scipy.stats* package to compute both KT and SR correlation coefficients.
3. *Clustering Metrics*: We used *Adjusted Rand Index* (ARI) and *Normalized Mutual Information* (NMI) metrics to assess the clustering performance of MARGARET embeddings. The ARI metric corrects the Rand Index for chance and is specified by the following formulation:

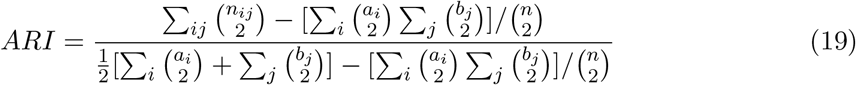

where *a*_*i*_, *b*_*j*_ and *n*_*ij*_ are values from the contingency table which is used to estimate the overlap between two partitionings. The NMI metric between two clusterings *C*_1_ and *C*_2_ can be formulated as follows:

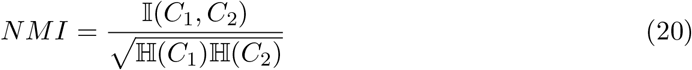

where 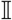 is the mutual information between the two clusterings *C*_1_ and *C*_2_ and 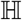 is the Shannon-Entropy of the clustering.

### Visualization of Embedding

We used UMAP (12) (with default parameters) for visualization of all the datasets except the early Hematopoiesis scRNA-seq data provided by (9), for which we used tSNE (64) visualization with a perplexity value of 180 for replicate 1 and 150 for replicate 2.

### Computing Crypt-axis Score

As suggested in (15), we used the expression of the following genes to define the crypt-axis (CA) score: *SEPP1*, *CEACAM7*, *PLAC8*, *CEACAM1*, *TSPAN1*, *CEACAM5*, *CEACAM6*, *IFI27*, *DHRS9*, *KRT20*, *RHOC*, *CD177*, *PKIB*, *HPGD* and *LYPD8*. The final crypt-axis score for each cell was then computed by summing over the normalized expression (between 0 and 1) values of each gene included in our set.

### Simulation of Benchmark Datasets

To benchmark MARGARET’s performance on several aspects of trajectory inference, we generated a suite of synthetic single-cell gene expression datasets representing different complex trajectory models. We used *dyntoy* (13) (https://github.com/dynverse/dyntoy) to generate the synthetic benchmark (see Supplementary Table 3 for the details of different synthetic datasets) spanning multifurcating and disconnected topologies. For benchmarking MARGARET, PAGA (16), and Monocle3 (7), all the simulated datasets were subjected to the same data preprocessing steps following Seurat (65): removal of genes expressed in less than 3 cells, normalization, log transformation with a pseudo count of 1.0, and finally z-score normalization. For benchmarking *Palantir*, the steps proposed by the authors in (9) were used to preprocess the simulated datasets.

### Data Availability

All the experimental biological datasets used in this study are openly available and can be referenced from Supplementary Table 3. The human hematpoiesis dataset is available through the Human Cell Atlas portal at https://prod.data.humancellatlas.org/explore/projects/29f53b7e-071b-44b5-998a-0ae70d0229a4. The scRNA-seq and bulk RNA-seq datasets for the embryoid body dataset can be accessed via the Mendeley Data repository at https://doi.org/10.17632/v6n743h5ng.1. The scRNA-seq data for colon differentiation can be accessed using the GEO accession number GSE116222. All the real datasets used for demonstrating the clustering efficiency of MARGARET can be accessed via the *scvi-tools* package (66).

### Code Availability

MARGARET has been implemented in Python and is freely available at https://github.com/Zafar-Lab/Margaret.

## Acknowledgements

This work was partially funded by the IIT Kanpur initiation grant to H.Z.

## Author contributions

K.P. and H.Z. designed the study, K.P. and H.Z. developed the model and algorithm. K.P. implemented the software and performed all experiments. All authors wrote and approved the paper.

