## Supplementary Information for "Inference of cell state transitions and cell fate plasticity from single-cell with MARGARET"

October 22, 2021

**Supplementary Figures** **8**

**Supplementary Tables** **36**

### Supplementary Notes

#### Supplementary Note 1: MARGARET applied to simulated datasets

##### Qualitative comparisons on a simulated cyclic dataset

We also applied MARGARET to a simulated dataset with a cyclic ground-truth trajectory (Cyclic_1) (Supplementary Fig. 4a, Supplementary Table 3a). For this dataset, MARGARET and PAGA were able to capture the global topology accurately while Palantir and Monocle3 failed to capture the cyclic structure (Supplementary Fig. 4b).

##### DP comparison between MARGARET and Palantir

We also compared the DP inference by MARGARET and Palantir for two complex simulated disconnected trajectories (Disconnected_5 and Disconnected_6) (Supplementary Fig. 5, Supplementary Table 3a). For both the datasets, MARGARET accurately captured the DP trends in different disconnected components and MARGARET inferred DP showed high negative Spearman-Rank correlation with the ground-truth pseudotime. DP is expected to have negative correlation with pseudotime (1) since as cells differentiate proceeds towards terminal states, pseudotime increases while the DP of the cell decreases. In contrast, Palantir's DP inference performed poorly for disconnected trajectories as observed from the correlation analysis between DP and ground truth pseudotime (Supplementary Fig. 5). The poor performance of Palantir could be the result of its single component assumption when inferring the DP which does not extend to multiple independent disconnected components within the same dataset.

#### Supplementary Note 2: Clustering quality analysis using MARGARET

##### MARGARET learns modular cluster representations during episodic training

Since MARGARET's metric-learning approach is aimed at inferring a lower-dimensional cell-state manifold where the distinct cell states are represented by compact cell clusters, we evaluated the compactness of the inferred clusters by tracking the clustering modularity scores and the number of clusters inferred at the end of each episode during training of two single-cell RNA sequencing human hematopoiesis datasets (1) using Phenograph (2) across Louvain and Leiden clustering backends. For both datasets, at the end of each training episode, MARGARET improved upon the modularity score of the previous episode before finally converging (Supplementary Fig. 6). The clustering modularity score can also be used for estimating MARGARET's convergence. Interestingly, the number of inferred clusters did not always increase monotonically with the number of episodes (Supplementary Fig 6a-c), suggesting that the increase in modularity is due to the improved clustering quality at each episode.

##### MARGARET can refine the single-cell embeddings learned using other methods

To investigate the ability of MARGARET's unsupervised metric learning-based approach, we initialized MARGARET with $d$-dimensional cell embeddings (as obtained from a linear or nonlinear dimension reduction method such as PCA or scVI (3)) to infer MARGARET-refined $d$-dimensional embeddings. The quality of MARGARET-inferred cell embeddings was evaluated by comparing its cell-type clustering performance (as measured by adjusted rand index (ARI) and normalized mutual information (NMI) metrics) against that of the initial embeddings on a suite of biological datasets for which the ground-truth clustering annotations were available (Methods). As compared to scVI-inferred cell embeddings, MARGARET's refined embeddings achieved higher ARI and NMI scores across all the datasets (Supplementary Fig. 7a). Similar results were observed using a PCA-based initialization (Supplementary Fig. 7b), which suggests that MARGARET's metric learning-based approach can refine the latent representations captured during earlier dimension reduction stages.

#### Supplementary Note 3: Application of MARGARET to human hematopoiesis (Extended results)

##### Characterization of heterogeneity in DC lineage

In the DC lineage for replicate 1, MARGARET inferred cluster 7 expressed markers for both cDCs (*ITGAX, CLEC10A*) and pDCs (*IRF7, IRF8, IL3RA*) (Fig. 3d) while cluster 23 only expressed pDC markers. We made a similar observation in replicate 2 with clusters 1 and 20 equivalent to clusters 7 and 23 in replicate 1 , respectively. To characterize the heterogeneity in the DC lineage at a finer resolution, we combined the cells in the DC lineage from clusters 4, 7, and 23 in replicate 1 and clusters 5, 1, and 20 in replicate 2 and analyzed the resulting 1406 cells using MARGARET. Fig. 4d shows the MARGARET inferred trajectory, consisting of 10 clusters (DC0-DC9). Since MDPs give rise to DC populations, based on the expression of MDP-specific markers *CSF1R*, and *ITGAX* (4) (Fig. 4b) we inferred cluster DC7 as the starting cluster for our analysis. Based on manually curated set of markers specific to pDCs and cDCs through prior literature review (5; 6; 7), we then identified pDCs marked by high expression of *E2-2* (*TCF4*) TF, its target TFs *IRF7*, *SPIB*, and pDC-specific marker genes *LILRA4* (*ILT7*) (8) and *PACSIN1* (9), in cluster DC0 (Fig. 4e (top), f) which was a terminal state. *E2-2* expression serves as a key event in pDC cell fate choice (10), *IRF7* is a key regulator of IFN expression and has been shown to be highly expressed in pDCs as compared to other cell types (11). Similarly, we localized the cDC lineage by observing high expression of *ITGAX* (*CD11b*), *ID2*, and *CD1c* (12) in clusters DC3 and DC8 (Fig. 4f). Furthermore, we observed high expression of cDC2-specfic TFs *NR4A3, SREBF2* and marker genes *CLEC10A, CD1E, CLEC12A, CX3CR1* (6) (Fig. 4e (bottom), f) and negligible expression of cDC1 marker gene *CLEC9A* (13) in clusters DC3 and DC8 (data not shown) indicating the presence of cDC2 and absence of cDC1 cells in these clusters.

##### MARGARET characterized erythroid-megakaryocytic lineage

We also characterized the erythroid-megakaryocytic lineage branching in MARGARET inferred trajectory. Both erythroid and megakaryocytic commitment were associated with a sharp decrease in DP (Supplementary Fig. 15). In the erythroid lineage, this decrease in DP was concordant with the elevated expressions of TFs *GATA1*, *KLF1*, and *MYB*, which are known to play crucial roles in erythropoiesis: *GATA1* is indispensable for erythropoiesis (14), *KLF1* modulates erythroid cell differentiation by regulating erythroid precursor genes and also antagonizes megakaryocyte differentiation (15; 16), *MYB* enhances erythropoiesis by suppressing megakaryopoiesis (17). Expression of these TFs also highly correlated with the erythroid branch probabilities ($> 0.9$) indicating their crucial regulatory role in erythroid commitment (Supplementary Fig. 16a). In the megakaryocyte lineage, the drop in DP was concordant with increasing expression of transcription factors *PBX1*, *FLI1*, and *MEIS1* (Supplementary Fig. 15), which were also closely correlated ($> 0.9$) with megakaryocytic branch probabilities (Supplementary Fig. 16b). These TFs are known to play central role in megakaryopoiesis: *FLI1* and *PBX1* are essential TFs for megakaryocyte differentiation (15; 18), and *MEIS1* is essential for fetal megakaryopoiesis (19). The cluster 15 (in replicate 1) at which the erythroid and megakaryocytic lineages diverged, expressed both *GATA2* (driver of erythroid commitment (20)) and *CD41* (responsible for megakaryocytic lineage commitment (15)), as well as genes like *SLC14A1* and *VWF*, which are responsible for a continuous transition from megakaryocyte-erythroid progenitors (MEP) to erythroid and megakaryocyte progenitors respectively (21) suggesting the presence of MEPs in cluster 15.

#### Supplementary Note 4: Application of MARGARET to embryoid body dataset

We identified the major lineages recovered by MARGARET in early human embryogenesis by examining the expression of essential marker genes for major lineages previously reported in the literature for this dataset (22) (Fig. 5d, Supplementary Fig. 17a,b). This preliminary analysis revealed the presence of endoderm (EN), mesoderm (ME), neural crest (NC), neuroectoderm (NE), and neuronal subtypes (NS) (including neural progenitors (NP)) lineages along with ESCs in the data. To further validate the inferred lineages, we grouped MARGARET clusters based on their lineage information to obtain five major clusters (Supplementary Fig. 18a) for which we performed differential expression (DE) analysis (Supplementary Fig. 18b). Gene ontology (GO) analysis (Methods) of the DE genes for these combined clusters (Supplementary Fig. 18c) revealed major functional differences between these clusters, with the enrichment of GO terms corresponding to these major lineages suggesting the validity of the inferred lineages (Supplementary File 1).

Due to the presence of ESCs in cluster 6 as marked by the high expression of *POU5F1*, *NANOG* (essential for maintaining pluripotency in ESCs (23)), and *DDPA2/4* (Fig. 5d,f), we selected a starting cell from cluster 6 for further trajectory analysis, including the inference of pseudotime and DP. MARGARET inferred pseudotime (Fig. 5c) followed the progression of cell types, where the pseudotime increased as cells progressed towards more specialized cell types from *POU5F1* enriched ESCs.

Furthermore, MARGARET recovered a detailed lineage specification map of embryoid bodies in a fully unsupervised manner (Fig. 5e). We further characterized the MARGARET inferred clusters for specific cell types. In the ectoderm lineage: neuroectoderm, neural crest, and neuronal subtype clusters were detected as terminal states. In the mesoderm lineage, MARGARET identified hemangioblasts (H), cardiac precursors (CPs), and smooth muscle precursors (SMPs) as the terminal states. Lastly, a single cluster in the endoderm lineage was also identified as a terminal state. To characterize the recovered lineage map, we explored the expression trends of key marker genes for the terminal cell types (Fig. 5f). The expression of ESC marker gene *POU5F1* decreased with pseudotime in all lineages. *CD34* was selectively upregulated in hemangioblasts, while *TNNT2*, and *TBX18* were upregulated in the CPs and SMPs respectively. *KLF5*, and *SOX10* were upregulated in the EN, and NC lineages respectively. *LHX5* was initially upregulated in the NS, NC, and NE lineages but was subsequently downregulated in the NS and NC lineages suggesting its importance in NE lineage commitment. While we identified five NS clusters, *ONECUT1* was upregulated in NS-5, the terminal neuronal subtype cluster.

The probability of cells branching towards a specific lineage as inferred by MARGARET increased towards later stages in cell differentiation (Supplementary Fig. 19). For this dataset also, DP decreased with an increase in pseudotime (Supplementary Fig. 20a) with the ESC cluster having the highest DP, followed by the transitional cell types and the terminal states having the lowest DP (Supplementary Figs. 20-21). Thus decrease in DP was concordant with the major lineage commitments in EB differentiation.

In the ectodermal lineage, ESCs differentiate into preneuroectoderm cells (showing downregulation of *POU5F1* (Fig. 5f)), which give rise to neuroectoderm cells (expressing *LHX2/5, SIX3* (Fig. 5d,f)). Similar to (22), MARGARET was able to identify the bipotent precursors (cluster 9 expressing *HOXA2, HOXB1* and *OLIG3* (Fig. 5d)) that originated from the neuroectoderm cells expressing *GBX2* and bifurcated from cluster 9 into the neural crest and neuronal sub-lineages (Fig. 5e). Further characterization of this branching revealed correlation between a decrease in MARGARET inferred DP and the up/down regulation of important TFs in the neural crest and neuronal lineages (Supplementary Fig. 22). The DP drop in the neural crest lineage was concordant with the upregulation of canonical TFs *SOX9/10* (24; 25) (Fig. 5d,f, Supplementary Fig. 22a,b (right)) while neuronal-subtype cluster 3 exhibited upregulation of TFs *SOX1* and *LHX2* (Fig. 5d, Supplementary Fig. 22a,b (left)), which have been shown to be important for subtype specification in certain types of neurons (26; 27). GO analysis at finer resolution (Fig. 5g) also revealed the enrichment of both neural crest and neuronal differentiation-related functions in cluster 9 further validating its bi-potency. We next validated the neural crest sub-branch detected by MARGARET using the bulk RNA-seq data provided by (22) for FACS purified $CD49d^{+}CD63^{-}$ cells. Correlation analysis between scRNA-seq profiles of cells in the EB dataset with the bulk RNA-seq expressions corresponding to $CD49^{+}$ cells revealed the highest correlation in the neural crest lineage which also showed high *ITGA4* expression (Supplementary Fig. 23a), suggesting the accurate localization of the neural crest cells in MARGARET trajectory.

#### Supplementary Note 5: Applying MARGARET for studying colon differentiation

For the healthy colon, MARGARET identified 15 clusters (Fig. 6a) and the inferred trajectory accurately delineated the absorptive and the secretory lineages (Fig. 6f). Projection of crypt-axis (CA) scores (28) (see Methods) for each cell on the 2D representation of the MARGARET-inferred cell embedding (Fig. 6d) revealed the cells in the absorptive lineage to have higher CA scores as compared to stem cells or cells in the secretory lineage indicating the presence of these cells towards the crypt-top indicating that MARGARET correctly localized the cell-types within the trajectory. We then investigated the expression of key marker genes (curated from prior literature (28; 29)) for different cell types in the absorptive and secretory lineages (Fig. 6e). We identified absorptive progenitors, colonocytes, crypt-top (CT) colonocytes, and BEST4/OTOP2 cells in the absorptive lineage; and secretory progenitors, goblet cells, and enteroendocrine cells (EECs) in the secretory lineage. Based on high expression of stem cell markers genes *MLEC* and *LGR5* in cluster 0 (Fig. 6e), we selected a cell from this cluster as the starting cell for the subsequent inference of pseudotime (Fig. 6b) and DP (Fig. 6c). Terminal state prediction using MARGARET further revealed four terminal states namely: EECs (cluster 14), goblet cells (cluster 5), BEST4/OTOP2 cells (cluster 13) and CT colonocytes (cluster 3). MARGARET inferred branch probabilities (Supplementary Fig. 24) and gene expression trends (Fig. 6h) in the detected lineages further validated the cell populations detected in the absorptive and the secretory cell lineages as the marker genes *MUC2*, *SCGN*, *AQP8*, and *BEST4*, were selectively upregulated in the goblet (30), EECs (31), CT colonocytes (32), and BEST4/OTOP2 cell (28) lineages respectively.

#### Supplementary Figures

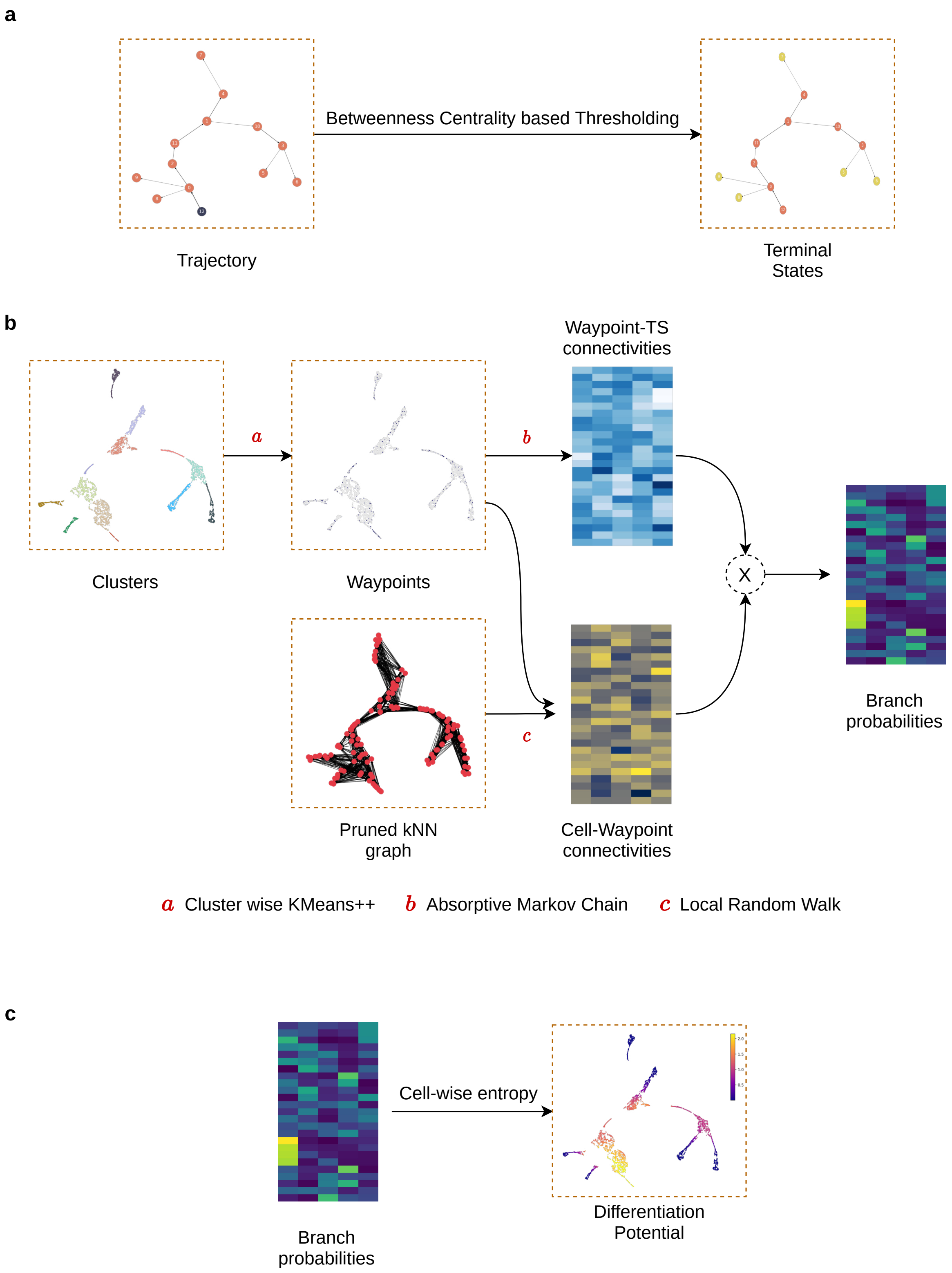

Supplementary Figure 1: **Prediction of terminal states and computation of differentiation potential (DP) in MARGARET.** (a) Prediction of terminal states using betweenness centrality followed by thresholding. (b) Computation of cell branch probabilities - probabilities for a cell differentiating into the terminal states by following the paths along the trajectory. (c) Computation of differentiation potential from cell branch probabilities (DP values are projected on the cellular embedding space). All illustrations (except the branch probabilities and connectivities) were generated by running MARGARET on a synthetic multifurcating dataset.

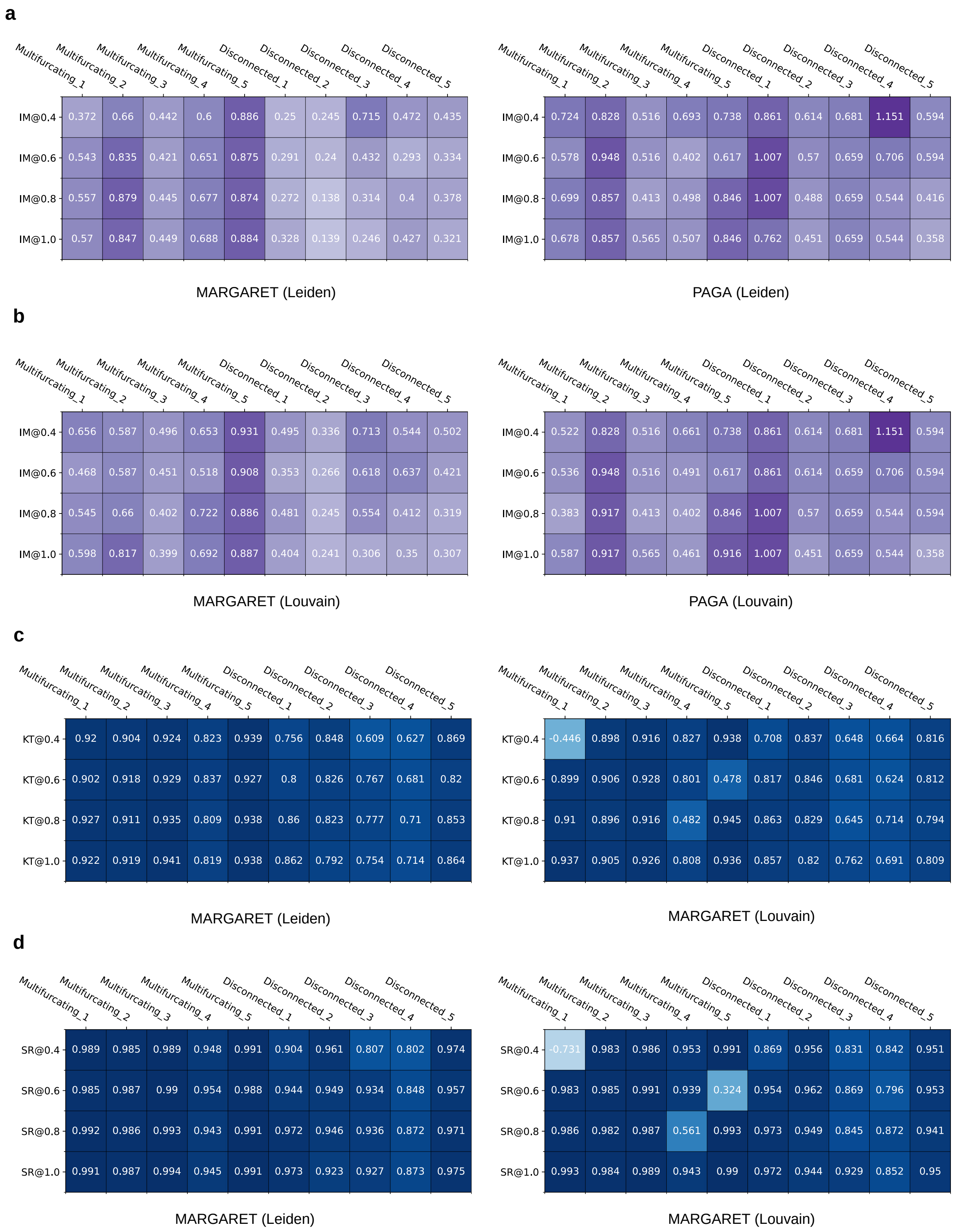

Supplementary Figure 2: **Quantitative Illustration of MARGARET's performance on the simulated benchmark.** The fractional suffix following @ denotes the resolution of clustering. (a) Annotated heatmaps illustrating the comparison between MARGARET and PAGA on the Global topology task when using Leiden clustering for community detection. IM stands for the Ipsen-Mikhailov (IM) distance. Lighter values indicate better performance. (b) Same as (a) but using Louvain clustering for community detection. (c) Quantitative results on the pseudotime ordering task using MARGARET. KT denotes the Kendalls-Tau coefficient. Darker colors indicate better performance. (d) Same as (c) but using the Spearman's Rank (SR) correlation metric.

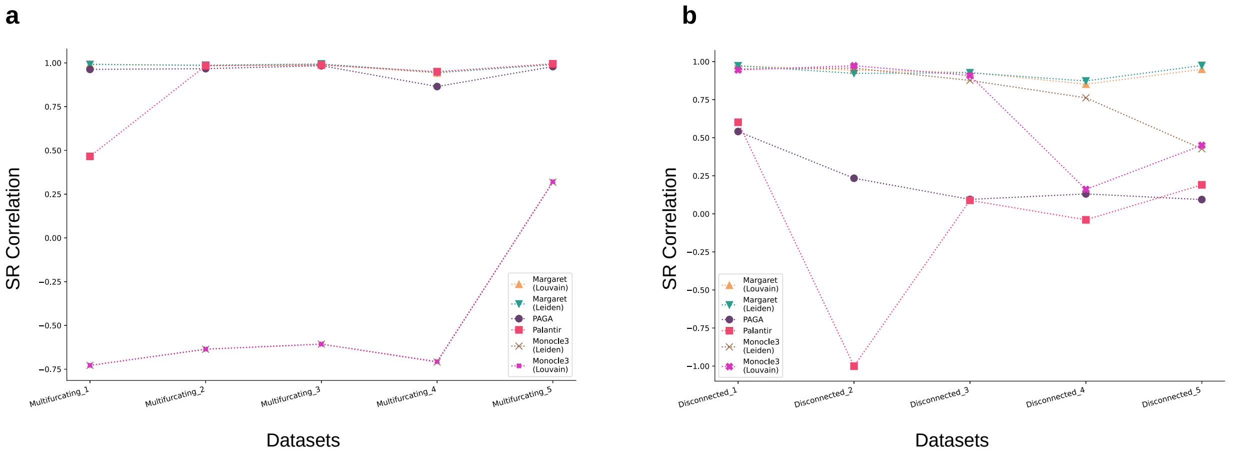

Supplementary Figure 3: **MARGARET outperforms other TI methods on the pseudotime ordering task.** (a) Pseudotime ordering comparison on five simulated multifurcating datasets between MARGARET, PAGA, Palantir and Monocle3 using the Spearman-Rank correlation for Louvain and Leiden clustering schemes (Higher is better). (b) Same as (a) but for the disconnected benchmark.

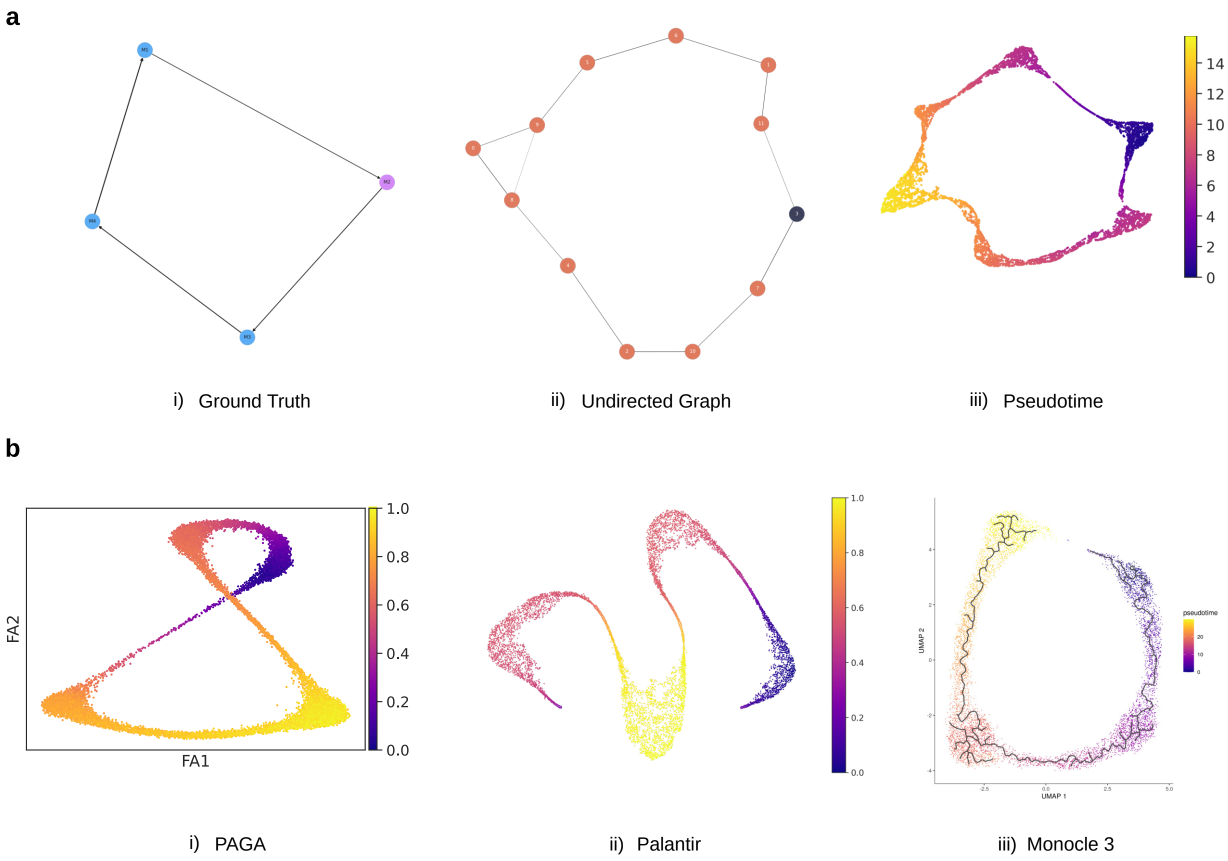

Supplementary Figure 4: **MARGARET's performance on cyclic trajectories** (a)(i) Reference cluster-connectivity graph for the synthetic cyclic dataset. (ii) Cluster connectivity graph inferred by MARGARET. (iii) MARGARET computed pseudotime projected on single-cell embeddings. (b)(i, ii, iii) Pseudotime projected on single-cell embeddings, computed using PAGA, Palantir and Monocle3, respectively.

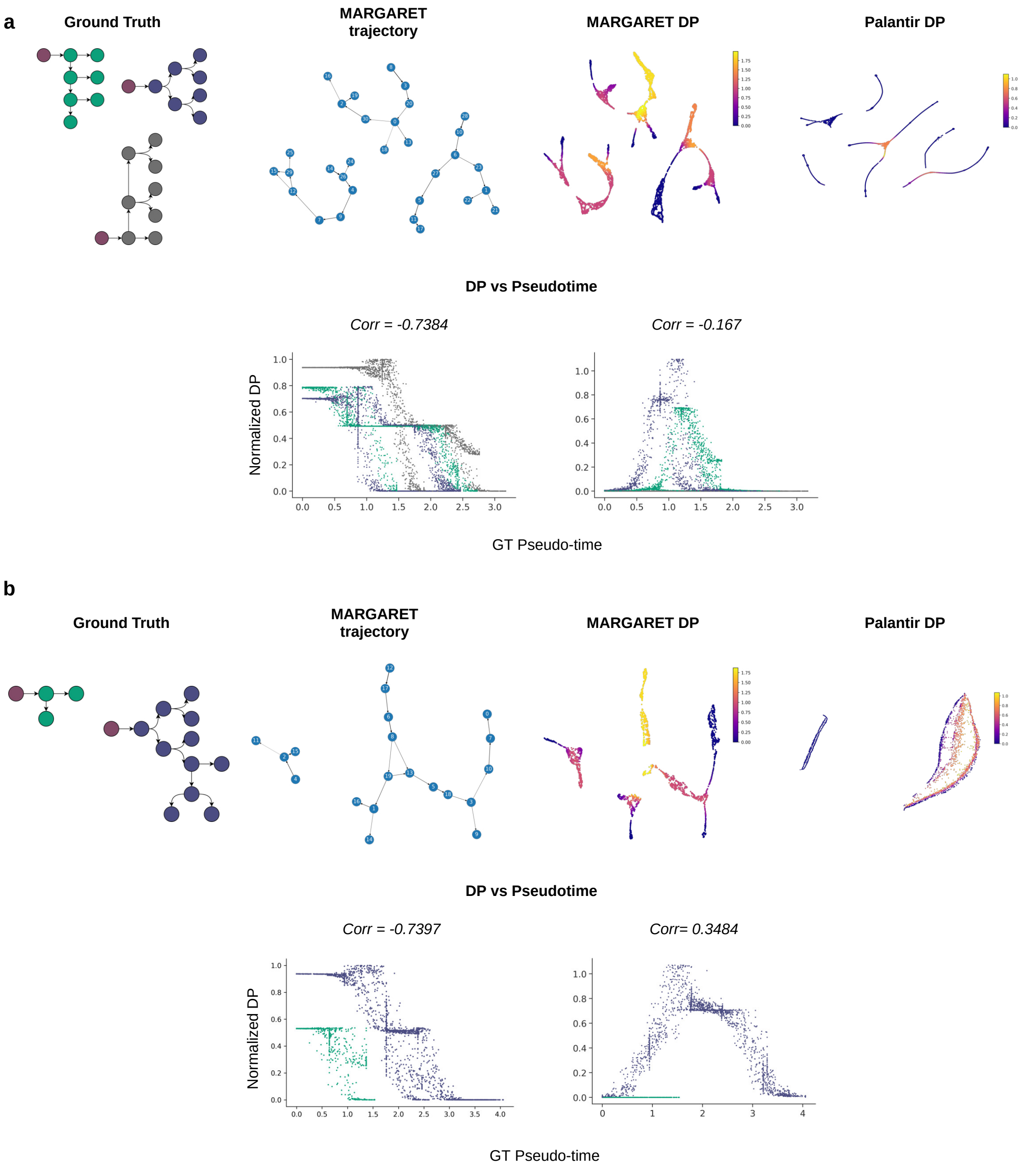

Supplementary Figure 5: **MARGARET inferred DP generalizes to disconnected trajectories.** a) (Top-Left) Schematic of a ground truth simulated dataset with three disconnected components (7500 cells). The starting cells are shown in red color while the trajectories can be uniquely identified by three different colors. (Top-Middle-1) MARGARET inferred trajectory. (Top-Middle-2) MARGARET inferred DP for this dataset. Inferred DP trends are qualitatively consistent across independent components. (Top-Right) Palantir inferred DP. The results are qualitatively inconsistent as Palantir is unable to capture the disconnected nature of the underlying trajectory. (Bottom) Scatter plot of Normalized DP vs ground truth pseudotime for both MARGARET (Left) and Palantir (Right). MARGARET shows a higher negative Spearman-rank correlation with the ground truth pseudotime as compared with Palantir. The cells in the scatter-plot are colored by their disconnected component id in the ground-truth trajectory. (b) Same as (a) but for a disconnected dataset with two components (2500 cells)

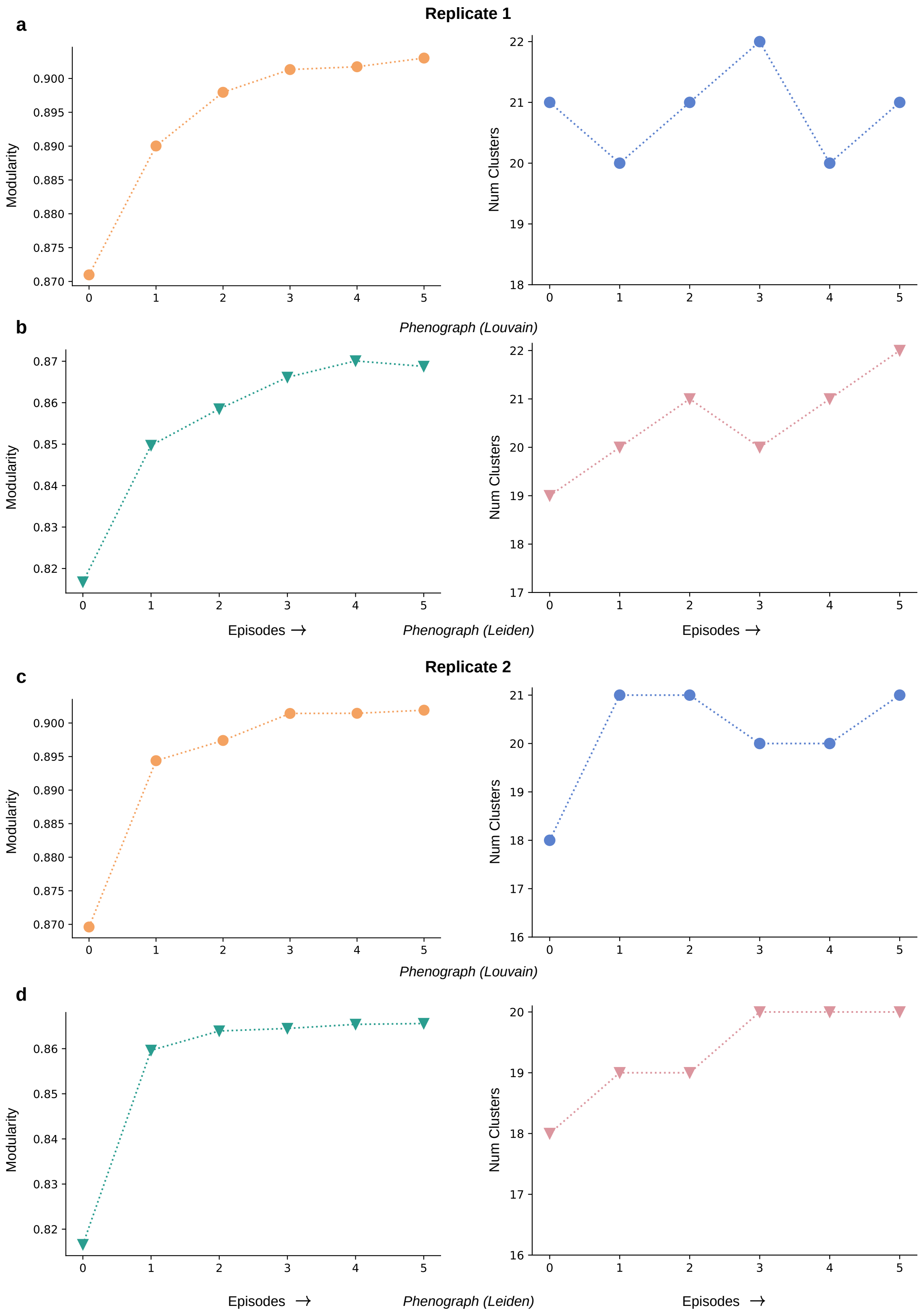

Supplementary Figure 6: **MARGARET improves the cluster compactness after each training episode.** (a) (Left) Change in the modularity score between training episodes for Replicate 1 in the scRNA-hematopoiesis dataset. (Right) Change in the number of clusters between training episodes using Phenograph clustering with a Louvain backend for Replicate 1. (b) Same as (a) but repeated using Phenograph clustering with a Leiden backend. (c) Same as (a) but using Replicate 2 in the scRNA-hematopoiesis dataset. (d) Same as (c) but using Phenograph clustering with a Leiden backend. The improvement in the modularity score is consistent across replicates and different clustering schemes.

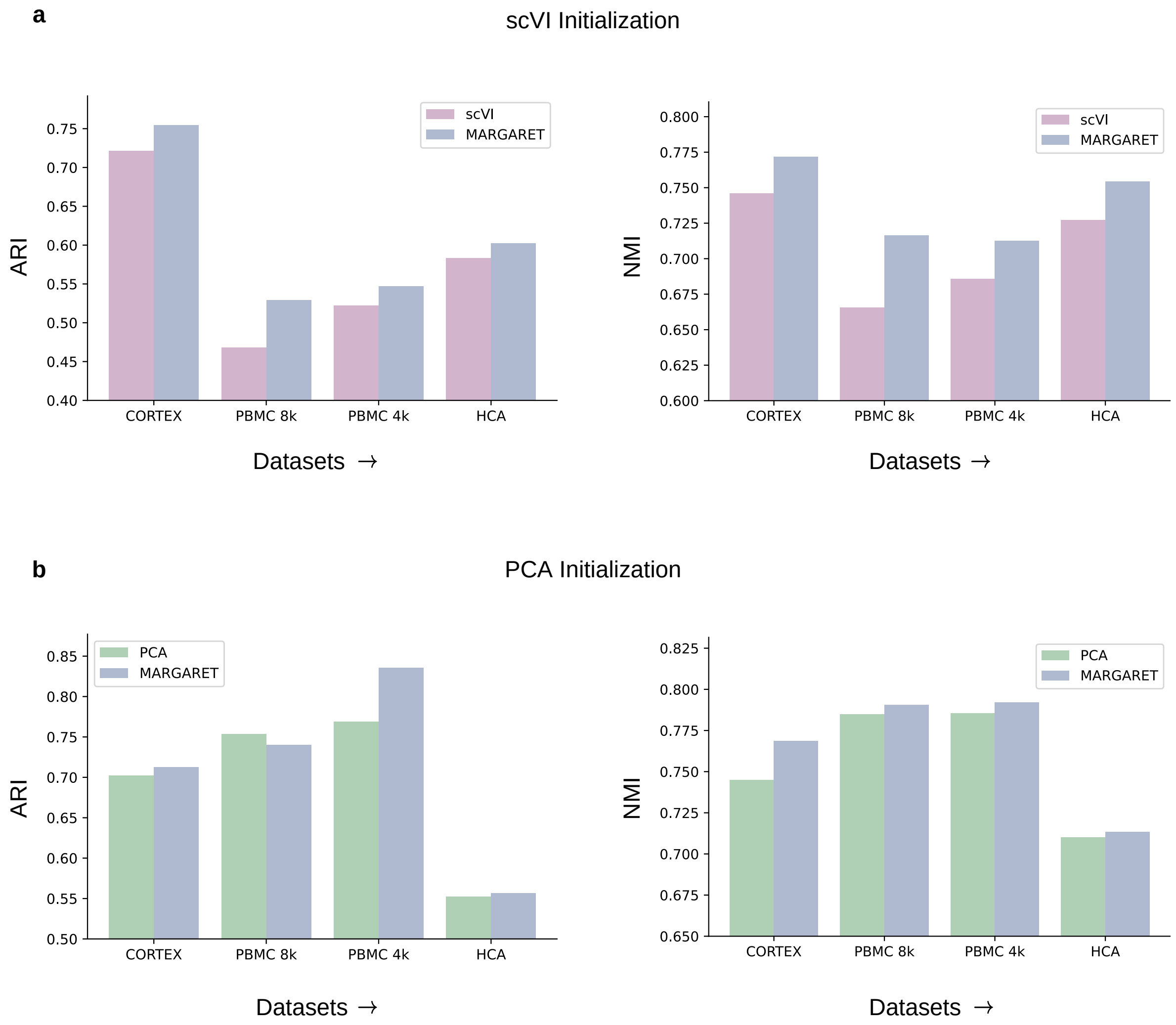

Supplementary Figure 7: **MARGARET improves upon the cell embeddings learned by other dimension reduction methods.** a) Comparison of MARGARET's clustering performance againsth that of scVI on experimental biological datasets (with gold standard cluster labels available) based on (Left) Adjusted Rand Index (ARI) and (Right) Normalized Mutual Information (NMI). b) Same as a) but using PCA as the dimensionality reduction method.

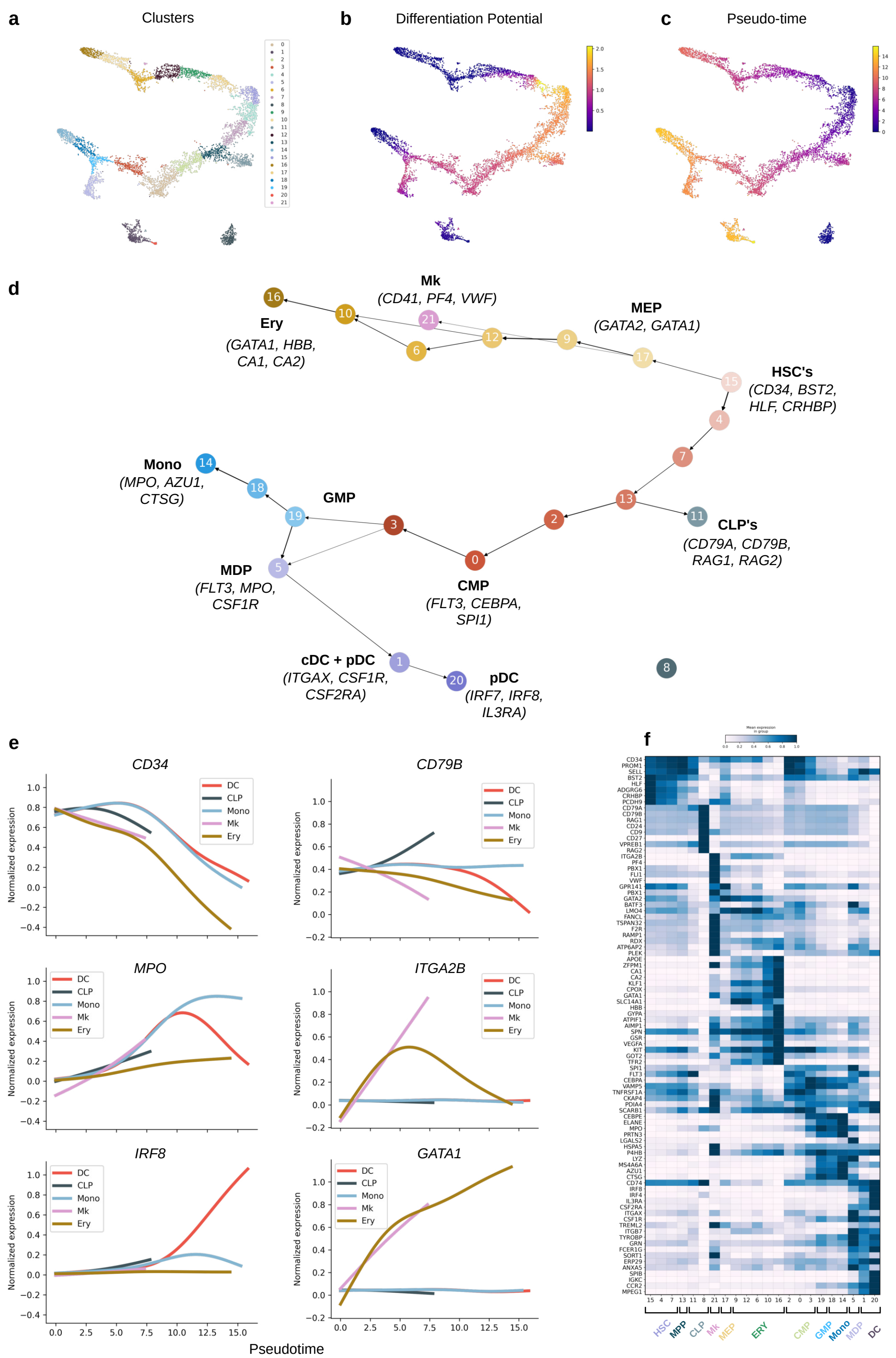

Supplementary Figure 8: **MARGARET results are consistent across replicates. Analysis of scRNA-seq data from replicate 2 of the human hematopoiesis dataset by MARGARET** (a) tSNE plot of cell-state embedding inferred by MARGARET, cells are colored by MARGARET inferred clusters. (b) MARGARET inferred differentiation potential and (c) pseudo-time for each cell projected on the cell embeddings. (d) MARGARET inferred trajectory annotated with cell-type and lineage information (Important marker genes are mentioned within parantheses with the cell-type information). Ery: Erythrocyte; Mk: Megakaryocyte; MEP: Megakaryocyte-Erythroid Progenitors; HSC: Hematopoeitic Stem Cells; CLP: Common Lymphoid Progenitors; CMP: Common Myeloid Progenitors; GMP: Granulocyte-Monocyte Progenitors; MDP: Monocyte-Dendritic Cell Progenitors; cDC: Classical Dendritic Cells; pDC: Plasmacytoid Dendritic Cells; Mono: Monocytes. (e) Gene expression trends for essential genes for major inferred lineages. (f) Heat map for marker genes for all MARGARET inferred clusters for replicate 2.

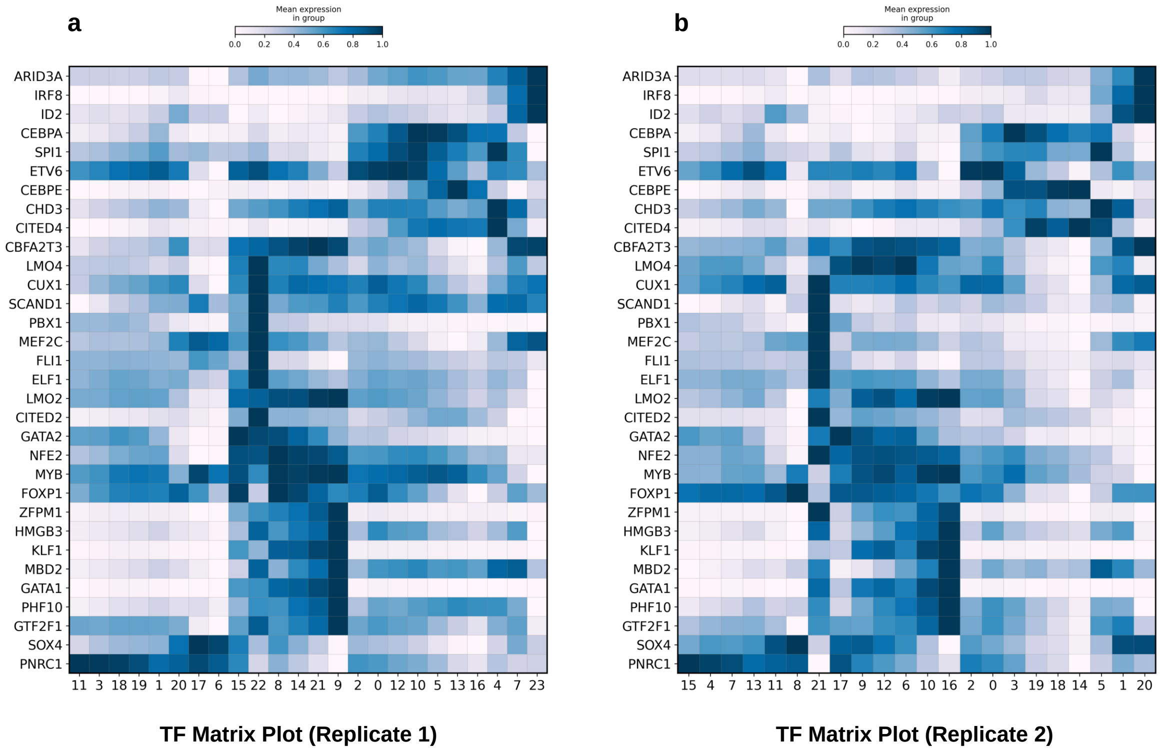

Supplementary Figure 9: **Heat map for transcription factors (TFs) for MARGARET generated clusters across replicates in the scRNA-Hematopoiesis dataset.** (a) Transcription Factor (TF) Matrix Plot for Replicate 1. (b) Transcription Factor (TF) Matrix Plot for Replicate 2.

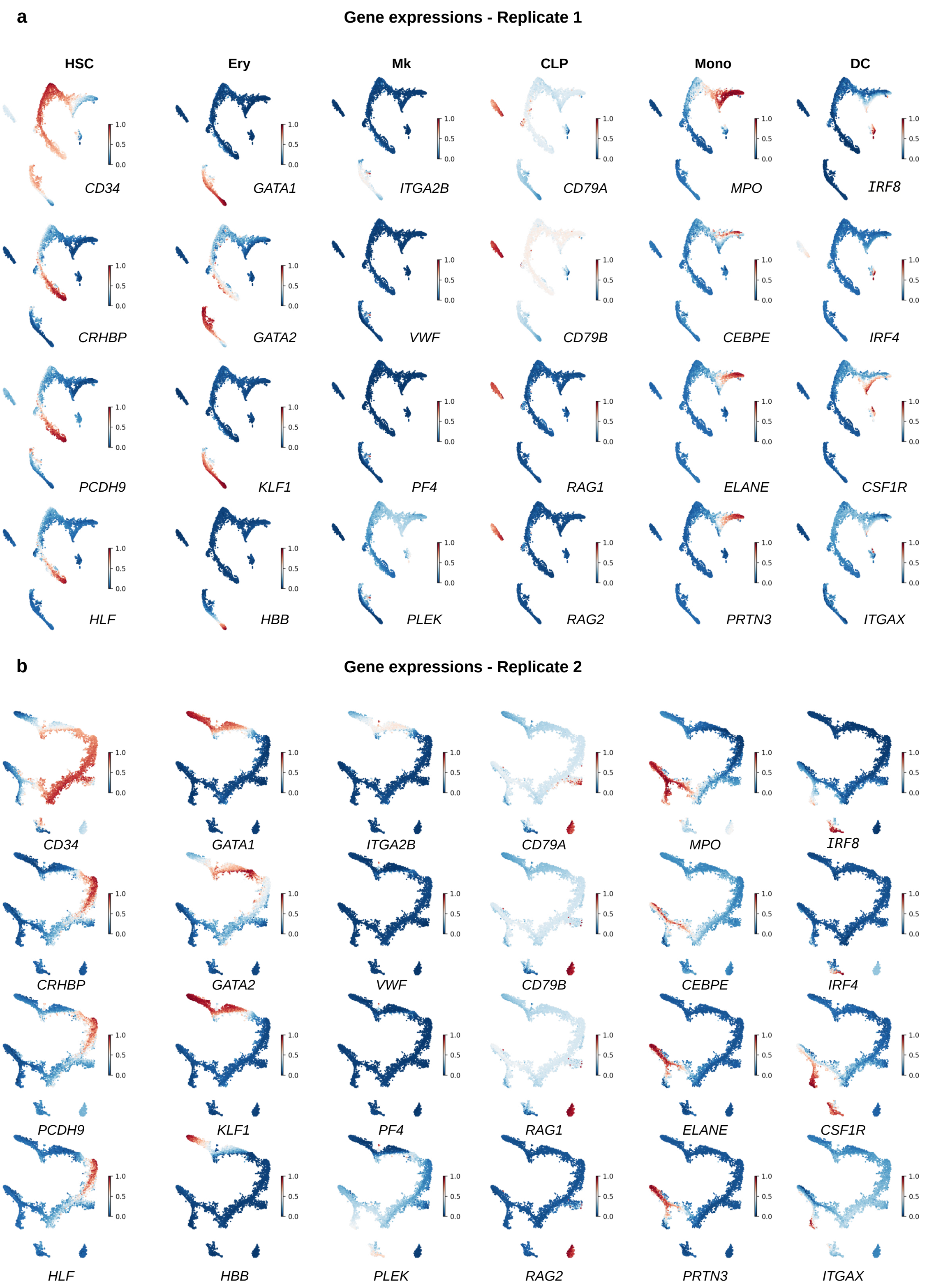

Supplementary Figure 10: **MARGARET captures the underlying global topology of cells for the scRNA-Hematopoiesis dataset across multiple replicates.** (a) Projection of marker gene expressions on the 2-d cell embeddings for Replicate 1 across major lineages inferred using MARGARET. HSC: Hematopoietic Stem Cell; Ery: Erythroid lineage; Mk: Megakaryocyte Lineage; CLP: Common Lymphoid Progenitors; Mono: Monocytes; DC: Dendritic Cell. (b) Same as (a) with the gene expressions projected on 2-d cell embeddings for Replicate 2.

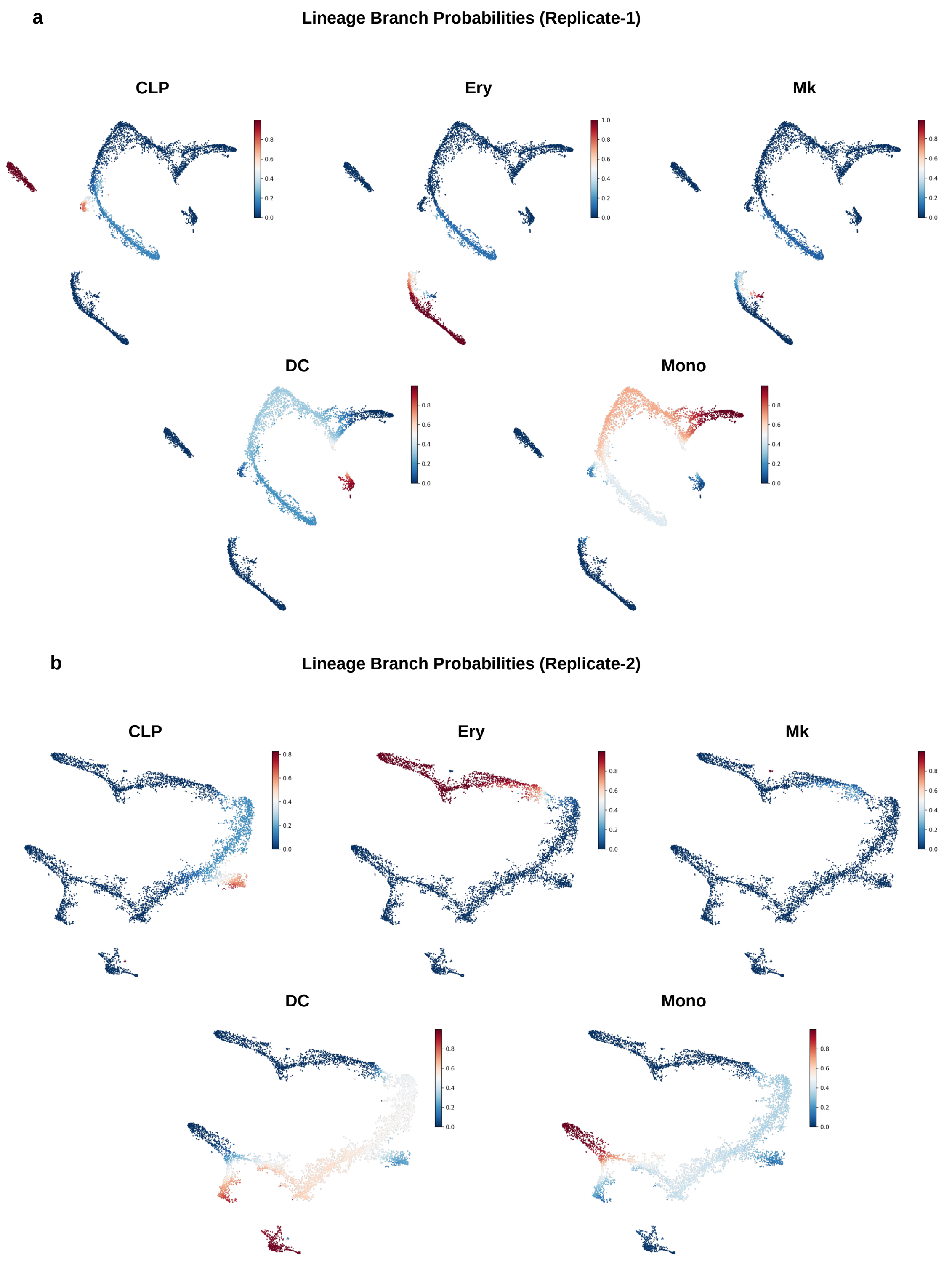

Supplementary Figure 11: **Visualization of lineage specific branch probabilities using MARGARET across replicates in human hematopoiesis data** (a) Cell branch probabilities projected on the 2-d cell embedding space of Replicate 1 for different lineages. CLP: Common Lymphoid Progenitors; Ery: Erythroid lineage; Mk: Megakaryocyte Lineage; Mono: Monocytes; DC: Dendritic Cell. (b) Same as (a) with cell branch probabilities projected on the 2-d cell embedding space of Replicate 2.

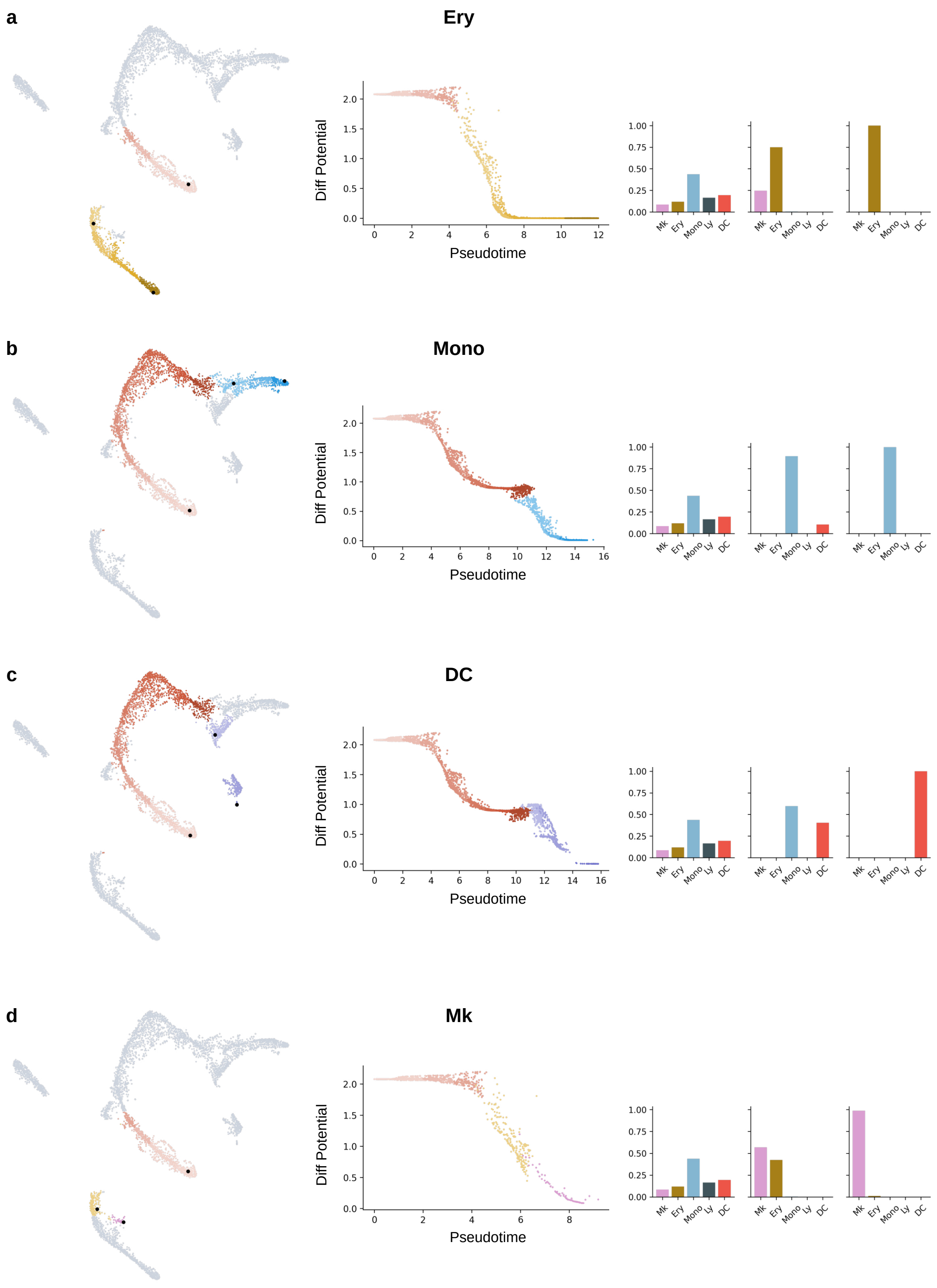

Supplementary Figure 12: **Visualization of lineage specific branch probabilities using MARGARET for replicate 1 in the scRNA-Hematopoiesis dataset** (a)(Left) Erythroid lineage highlighted on the 2-d cell-embedding. Black dots (in bold) represent candidate cells sampled for branch probability visualizations. (Middle) Variation of Differentiation Potential vs Pseudotime for each cell in the erythroid lineage. Each dot is a cell, color coded by their respective cluster in the lineage map (Left). (Right) Branch probabilities for the highlighted cells in the (Left) subfigure for every lineage. (b,c,d) Same as (a) but for the Monocyte (Mono), Dendritic Cell (DC) and Megakaryocyte (Mk) lineages respectively.

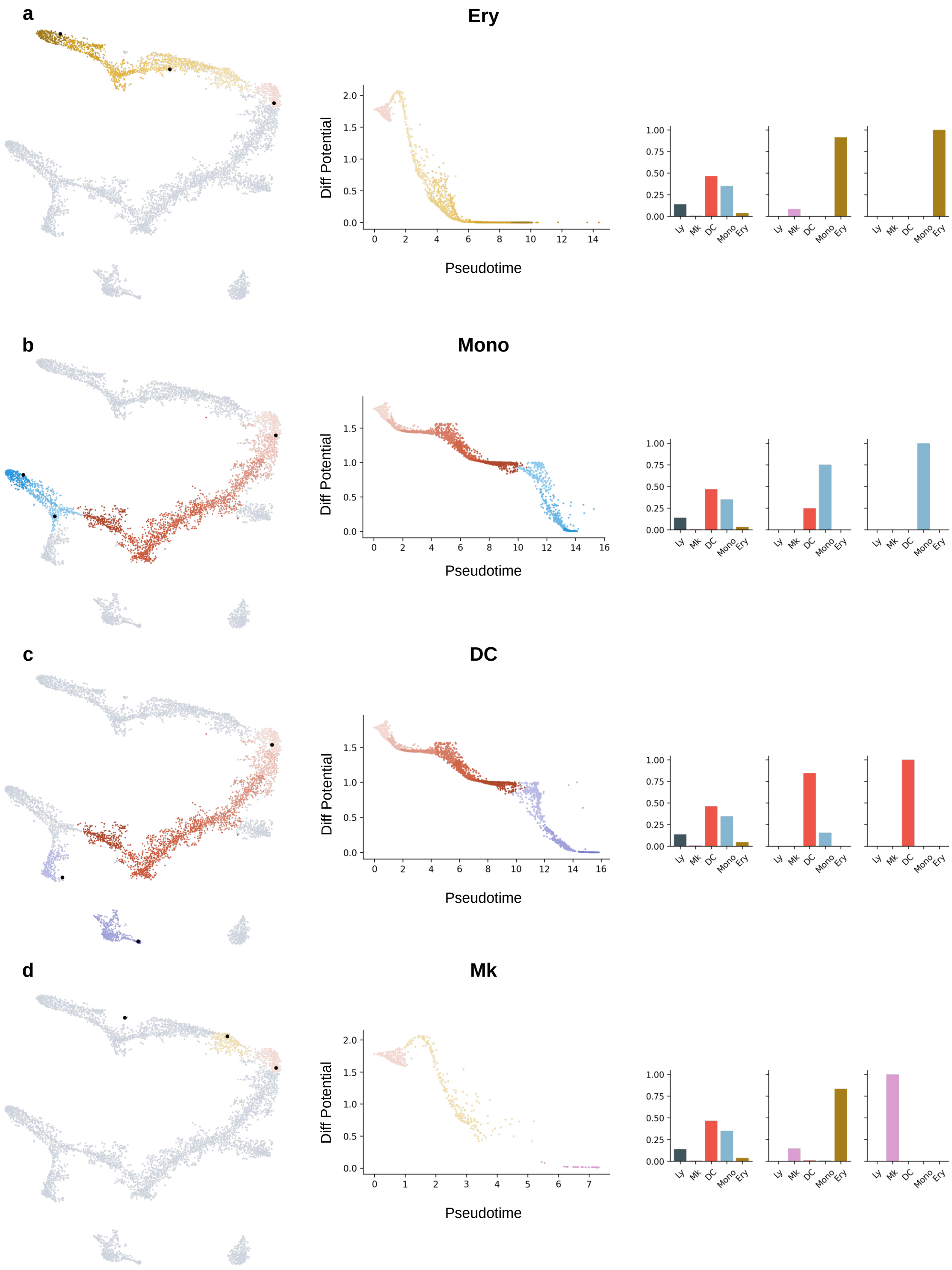

Supplementary Figure 13: **Visualization of lineage specific branch probabilities using MARGARET for replicate 2 in the scRNA-Hematopoiesis dataset** (a)(Left) Erythroid lineage projected on the 2-d cell-embedding visualization. Black dots (in bold) represent candidate cells sampled for Branch probability visualizations. (Middle) Variation of Differentiation Potential vs Pseudotime for each cell in the erythroid lineage. Each dot is a cell, color coded by their respective cluster in the lineage map (Left). (Right) Branch probabilities for the highlighted cells in the (Left) subfigure for every lineage. (b,c,d) Same as (a) but for the Monocyte (Mono), Dendritic Cell (DC) and Megakaryocyte (Mk) lineages respectively.

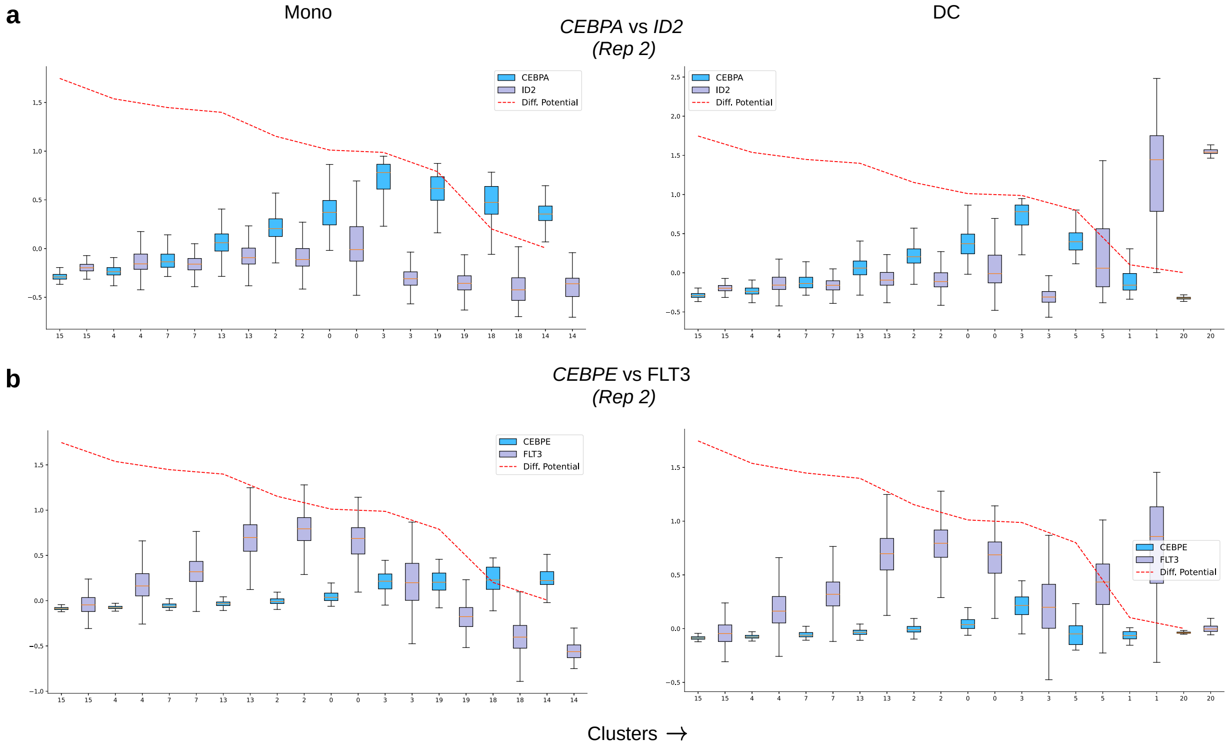

Supplementary Figure 14: **Changes in Differentiation Potential computed using MARGARET correlate with important branching events in the myeloid lineage across replicates in the scRNA-Hematopoiesis data** (a) (Left)Variation of the gene expression of *CEBPA* and *ID2* across the monocyte (Mono) lineage for Replicate 2. (Right) Same as the (Left) subfigure but visualized along the dendritic cell (DC) lineage. (b) Same as (a) but using genes *CEBPE* and *FLT3*. The boxplots summarize the expression of the gene in each cluster in the lineage, where the box depicts the interquartile range (IQR, the range between the 25th and 75th percentile) with the median value, whiskers indicate the maximum and minimum value within 1.5 times the IQR. The red dotted line represents the mean differentiation potential for each cluster in the lineage.

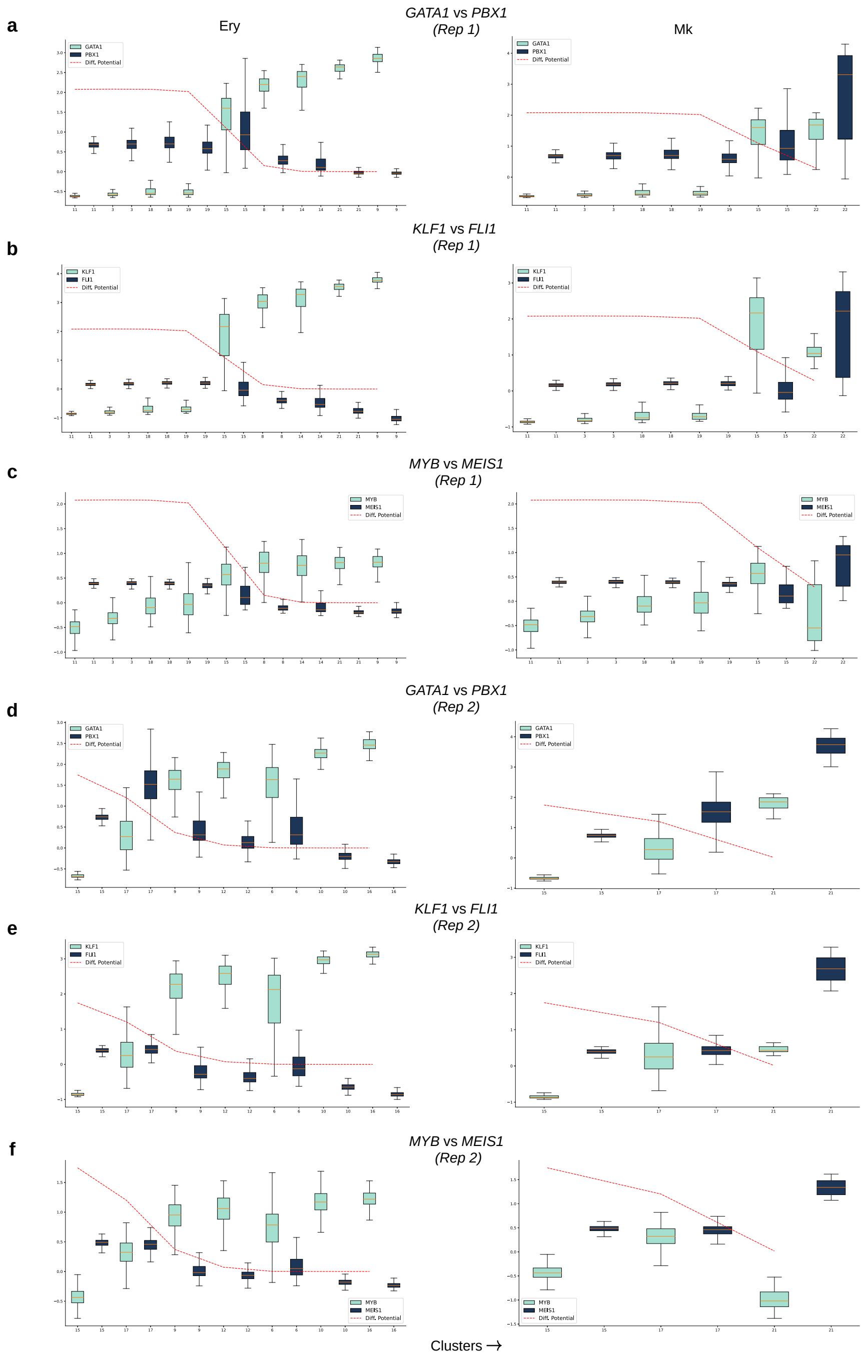

Supplementary Figure 15: **Changes in Differentiation Potential computed using MARGARET correlate with important branching events in the Megakaryocyte-Erythroid lineage across replicates in the scRNA-Hematopoiesis data** (a) (Left)Variation of the gene expression of *GATA1* and *PBX1* across the Erythroid (Ery) lineage for Replicate 1. (Right) Same as the (Left) subfigure but visualized along the megakaryocyte (Mk) lineage. (b, c) Same as (a) but using genes (*KLF1*, *FLI1*) and (*MYB*, *MEIS1*) respectively. (d, e, f) Same as (a, b, c) respectively but for Replicate 2. The boxplots summarize the expression of the gene in each cluster in the lineage, where the box depicts the interquartile range (IQR, the range between the 25th and 75th percentile) with the median value, whiskers indicate the maximum and minimum value within 1.5 times the IQR. The red dotted line represents the mean differentiation potential for each cluster in the lineage.

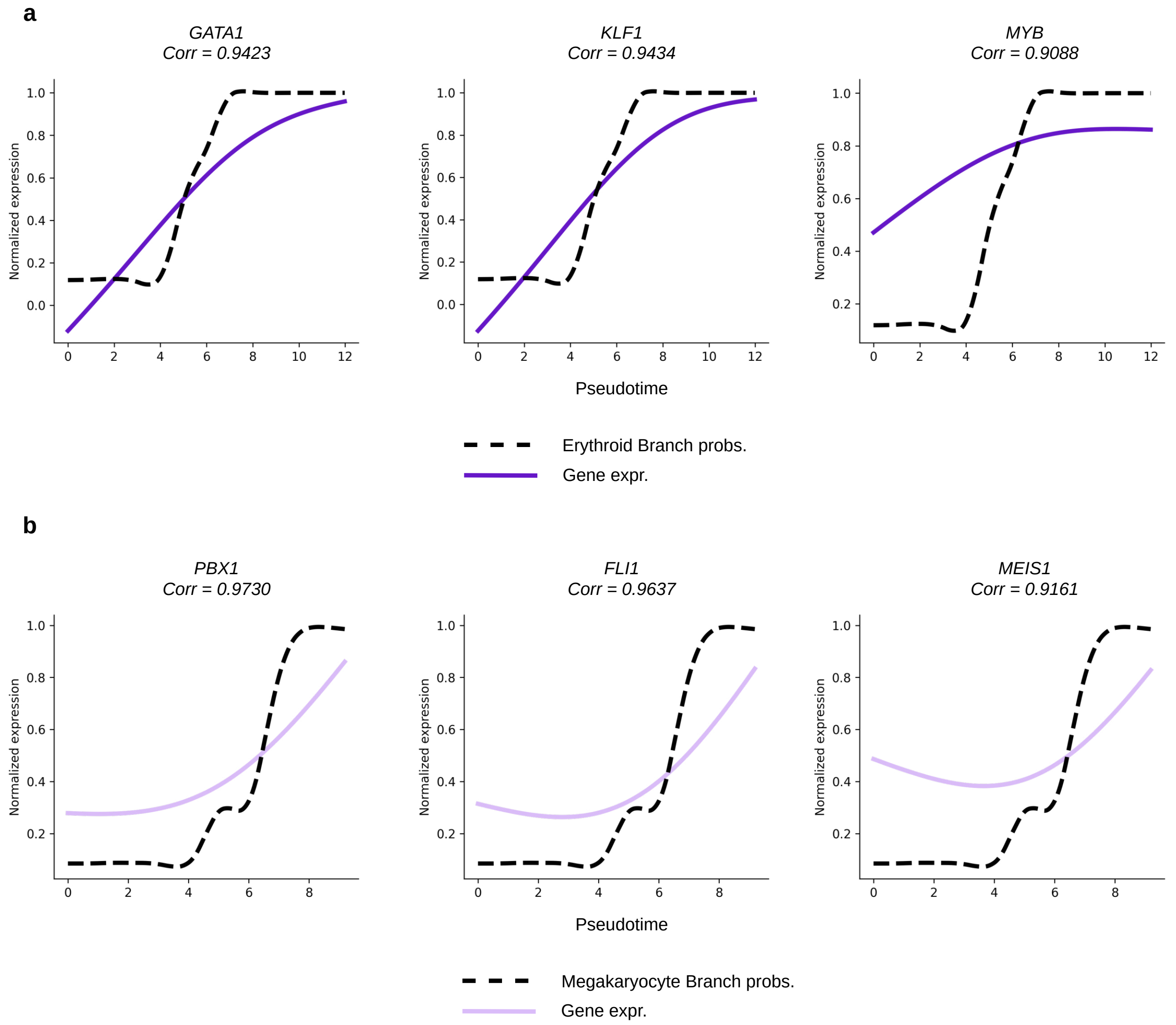

Supplementary Figure 16: **MARGARET inferred erythroid and megakaryocytic branch probabilities correlate with expression of important lineage markers in scRNA human hematopoiesis** (a) Variation of the erythroid branch probabilities and gene expression of erythroid marker genes: *GATA1*, *KLF1*, *MYB*. Correlation between the branch probability and gene expressions trends is shown in the caption for each gene. The plot of branch probabilities vs pseudotime was obtained by fitting a GAM (See methods) on erythroid branch probabilities. (b) Same as (a) but in the megakaryocytic lineage. Correlations were computed for the genes *PBX1, FLI1, MEIS1*.

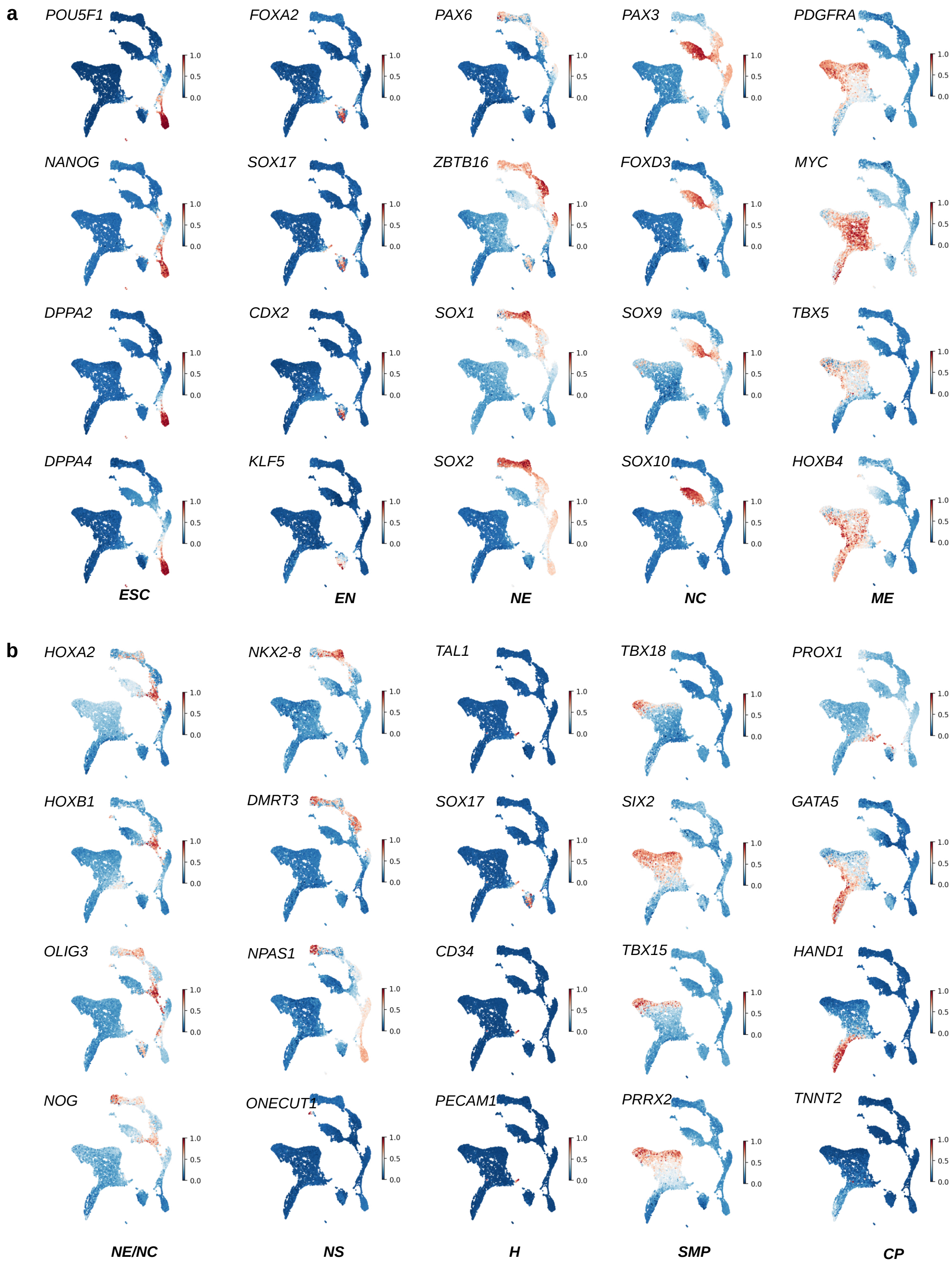

Supplementary Figure 17: **MARGARET captures the underlying global topology of cells for the Embryoid Body dataset.** (a, b) Projection of marker gene expressions on the 2-d cell embeddings across major cell types inferred using MARGARET. ESC: Embryonic Stem Cell; EN: Endoderm; NE: Neuroectoderm; NC: Neural Crest; ME: Mesoderm; NS: Neuronal Subtypes; H: Hemangioblasts; SMP: Smooth Muscle Precursor; CP: Cardiac Progenitors

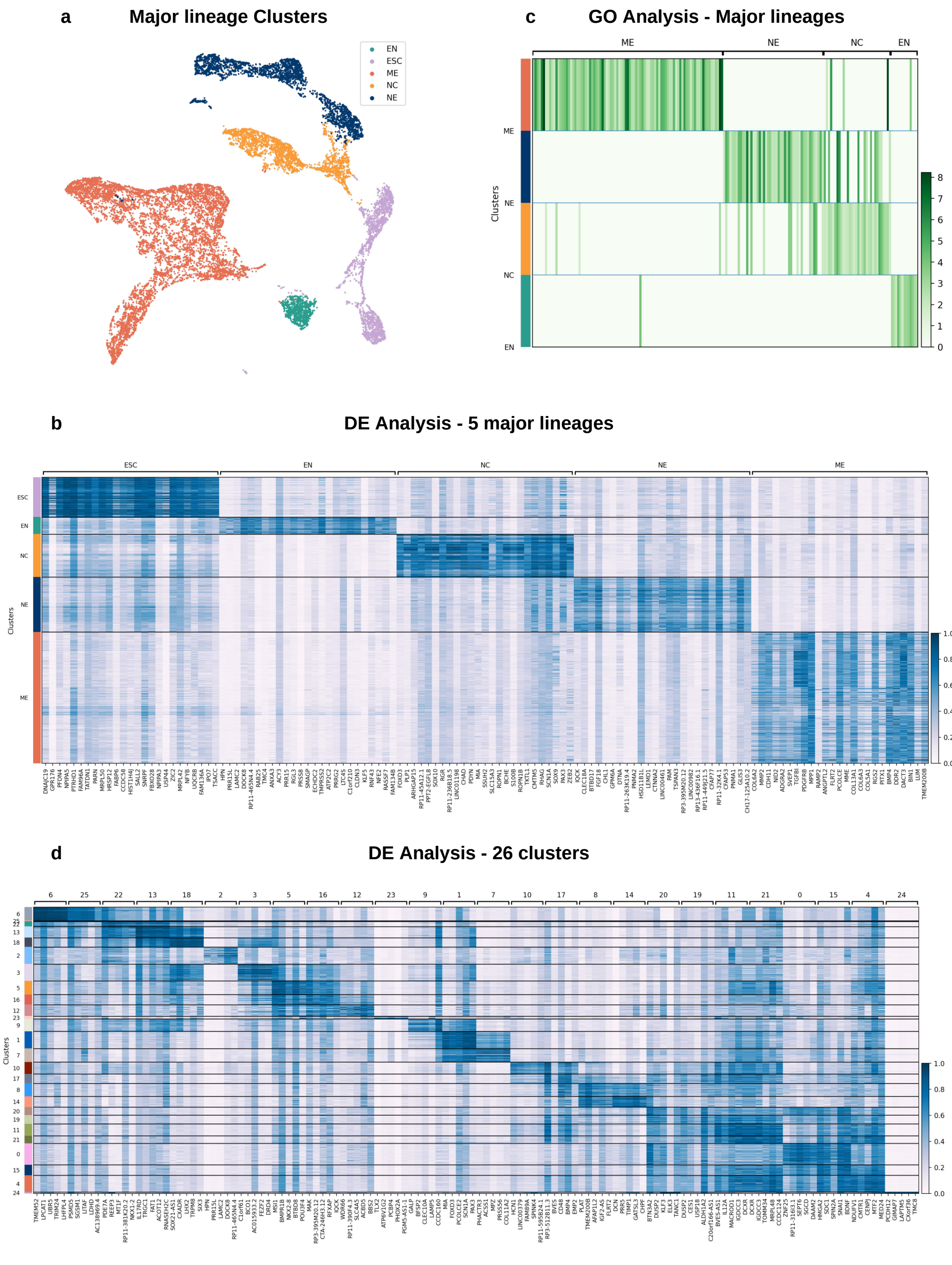

Supplementary Figure 18: **MARGARET accurately delineates major lineages in the Embryoid Body Dataset.** (a) MARGARET cell-state embedding depicting the major lineages in the embryoid body dataset. (b) Differential expression (DE) analysis for the major lineages in the EB Dataset (See (a)). (c) Heatmap of the $p$-values for gene ontology (GO) terms based on the DE genes obtained from (b) for major lineages in the embryoid body dataset (excluding the ESCs). The heatmap value for a GO term was set to $\sqrt{-\log p_{val}}$, where $p_{val}$ is the $p$-value for the corresponding GO term. (d) DE Analysis for all 26 clusters in the embryoid body dataset.

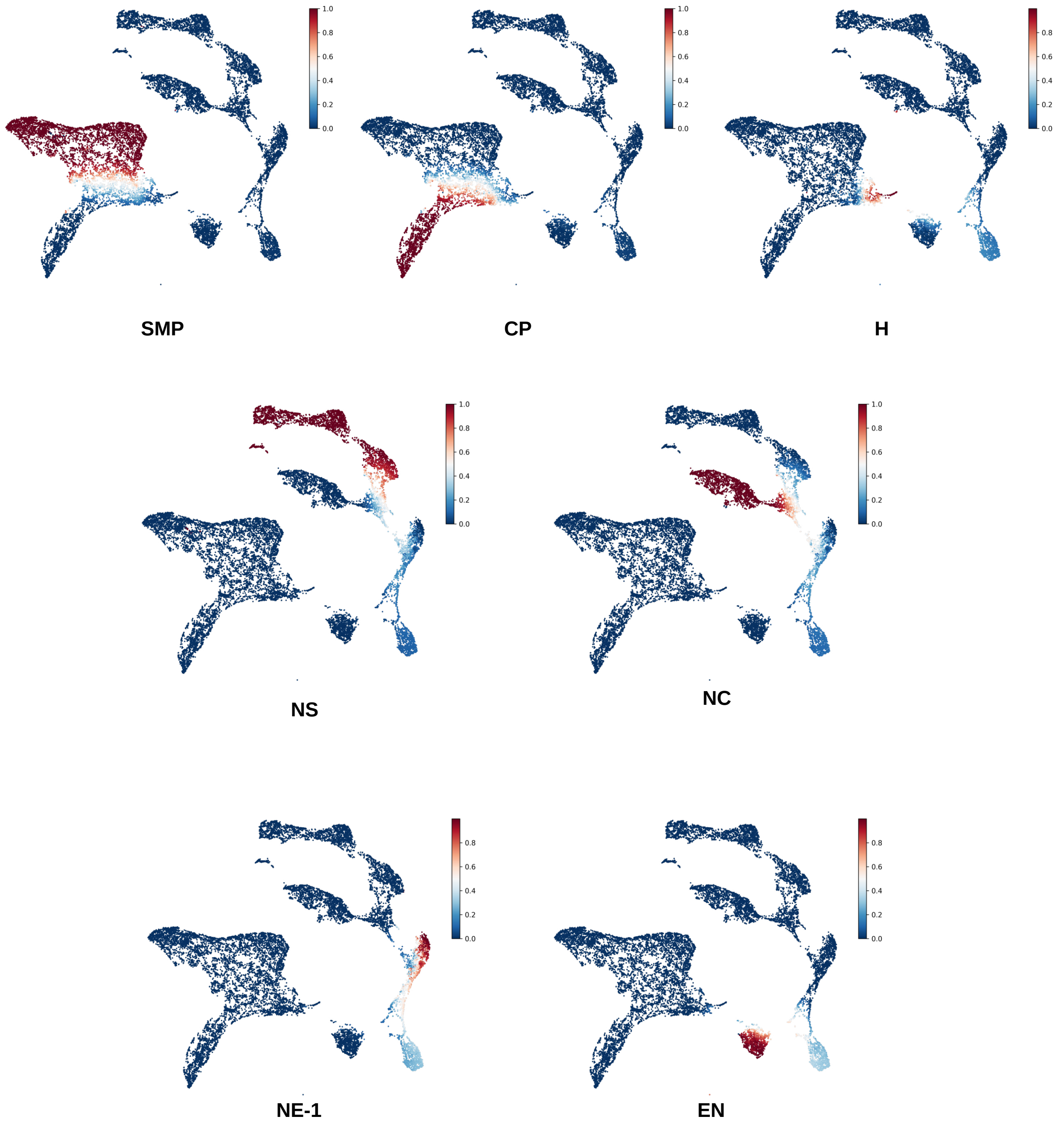

Supplementary Figure 19: **Visualization of lineage specific branch probabilities using MARGARET in the embryoid body dataset.** Cell branch probabilities for the different terminal states were projected on the 2-d cell embedding space for different lineages. SMP: Smooth Muscle Precursor; CP: Cardiac Progenitors; H: Hemangioblasts; NS: Neuronal Subtypes; NC: Neural Crest; NE-1: Neuroectoderm-1; EN: Endoderm

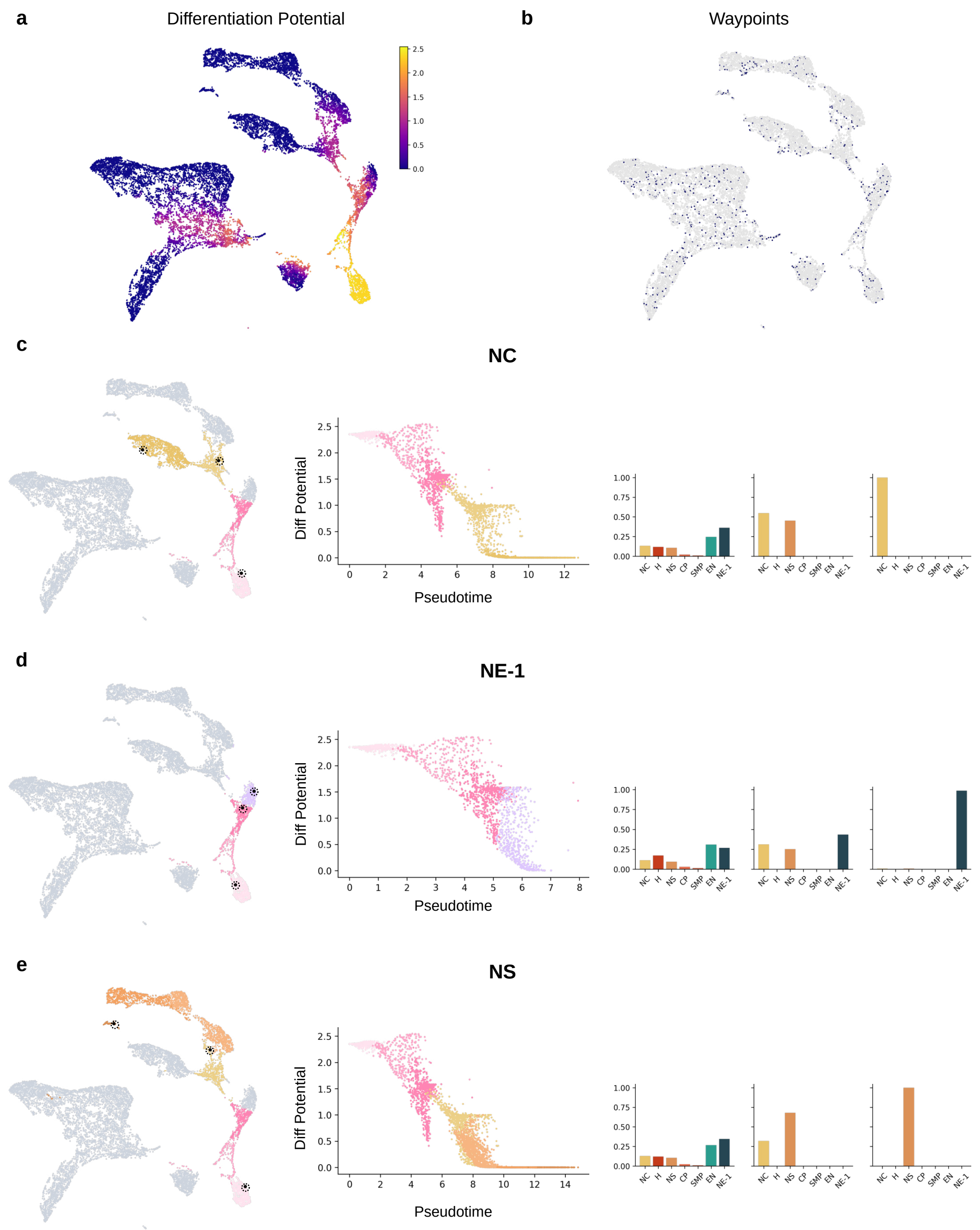

Supplementary Figure 20: **Differential potential analysis for the Neural Crest (NC), Neuroectoderm-1 (NE-1) and Neuronal Subtype (NS) lineages in the embryoid body dataset using MARGARET.** (a) UMAP plot of cells where cells are colored by their differentiation potential as computed by MARGARET. (b) Waypoints sampled during differentiation potential computation (30 waypoints per cluster) (c)(Left) Cells in the neural crest (NC) lineage highlighted on the 2-d cell-embedding. Black dots (in bold) represent candidate cells sampled for branch probability visualizations. (Middle) Variation of differentiation potential (y-axis) as a function pseudotime (x-axis) for the NC lineage. Each dot is a cell, color coded by their respective cluster in the lineage map (Left). (Right) Branch probabilities for the highlighted cells in the (Left) subfigure for every lineage. (d,e) Same as (c) but for the Neuroectoderm-1 (NE-1) and Neuronal Subtype (NS) lineages respectively.

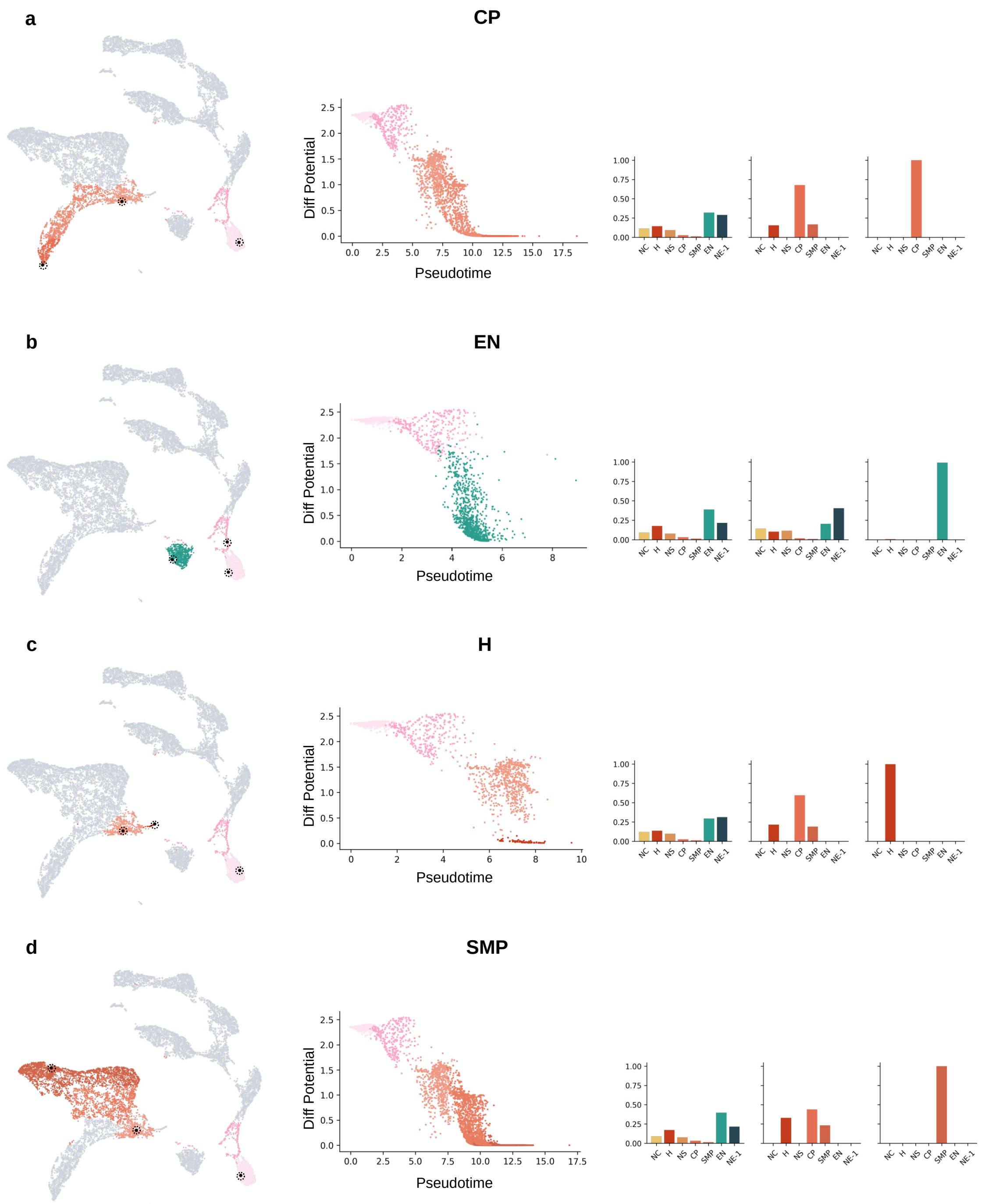

Supplementary Figure 21: **Differential potential analysis for the Cardiac Progenitors (CP), Endoderm (EN), Hemangioblast (H) and Smooth Muscle Precursors (SMP) lineages in the Embryoid Body dataset using MARGARET.** (a)(Left) Cells in the cardiac precursor (CP) lineage highlighted on the 2-d cell-embedding. Black dots (in bold) represent candidate cells sampled for branch probability visualizations. (Middle) Variation of differentiation potential (y-axis) as a function pseudotime (x-axis) for the CP lineage. Each dot is a cell, color coded by their respective cluster in the lineage map in (Left). (Right) Branch probabilities for the highlighted cells in the (Left) subfigure for every lineage. (b,c,d) Same as (a) but for the Endoderm (EN), Hemangioblast (H) and Smooth Muscle Precursors (SMP) lineages respectively.

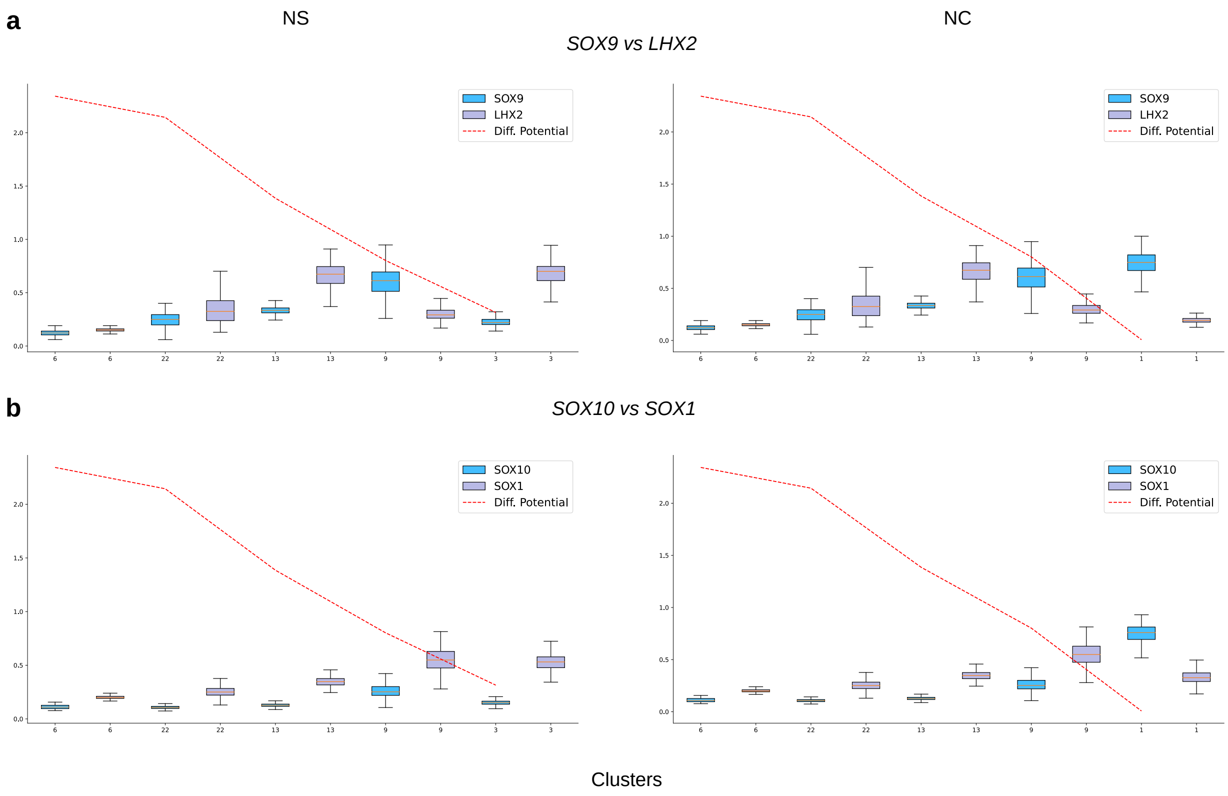

Supplementary Figure 22: **Changes in differentiation potential computed using MARGARET correlate with important branching events in the ectoderm lineage in the embryoid body dataset.** (a) (Left) Variation of the expression of transcription factors (TFs) *SOX9* and *LHX2* across the neuronal subtype (NS) lineage. The red dotted line represents the mean differentiation potential for each cluster. (Right) Same as the (Left) subfigure but visualized along the neuronal crest (NC) lineage. (b) Same as (a) but using TFs *SOX10* and *SOX1*.

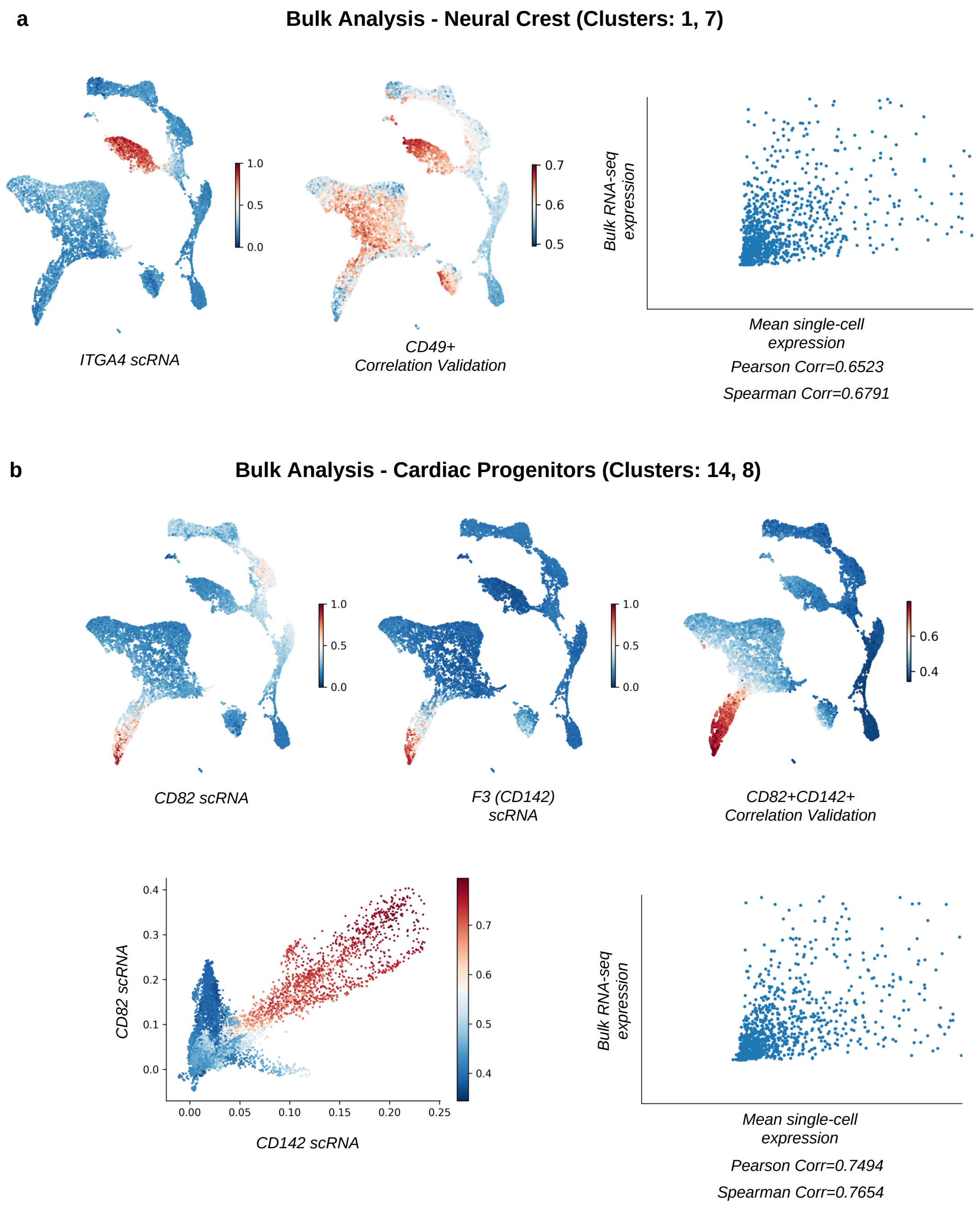

Supplementary Figure 23: **Bulk analysis validates the Neural Crest (NC) and Cardiac Progenitor (CP) lineages detected by MARGARET.** (a) (Left): *ITGA4* gene expression projected on the 2-d cell embeddings. (Middle): Correlation between CD49+ sorted Bulk RNA-seq transcription factor expression and the scRNA-seq expressions of each cell projected on the 2-d cell embeddings. (Right): Scatter plot of Bulk RNA-seq expression vs the mean scRNA-seq expression (for cells in clusters 1, 7). (b) (Top-Left): *CD82* gene expression projected on the 2-d cell embeddings. (Top-Middle): *CD142* gene expression projected on the 2-d cell embeddings. (Top-Right): Correlation between $CD82^{+}CD142^{+}$ sorted Bulk RNA-seq transcription factor expression and the scRNA-seq expressions of each cell projected on the 2-d cell embeddings. (Bottom-Left): Scatter plot of *CD82* vs *CD142* scRNA-seq expression values for the Embryoid Body dataset. Each cell is colored by the correlation obtained in (Top-Right) figure. (Bottom-Right): Scatter plot of Bulk RNA-seq expression vs the mean scRNA-seq expression (for cells in clusters 14,8).

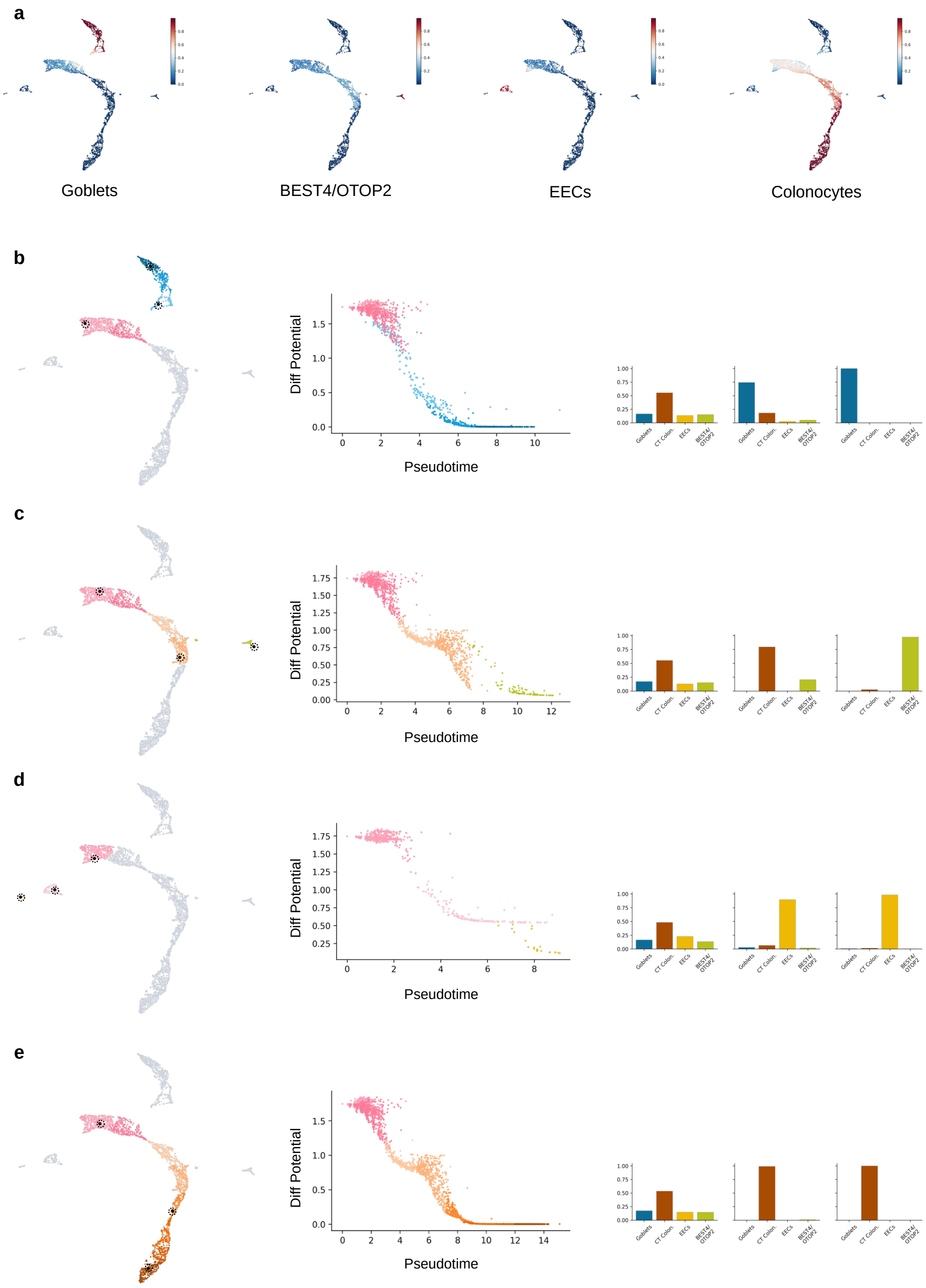

Supplementary Figure 24: **Visualization of lineage specific branch probabilities using MARGARET in the Colon IBD dataset under normal conditions.** (a) Cell branch probabilities projected on the 2-d cell embedding space for different lineages. (b) (Left) Goblet cell lineage projected on the 2-d cell-embedding visualization. Black dots (in bold) represent candidate cells sampled for Branch probability visualizations. (Middle) Variation of differentiation potential vs pseudotime for each cell in the goblet lineage. Cells are colored by the color of their respective cluster in the lineage map in (Left). (Right) Branch probabilities for the highlighted cells in the (Left) subfigure for every lineage. (c, d, e) Same as (b) but for BEST4/OTOP2, Enteroendocrine cells (EEC's) and CT colonocyte lineages, respectively.

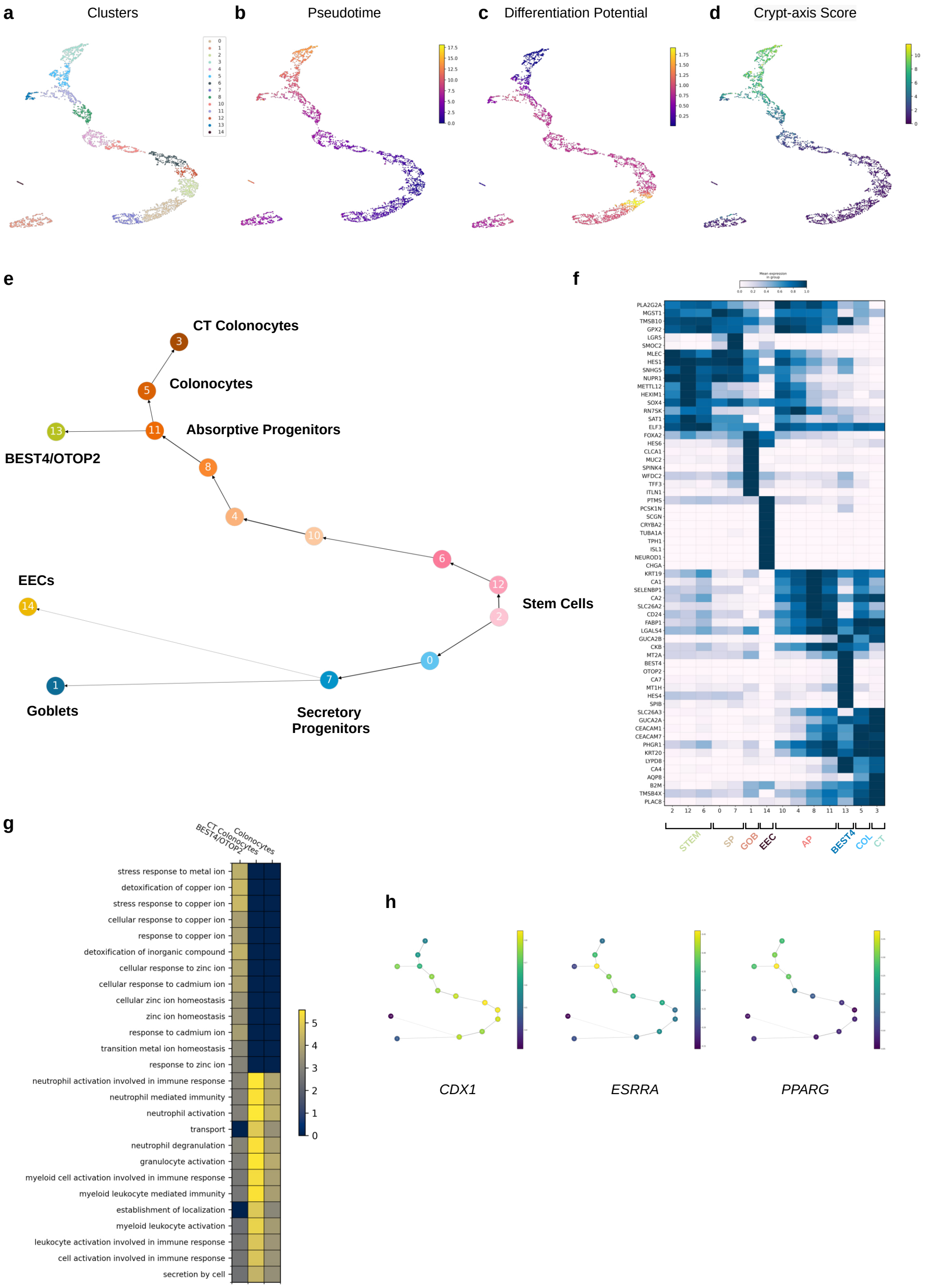

Supplementary Figure 25: **MARGARET applied to the Colon IBD dataset under non-inflamed conditions.** (a) MARGARET inferred clusters. (b) Inferred Pseudotime (c) Differentiation potential projections on the inferred 2-d cell embeddings. (d) CA score projected on the 2-d cell embeddings (e) MARGARET inferred trajectory for non-inflamed cells showcasing the branching of BEST4/OTOP2 cells from absorptive progenitors. (f) Matrix Plot for the UC non-inflamed cells. (g) GO analysis for characterizing the functional significance of BEST4/OTOP2 cells, colonocytes and CT colonocytes in the absorptive lineage for the UC non-inflamed case. The heatmap value for a GO term was set to $\sqrt{-\log p_{val}}$, where $p_{val}$ is the p-value for the corresponding GO term. (h) Mean expression of *CDX1*, *ESRRA* and *PPARG*, projected on the MARGARET inferred connectivity graph for non-inflamed colon.

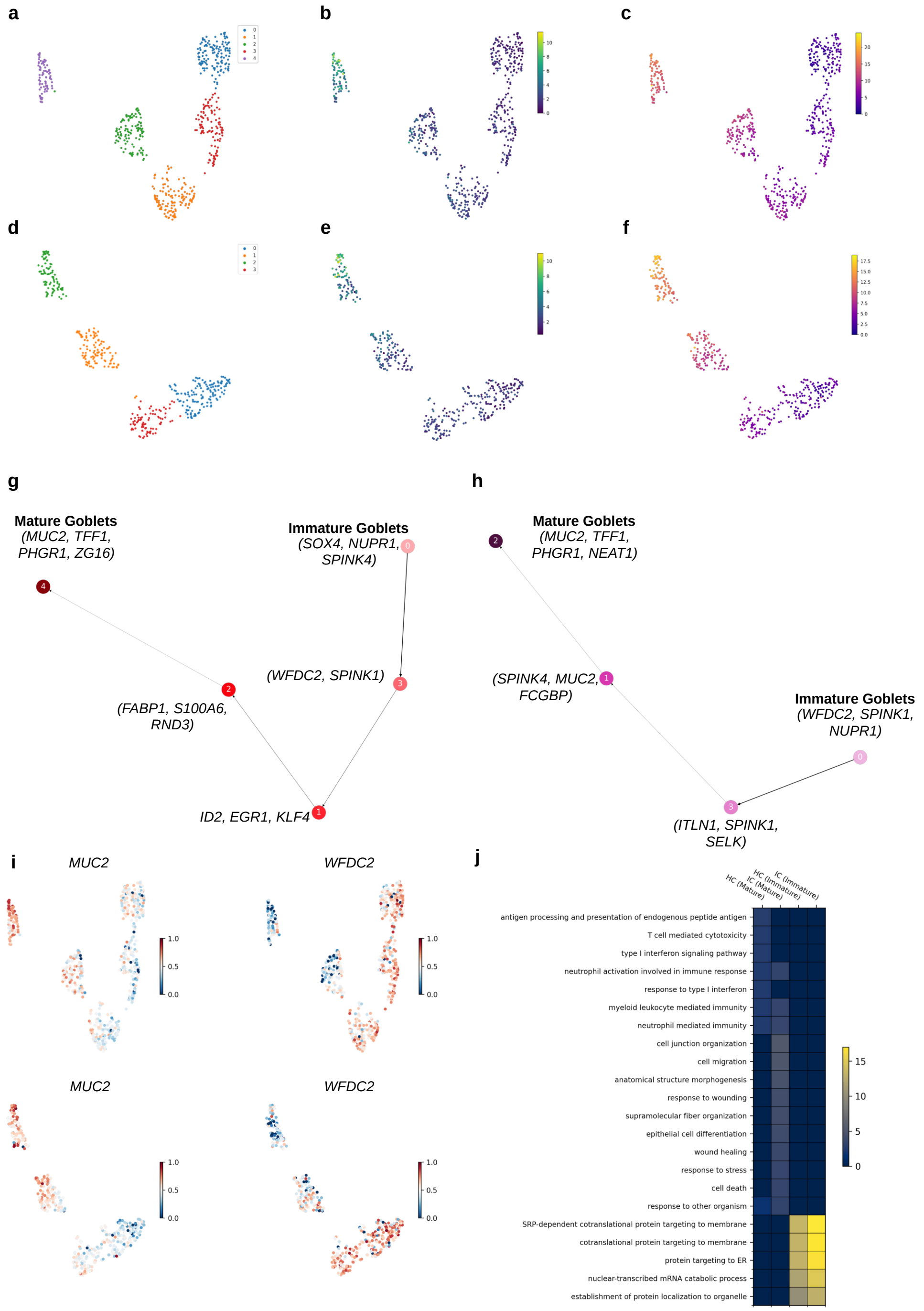

Supplementary Figure 26: **Analysis of the goblet cell lineage under normal and inflamed conditions using MARGARET.** (a, b, c) Clusters, crypt-axis score projections and pseudotime projections on the 2-d goblet cell embeddings, respectively for the normal colon. (d, e, f) Clusters, crypt-axis score projections and pseudotime projections on the 2-d goblet cell embeddings, respectively for the inflamed colon. (g) MARGARET inferred trajectory for the goblet cell lineage in normal colon. (h) Same as (g) but for the inflamed colon. (i) Visualization of the expression of important marker genes: *WFDC2* and *MUC2* in the normal colon (Top) and the inflamed colon (Bottom) (j) GO analysis between mature and immature goblet cells under normal and inflamed conditions. The heatmap value for a GO term was set to $\sqrt{-\log p_{val}}$, where $p_{val}$ is the p-value for the corresponding GO term

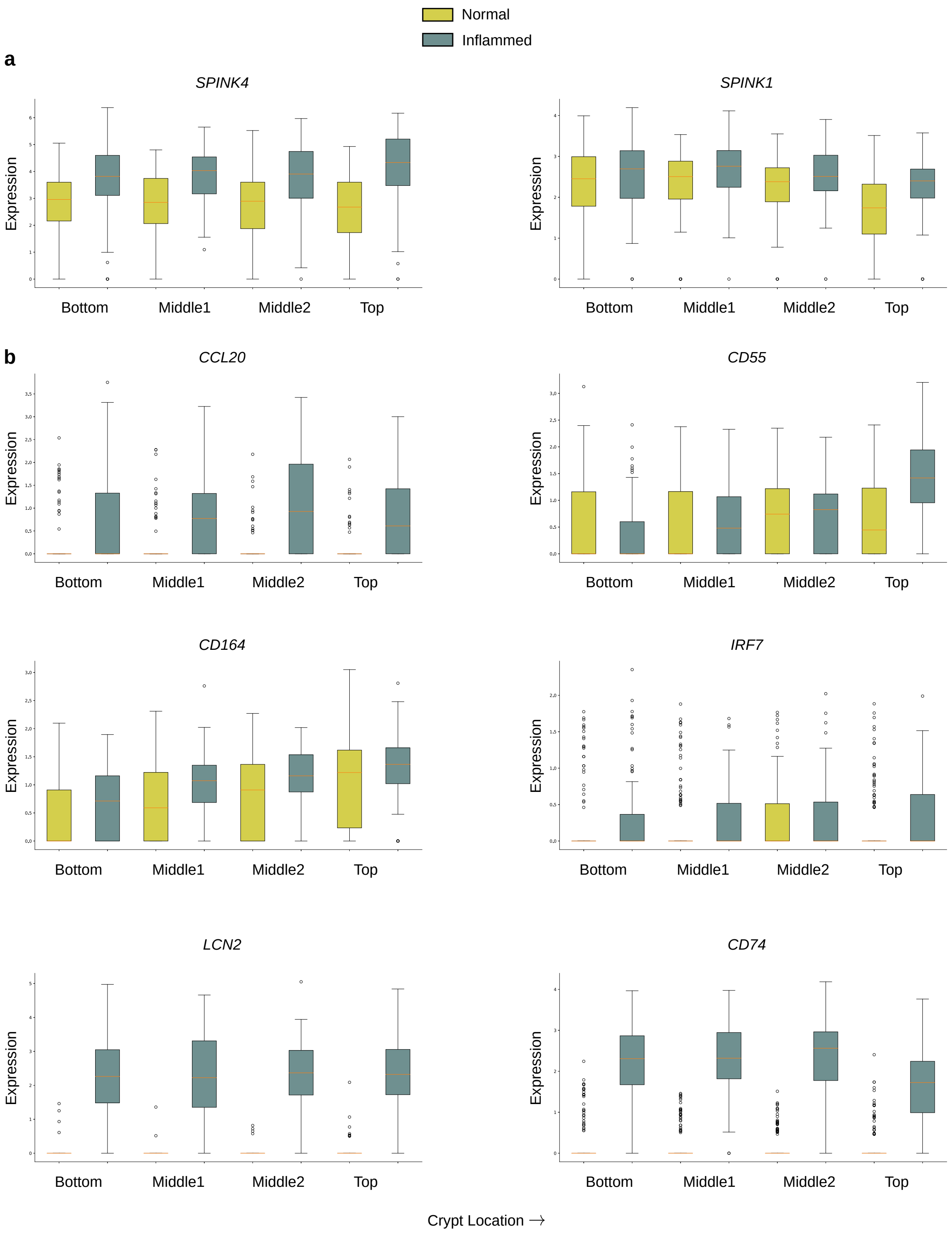

Supplementary Figure 27: **Dysregulation of marker genes and cytokines in MARGARET inferred goblet cell lineages under UC-inflamed conditions.** (a) Dysregulation in early goblet cell markers (b) Dysregulation in the expression of cytokines under inflamed conditions. The boxplots of expressions were obtained by binning the cells in the goblet lineage into four quartile groups on the basis of their inferred crypt-axis scores. The labels on the x-axis denote the corresponding crypt-axis locations. The boxplots summarize the expression of the gene in the binned crypt-locations, where the box depicts the interquartile range (IQR, the range between the 25th and 75th percentile) with the median value, whiskers indicate the maximum and minimum value within 1.5 times the IQR.

### Supplementary Tables

| Method | Prior Information | Global Topology | Pseudotime Ordering | Multiple Start cells | Terminal State Detection | Cell Branch Probabilities | Differentiation Potential |
|---|---|---|---|---|---|---|---|
| MARGARET | Start cell | ▲ | ▲ | ▲ | ▲ | ▲ | ▲ |
| Monocle 3 | Start cell | ▲ | ▲ | ▲ | ▼ | ▼ | ▼ |
| PAGA | Start cell | ▲ | ▲ | ▼ | ▼ | ▼ | ▼ |
| Palantir | Start cell | ▼ | ▲ | ▼ | ▲ | ▲ | ▲ |
| DPT | Start cell | ▼ | ▲ | ▼ | ▼ | ▼ | ▼ |

▲ Supported

▼ Not-supported

**Supplementary Table 1: Qualitative Comparison between MARGARET and other TI methods in terms of supported inputs-outputs** Column explanations: **Prior Information** User-defined input to the method; **Global Topology** Whether the method captures all types of trajectory topologies; **Pseudotime Ordering** Whether the method orders the cells along the inferred trajectory; **Multiple Start Cells** Whether the method allows specifying multiple start cells (useful for modelling disconnected processes); **Terminal State Detection** Whether the method allows automatic Terminal state prediction; **Cell Branch Probabilities** Whether the method computes the probability of branching into a terminal state at the cell-level; **Differentiation Potential** Whether the method allows computation of Cell-differentiation potential for each cell.

| Dataset | MARGARET (Louvain@1.0) | MARGARET (Leiden @1.0) | PAGA | Palantir | Monocle3 (Leiden) | Monocle3 (Louvain) |
|---|---|---|---|---|---|---|
| Multifurcating_1 | **0.937** | 0.922 | 0.846 | 0.303 | -0.4375 | -0.4354 |
| Multifurcating_2 | 0.905 | **0.919** | 0.848 | 0.908 | -0.3174 | -0.312 |
| Multifurcating_3 | 0.926 | **0.941** | 0.902 | 0.923 | -0.2726 | -0.2746 |
| Multifurcating_4 | 0.808 | 0.819 | 0.693 | **0.82** | -0.4765 | -0.4665 |
| Multifurcating_5 | 0.936 | 0.938 | 0.899 | **0.958** | 0.4573 | 0.4667 |
| Disconnected_1 | 0.857 | **0.862** | 0.451 | 0.44 | 0.8116 | 0.805 |
| Disconnected_2 | 0.82 | 0.792 | 0.235 | - | 0.8378 | **0.8671** |
| Disconnected_3 | **0.762** | 0.754 | 0.063 | 0.045 | 0.7144 | 0.7338 |
| Disconnected_4 | 0.691 | **0.714** | 0.096 | -0.021 | 0.6097 | 0.2118 |
| Disconnected_5 | 0.809 | **0.864** | 0.101 | 0.161 | 0.4549 | 0.5142 |

(a) KT Correlation Score comparison between MARGARET and other TI methods

| Dataset | MARGARET (Louvain@1.0) | MARGARET (Leiden @1.0) | PAGA | Palantir | Monocle3 (Leiden) | Monocle3 (Louvain) |
|---|---|---|---|---|---|---|
| Multifurcating_1 | **0.993** | 0.991 | 0.963 | 0.466 | -0.7283 | -0.728 |
| Multifurcating_2 | 0.984 | **0.987** | 0.967 | 0.984 | -0.6362 | -0.6354 |
| Multifurcating_3 | 0.989 | **0.994** | 0.984 | 0.989 | -0.6066 | -0.6067 |
| Multifurcating_4 | 0.943 | 0.945 | 0.865 | **0.95** | -0.7092 | -0.7068 |
| Multifurcating_5 | 0.99 | 0.991 | 0.979 | **0.995** | 0.3185 | 0.3209 |
| Disconnected_1 | 0.972 | **0.973** | 0.541 | 0.602 | 0.9523 | 0.9456 |
| Disconnected_2 | 0.944 | 0.923 | 0.234 | - | 0.9576 | **0.9732** |
| Disconnected_3 | **0.929** | 0.927 | 0.095 | 0.089 | 0.8768 | 0.9092 |
| Disconnected_4 | 0.852 | **0.873** | 0.131 | -0.039 | 0.7624 | 0.1609 |
| Disconnected_5 | 0.95 | **0.975** | 0.094 | 0.191 | 0.4273 | 0.4502 |

(b) SR Correlation Score comparison between MARGARET and other TI methods

**Supplementary Table 2: Quantitative comparison of pseudotime ordering between MARGARET, PAGA, Palantir and Monocle 3 using the Kendall's Tau (KT) and Spearman Rank (SR) correlation metrics on our simulated benchmark.** (a) KT-score based comparison. (b) SR-score based comparison. It is worth noting that the KT and the SR metrics are consistent with each other (i.e. a method having a higher KT score also has a higher SR score). The symbol - represents that the method was not stable when evaluating for the dataset. In such a case we used a value of -1.0 (minimum correlation value) in the plot for illustration purposes only.

| Dataset | No. of cells | No. of genes | Trajectory type |
|---|---|---|---|
| Disconnected_1 | 1966 | 501 | Disconnected (2) |
| Disconnected_2 | 4929 | 3795 | Disconnected (3) |
| Disconnected_3 | 4945 | 395 | Disconnected (4) |
| Disconnected_4 | 5000 | 2500 | Disconnected (4) |
| Disconnected_5 | 7500 | 3500 | Disconnected (3) |
| Disconnected_6 | 2500 | 1500 | Disconnected (2) |
| Multifurcating_1 | 4945 | 3870 | Multifurcating |
| Multifurcating_2 | 5000 | 2500 | Multifurcating |
| Multifurcating_3 | 5000 | 1000 | Multifurcating |
| Multifurcating_4 | 4500 | 3000 | Multifurcating |
| Multifurcating_5 | 7500 | 2500 | Multifurcating |
| Cyclic_1 | 8145 | 531 | Cyclic (1) |

(a) Simulated dataset statistics used for MARGARET benchmarking. The numerical value in paranthesis for the datasets indicates the number of disconnected components and the number of cycles in the disconnected and cyclic simulated datasets, respectively.

| Dataset | Modality | Organism | No. of cells | No. of genes | Tissue | Reference |
|---|---|---|---|---|---|---|
| Hematopoiesis (Replicate 1) | scRNA-seq | Human | 5780 | 14651 | Blood (Hematopoiesis) | (1) |
| Hematopoiesis (Replicate 2) | scRNA-seq | Human | 6501 | 14913 | Blood (Hematopoiesis) | (1) |
| Embryoid Body | scRNA-seq | Human | 16821 | 17845 | Embryo (Embryogenesis) | (22) |
| Colon IBD | scRNA-seq | Human | 11175 | 58690 | Colon (IBD) | (28) |
| PBMC 8k | scRNA-seq | Human | 7982 | 3346 | Blood | 10x Genomics |
| PBMC 4k | scRNA-seq | Human | 4008 | 3346 | Blood | 10x Genomics |
| HCA (subsampled) | scRNA-seq | Human | 18641 | 26662 | Heart Cell Atlas | (33) |
| CORTEX | scRNA-seq | Mouse | 3005 | 19972 | Brain | (34) |

(b) Real biological dataset statistics used in this study.

**Supplementary Table 3: Overview of simulated and real biological datasets used in this study.**
